# Comparative analysis of trans-chromosomic rodent models reveals improved somatic hypermutation and class-switch recombination in rats

**DOI:** 10.1101/2024.08.26.609625

**Authors:** Hiroyuki Satofuka, Satoshi Abe, Takashi Moriwaki, Akane Okada, Kanako Kazuki, Shusei Hamamichi, Masaharu Hiratsuka, Masumi Hirabayashi, Kazuomi Nakamura, Tetsushi Sakuma, Takashi Yamamoto, Yoshihiro Baba, Kazuma Tomizuka, Yasuhiro Kazuki

## Abstract

Humanized rodent models, especially humanization of genetic/genomic components involved in immunity have significantly advanced our understanding of human immune system. Here, we utilized trans-chromosomic (Tc) technology to generate a TC-mAb rat model that stably harbors a mouse artificial chromosome carrying full-length human immunoglobulin (Ig) heavy and kappa light chain genes (IGHK-NAC) in a rat Ig knockout background. In contrast with TC-mAb mice, serum human IgG concentration was found higher than IgM. Number of lymphocytes was recovered, and B cell population in the spleen was normal. Remarkably, repertoire analysis revealed similarities between the model and human PBMCs; somatic hypermutation and class-switch recombination also more closely resembled humans. Furthermore, immunization resulted in generation of antigen-specific human antibodies. Collectively, our strategy to generate both rat and mouse models through introduction of the identical IGHK-NAC offers unprecedented opportunities to comprehensively evaluate genomic regulation and its outcomes associated with genomic sequences and host-derived protein factors.

## Introduction

Genomically humanized rodent models wherein their rodent genetic or chromosomal components have been modified to carry human genomic sequences are commonly used to recapitulate human gene expression patterns and to elucidate biological functions associated with human health and diseases^1,2^. To this end, we have generated multiple humanized rat and mouse models termed trans-chromosomic (Tc) animals that stably maintain a series of artificial chromosomes encoding human chromosome fragments or megabase-sized genes, and demonstrated compatible expression and regulation of the human genomes in the rodents^3–6^. Since utilization of the artificial chromosomes allows stable and independent retention of human genome of unlimited size inside the host cells with minimal incidences of overexpression or silencing, these models have been extensively investigated for studying genetic disorders, pharmacological effects, and human antibody (Ab) production.

Previously, we reported a TC-mAb mouse model that stably maintained a mouse artificial chromosome (MAC) containing full-length human immunoglobulin (Ig) heavy (*IGH,* 1.8 Mb from human chromosome 14) and kappa (*IGK,* 1.7 Mb from human chromosome 2) gene loci including all regulatory elements (IGHK-NAC) in a mouse Ig knockout background^7^. In contrast with wildtype ICR mice, TC-mAb mice exhibited: 1) reduced class-switch recombination (CSR) from IgM to IgG, 2) delayed B cell development, 3) enhanced elicitation of plasmablast and plasma cell differentiation after antigen administration, while compared with human samples, our model demonstrated: 4) similar use and recombination of human variable (V), diversity (D), and joining (J) gene segments and 5) reduced somatic hypermutation (SHM) of human Ig genes. These comparative results suggested the alteration of phenotypes associated with introduction of the human genome in the mouse cells or evolutionary differences between mouse and human cellular environment.

To exploit our earlier findings, we herein reasoned that comparative functional genomic approach among multiple TC-mAb animal models and human samples would further clarify the role of evolutionary conserved interconnection between genomes and cellular environment. Since the MACs including IGHK-NAC are stable not only in mice but also in rats^6,8^, we chose rats as an ideal model organism for this study. Additionally, rats have a longer lifespan than mice, which is suitable for immunological research such as autoimmune diseases and grafting experiments^9^, and immunization may result in production of monoclonal Abs (mAbs) that cross-react with human and mouse antigens^10,11^. Furthermore, our strategy is advantageous in a manner that the identical genomic information is maintained in the rat or mouse cellular setting, minimizing the confounding effects that are crucial for interpretation of data.

Here, we generated a TC-mAb rat model that retained the IGHK-NAC in a rat Ig knockout background, and functionally evaluated its immune response. In contrast to TC-mAb mice, we determined that serum human IgG level in the TC-mAb rats was higher than IgM. Number of splenocytes in TC-mAb rats was significantly recovered to the same level as wild-type rats. Human V(D)J recombination was reproduced in TC-mAb rats, and frequencies of CSR and SHM were significantly improved compared to TC-mAb mice. Furthermore, we immunized TC-mAb rats with ovalbumin (OVA), and efficiently obtained OVA-specific human IgGs. Taken together, our study design provides a strategic platform to functionally decipher the alteration of phenotypes associated with incorporation of exogeneous genomic information and differing cellular environment.

## Results

### Generation of Tc rats with IGHK-NAC

Tc rat producing fully human Ab (designated as TC-mAb rat) was generated by using the IGHK-NAC as utilized for TC-mAb mouse generation (Fig. 1a)^7^. The IGHK-NAC was transferred from CHO donor cells to BLK2i-1 rat^12^ ESCs by microcell-mediated chromosome transfer (MMCT)^3^. The PCR confirmed the region on the IGHK-NAC and fluorescent *in situ* hybridization (FISH) analysis indicated that the IGHK-NAC was independently maintained without integration into the host genome in the rat ESCs (Fig. 1b). The rat ESCs carrying the IGHK-NAC that were karyotypically normal, were used for chimera production. The chimera rats were obtained by blastocyst injection of the rat ESCs and the IGHK-NAC was successfully transmitted to next generation through the germline.

**Figure 1.**
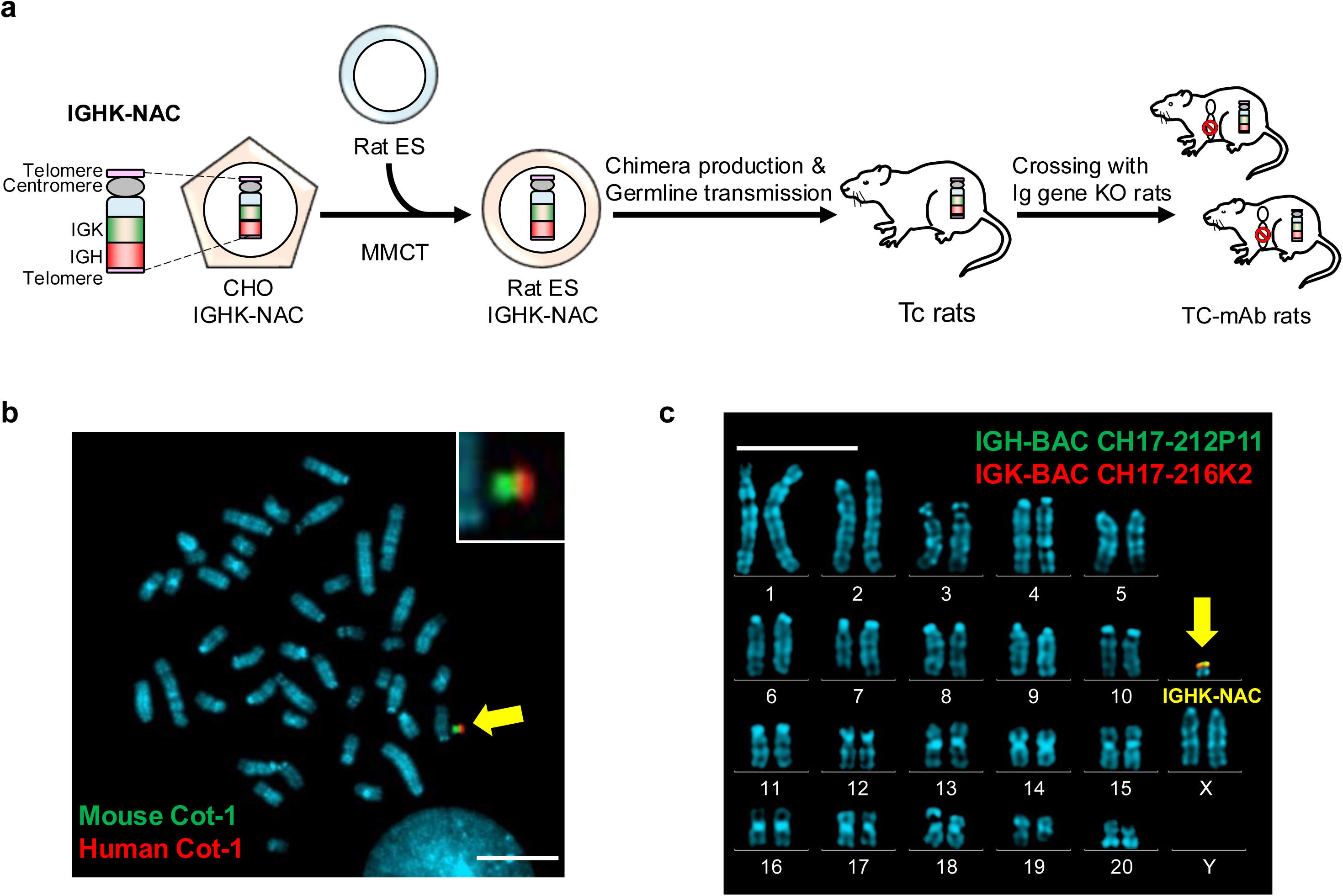
Generation of fully human Ab producing rats. **(a)** Strategy of generating TC-mAb rats by trans-chromosomic technology. The IGHK-NAC was transferred from CHO cells to rat ESCs by MMCT. Chimeric rats were generated from rat ESCs carrying the IGHK-NAC, and Tc rats were obtained through the germline. Tc rats carrying the IGHK-NAC were crossed with endogenous Ig gene KO rats to generate TC-mAb rats. **(b)** Representative image of metaphase FISH analysis with mouse Cot-1 (green) and human Cot-1 (red) detecting the IGHK-NAC in rat ESCs. Arrow indicates the IGHK-NAC which is enlarged in the inset. Scale bar indicates 10 μm. **(c)** Representative karyotyping result of lymphocytes in TC-mAb rats. Arrow indicates the IGHK-NAC. Red and green indicate the human *IGK* and *IGH* on the IGHK-NAC, respectively. Scale bar indicates 10 μm.

For complete Ig gene humanization, Ig genes KO (HKLD) rats for each endogenous Ig gene (*Igh*, *Igκ*, and *Igλ*) were generated by genome editing technology; i.e., injection of transcription activator-like effector nucleases (TALEN) into fertilized eggs. In the generation of each knockout strain of the IgM, Igκ, and Igλ genes, the absence of rat Ab protein in the blood after KO procedure was confirmed serologically by sandwich enzyme-linked immunosorbent assay (S. ELISA). (Supplementary Fig. 1). Tc rats carrying the IGHK-NAC were crossed with HKLD rats to generate Tc rats producing fully human Ab (Supplementary Fig. 2). The resultant rats (designated as TC-mAb rats) maintained the IGHK-NAC independently as the 43rd chromosome, which was confirmed by karyotyping (Fig. 1c and Supplementary Fig. 3).

### IGHK-NAC retention and Ig gene expression in TC-mAb rats

By detecting enhanced green florescent protein (EGFP) signal coded on the IGHK-NAC, all tissues of TC-mAb rats showed fluorescence signals (Fig. 2a). FISH analysis of the cells derived from several tissues indicated that a human DNA gene sequence on the IGHK-NAC were retained in most cells of TC-mAb rats (Fig. 2b, 2c, and Supplementary Fig. 4). Flow cytometry (FCM) analysis of a retention rate in lymphocytes, which is simple and reliable indicator for the MAC stability in the individuals^13^ indicated that the EGFP-positive rate of lymphocytes was 89.3−91.4% in TC-mAb rats (Fig. 2d), suggesting the stable retention of IGHK-NAC through generations. Transcription analysis of the Ig genes in various tissues by reverse-transcription (RT) PCR also indicated *IGH* and *IGK* were significantly expressed in the spleen and thymus (Fig. 2e, Supplementary Fig.5). Those of *IGK* were also detected in the intestine where B cells are accumulated. These results provide the evidence that the human genes are correctly transcribed in a tissue-specific manner in TC-mAb rats.

**Figure 2.**
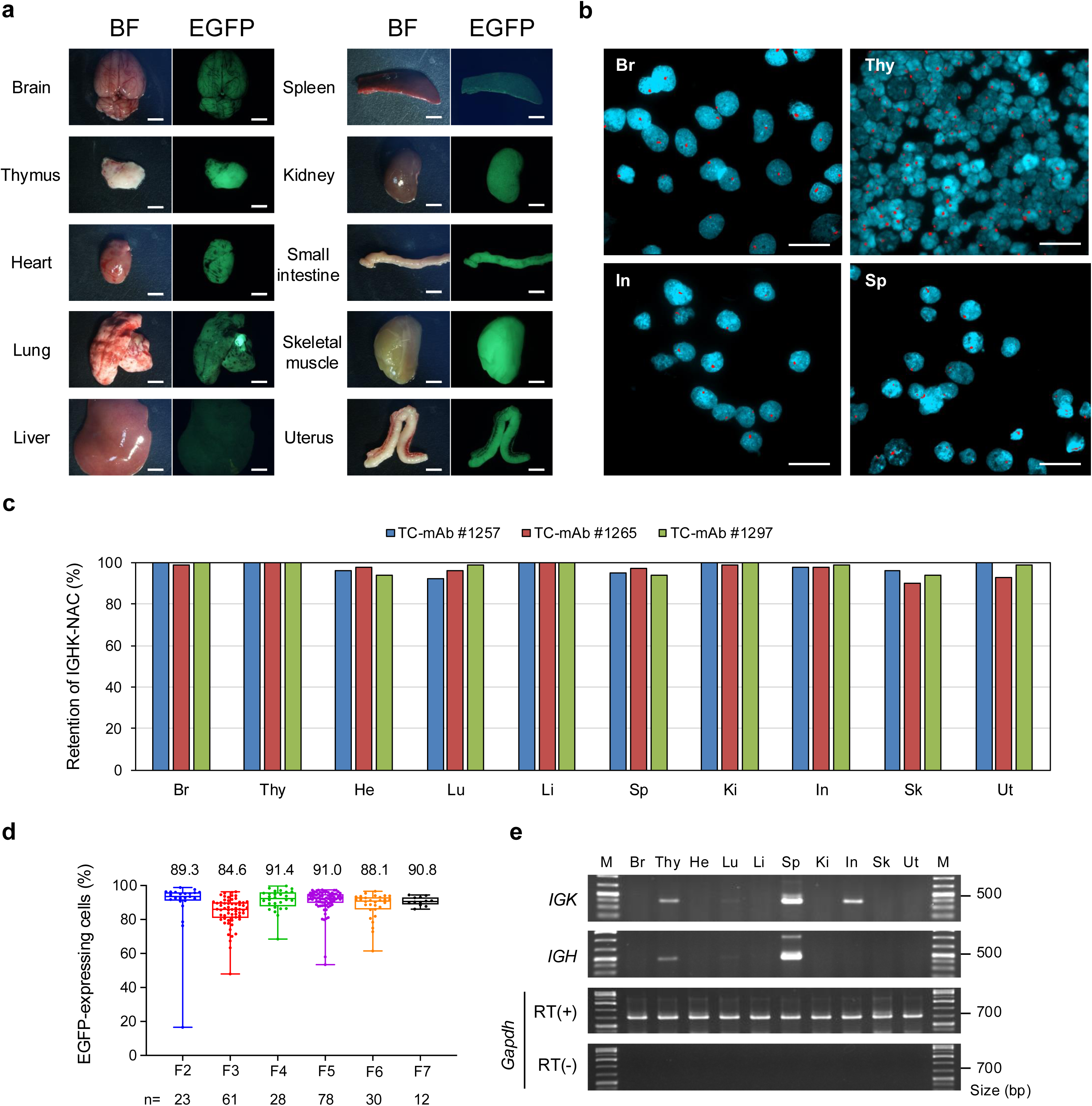
Stability of the IGHK-NAC and human gene expression in TC-mAb rats. **(a)** Bright and EGFP images of various tissues from a TC-mAb rat. Exposure times for each EGFP image were 40 ms for skeletal muscle, 150 ms for the brain, thymus, heart, kidney, small intestine, and uterus, and 400 ms for lung, 800 ms for liver, and 2 s for spleen. BF: Bright field. Scale bars indicate 5 mm. **(b)** Representative FISH images of each tissue from TC-mAb rat. The IGHK-NAC is detected in red by using human Cot-1 DNA as a probe. Br brain, Thy thymus, In small intestine, and Sp spleen are indicated. Scale bars indicate 20 μm. **(c)** FISH analysis of the IGHK-NAC retention rate in each tissue of TC-mAb rats. Br brain, Thy thymus, He heart, Lu lung, Li liver, Sp spleen, Ki kidney, In small intestine, Sk skeletal muscle, and Ut uterus. The numbers in each bar at the top of the figure indicate the individual numbers of the TC-mAb rats. **(d)** FCM analysis of the IGHK-NAC retention rate in lymphocytes by detecting EGFP signals in TC-mAb rats. F2−F7 represent each generation of TC-mAb rats. EGFP-positive average percentage is indicated at the top of the plot and the number of individuals analysed is indicated at the bottom of the figure, respectively. Box plots are indicated in terms of medians, bounds of box and whiskers (minima and maxima). **(e)** RT-PCR analysis of cDNA from various tissues of TC-mAb rat. Br brain, Thy thymus, He heart, Lu lung, Li liver, Sp spleen, Ki kidney, In small intestine, Sk skeletal muscle, and Ut uterus. Rat glyceraldehyde-3-phosphate dehydrogenase (*Gapdh*) gene is used as an endogenous control.

### Protein expression of human Igs in TC-mAb rat serum

Serum Ig levels in unimmunized 10-week-old TC-mAb rats were measured by S.ELISA. The average concentrations of human Igμ, Igγ, and Igκ were 33.1, 421.7, and 250.0 µg/ml, respectively (Fig. 3a and Supplementary Table 2). Of note, TC-mAb rats showed higher levels of Igγ than Igμ, indicating a similar pattern of Ab production to Wistar rats and humans^14^, in contrast with TC-mAb mice^7^. The average levels of Igγ and Igμ in TC-mAb rats are 2.75- and 0.064-fold compared to TC-mAb mice, suggesting that the progression of CSR from IgM to IgG is different, even though they have the same IGHK-NAC carrying full-length human Ig genes.

**Figure 3.**
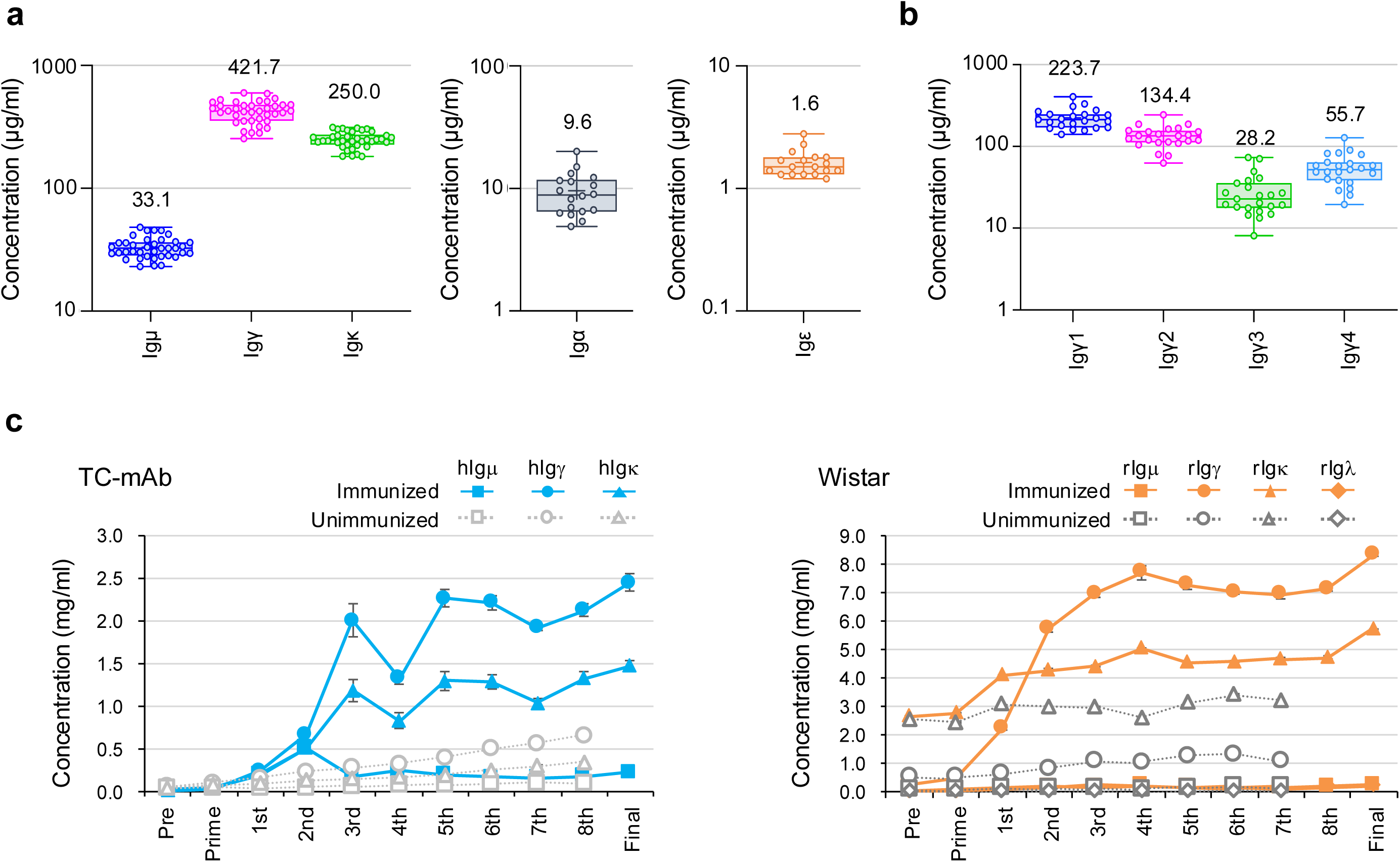
Production of human Ig in TC-mAb rats. **(a)** The serum concentration of different classes of human Igs in 10 week-old TC-mAb rats. The concentration of human Igμ, Igγ, Igκ, Igα, and Igε in TC-mAb rats (μ, γ, κ: *n=38*, α, ε: *n=18*) was measured. Each symbol represents one serum concentration. The numbers above boxplot indicate each average concentration (μg/ml). **(b)** The serum concentration of human IgG subclasses. The concentrations of Ig γ1, 2, 3, and 4 were measured in 10 week-old TC-mAb rats (*n=23*). The numbers above boxplot indicate each average concentration (μg/ml). Each symbol represents one serum concentration. **(c)** The serum concentration of Igμ, Igγ, Igκ and Igλ of TC-mAb (left) and Wistar rats (right) during OVA immunization. Human Igs are represented as hIgμ (square), hIgγ (round), and hIgκ (triangle) and rats Igs are represented as rIgμ (square), rIgγ (round), rIgκ (triangle), and rIgλ (diamond). Filled symbols and solid lines indicate immunized concentrations; unfilled symbols and dashed lines indicate non-immunized concentrations. Error bars indicate the standard deviation of triplicate measurements.

All four subclasses of Igγ, Igγ1, Igγ2, Igγ3, and Igγ4, were detected in TC-mAb rats, with average concentrations of 223.7, 134.4, 28.2, and 55.7 µg/ml, respectively (Fig. 3b and Supplementary Table 2). The distribution of human IgG subclasses in the human serum is reported as Igγ1>Igγ2>Igγ4>Igγ3^15^, suggesting the similar IgG subclass distribution between TC-mAb rats and humans. Furthermore, expression of Igα and Igε was also detected in the serum with average concentrations of Igα (9.6 µg/ml) and Igε (1.6 µg/ml) at 2.8- and 8.0-fold higher than TC-mAb mice^7^ (Fig. 3a and Supplementary Table 2).

To examine changes in serum Ig levels during the immune response, TC-mAb and Wistar rats were immunized with ovalbumin (OVA) at 2-weekly intervals and sera were collected over time. An increase in Ig levels was observed with each repeated immunization, reaching a plateau at the third booster step in both TC-mAb and Wistar rats (Fig. 3c and Supplementary Fig. 6). Comparing Ig levels before and after immunization in two TC-mAb rats, the concentration of human Igγ was increased from 177.6 and 31.4 (before immunization) to 2,276.6 (12.8-fold) and 2,448.8 µg/ml (78.0-fold). The concentration of rat Igγ in two Wistar rats were increased from 299.0 and 273.2 (before immunization) to 5,933.1 (19.8-fold) and 8,343.2 µg/ml (30.5-fold) (Supplementary. Fig.6a and 6b). In both TC-mAb and Wistar rats, their Igκ levels increased in parallel with Igγ levels after each booster. The Igμ levels are occasionally increased temporarily in the booster step but remained at low levels after the repeated boosters.

### B cell development in TC-mAb rats

FCM analyses of the peripheral blood mononuclear cells (PBMCs) obtained from TC-mAb, HKLD and Wistar rats were performed to evaluate whether TC-mAb rats are capable of reconstituting normal B cell development. As expected, the number of B cells (CD45R^+^ lymphocytes) was significantly lower in HKLD rats (average 1.4%, n=16) compared to parental rat strain (Wistar rats) (average 15.8%, n=6). The number was partially recovered in TC-mAb rats to approximately one-third (average 4.8%, n=166) of Wistar rats (Fig.4a, 4b, Supplementary Fig. 7, and Supplementary Table 3). Accordingly, the absolute number of splenocytes was clearly decreased in HKLD rats (average 8.7×10^7^ cells, n=3) and recovered to the comparable level as Wistar rats (average 3.1×10^8^ cells, n=3) in TC-mAb rats (average 2.7×10^8^ cells, n=3) (Fig. 4c).

**Figure 4.**
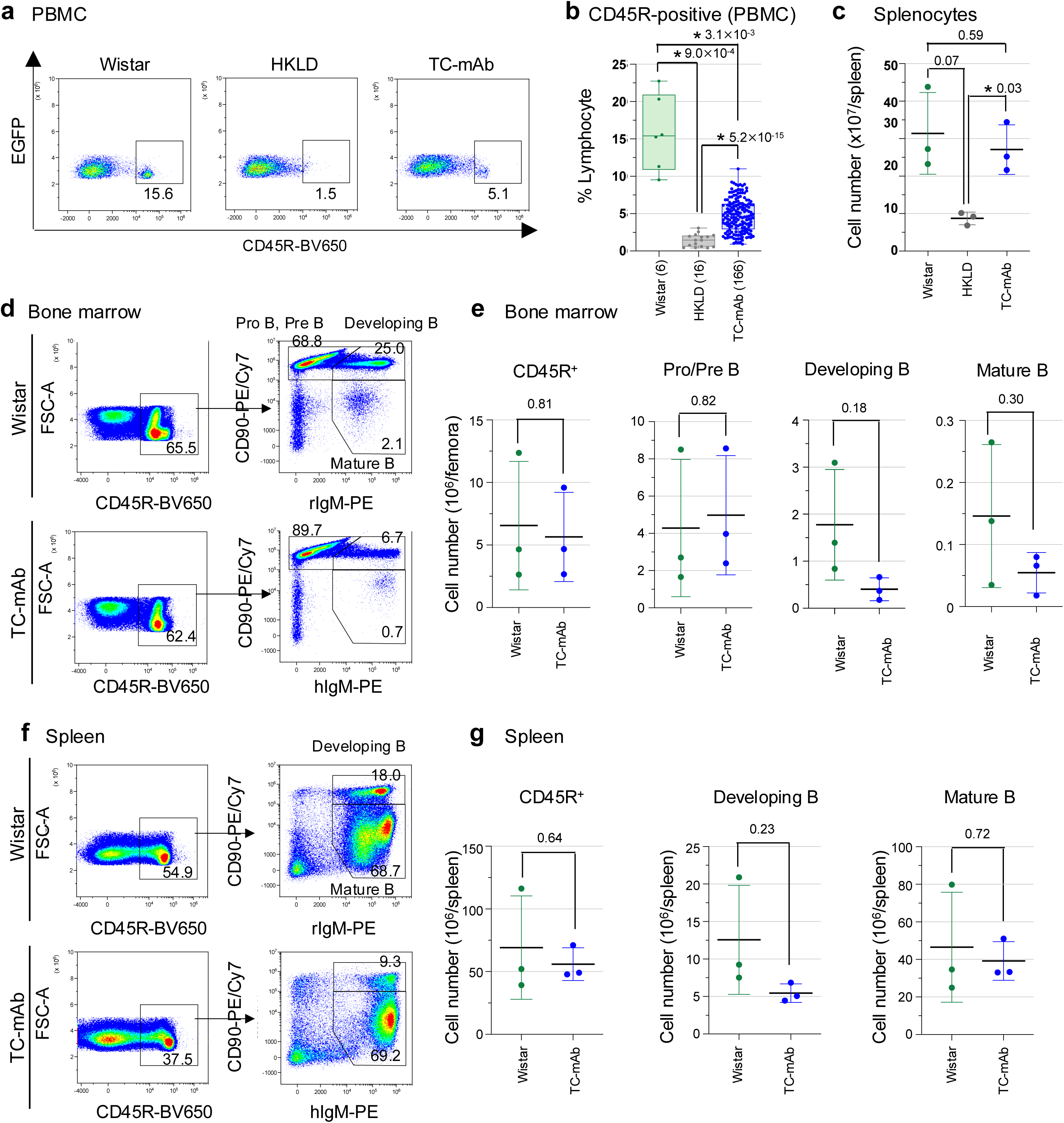
Expression and function of human Ig genes in TC-mAb rats. **(a)** Flow cytometry analysis of B cells in peripheral blood mononuclear cells of Wistar and TC-mAb rats. Panels indicate pan B cells (CD45R^+^) **(b)** Percentage of CD45R-positive B cells in the lymphocyte fraction of peripheral blood in Wistar (n = 6), HKLD (n = 16, Igh^−/−^, Igk^−/−^ and Igl^−/−^) and TC-mAb rats (n=166). Box plots are indicated in terms of medians, bounds of box and whiskers (minima and maxima). \**P* < 0.001 (unpaired Dunn’s test for multiple comparison). **(c)** The leukocyte numbers of spleen of Wistar, HKLD, and TC-mAb rats. **(d)** Flow cytometry of B cell compartments in the bone marrow of Wistar and TC-mAb rats. Panels indicate pan (CD45R^+^), Pro/Pre (Pro-B, Pre-B;CD45R^+^CD90^hi^IgM^−^), Developing (Dev-B;CD45R^+^CD90^hi^IgM^+^), and Mature (Mat-B;CD45R^+^CD90^lo^IgM^+^) B cells. Gating strategies are presented on Supplementary Figure 8. **(e)** The cell number of pan, pro/pre, developing, and mature B cell subsets in spleen (n=3) are presented. *P*-values between Wistar and TC-mAb rats of the cell numbers of B cell subsets were 0.81 (pan B cell), 0.82 (Pro B, Pre B), 0.18 (Dev-B), and 0.30 (Mat-B) in bone marrow, respectively. Error bars indicate the standard deviation of triplicate measurements. **(f)** Flow cytometry of B cell compartments in the spleen of Wistar and TC-mAb rats. Panels indicate pan (CD45R^+^), developing (Dev-B; CD45R^+^CD90^hi^IgM^+^), and mature (Mat-B;CD45R^+^CD90^lo^IgM^+^) B cells. Gating strategies are presented on Supplementary Figure 10. **(g)** The cell number of pan, developing, and mature B cell subsets in spleen (n=3) are presented. The 10-12 week-old unimmunized rats were analysed. All gating panels of B cell subsets are presented on Supplementary Figure 9 and 11. *P*-values between Wistar and TC-mAb rats of the cell numbers of B cell subsets were 0.64 (pan B cell), 0.23 (Dev-B), and 0.72 (Mat-B) in spleen, respectively. Error bars indicate the standard deviation of triplicate measurements. \**P*<0.05 (two-tailed unpaired Welch’s *t*-test).

CD90 and IgM expression in the cells from bone marrow and spleen was used to further analyse the developing B (including immature B, newly formed B, and early follicular B) and mature B cell (including mature follicular B and marginal zone B) populations^16,17^ (Supplementary Fig. 8 and 10). In comparison of the B cell population in the bone marrow between TC-mAb and Wistar rats, the absolute numbers of B cells (CD45R^+^) are comparable (6.6 and 5.6 ×10^6^ cells), Pro B/Pre B cells are slightly increased (4.3 and 5.0×10^6^ cells), and developing B (1.8 and 0.4×10^6^ cells) and mature B (0.15 and 0.05×10^6^ cells) are decreased in TC-mAb rats (Fig. 4d, 4e, Supplementary Fig. 9, and Supplementary Table 4). In the spleen, the absolute numbers of B cells (CD45R^+^) are comparable (69.2 and 56.0 ×10^6^ cells), developing B cells are decreased by half (12.6 and 5.5×10^6^ cells), and mature B cells are comparable (46.5 and 39.2×10^6^ cells) (Fig. 4f, 4g, Supplementary Fig. 11, and Supplementary Table 5).

Focusing on the mature B cell fractions in the spleen, which are important for Ab production, the absolute number is significantly recovered to the same levels as in Wistar rats (Fig. 4g). This recovery is clearly better in TC-mAb rats than in mice because TC-mAb mice have significantly less absolute numbers of follicular and marginal zone B cells in the spleen than wild-type mice. On the other hand, the number of Pre/Pro B cells and developing B cells in the bone marrow of TC-mAb rats tends to decrease compared to Wister rats.

### Analysis of human Ig heavy chain repertoire in TC-mAb rats

RNA samples extracted from splenocytes of five unimmunized TC-mAb rats, two OVA-immunized TC-mAb rats, and human pooled PBMCs from healthy adult donors (hPBMCs) were analysed for comprehensive profiling of the human Ab repertoire by next-generation sequencing (NGS) analysis, allowing analysis of frequency use of V(D)J gene segment, and junctional diversity in V(D)J gene recombination. Over 1.6 (*IGH*) and 0.8 million (*IGK*) qualified reads were collected from each sample, assembled into merged reads, and assigned V, D, and J segments using IgBLAST to be collated into data sets for subsequent analyses (Supplementary Table 6). The saturation of clonotype variations in each rarefaction curve were detected. The *IGH* and *IGK* transcripts in TC-mAb rats and hPBMCs were similar and over 91% productive.

All functional V segments (41 segments) were detected in the productive repertoire of TC-mAb rats (Fig. 5a and Supplementary Table 7). The broad distribution of V segment use (under 10%) was similar among all samples analysed excluding IGV2-5 gene segment of unimmunized TC-mAb rats (17.48%). Comparing the frequency of V segment use between hPBMCs and TC-mAb rats, *IGHV3-30* (5.40 and 0.12-0.14%) and *IGHV3-23* (12.45 and 3.45-4.01%) genes show a decrease and *IGHV6-1* (0.53 and 7.40-9.61%) gene show an increase in TC-mAb rats (Supplementary Table 7) presumably due to species specific differences in variable gene order at the genomic loci^18,19^. These results suggest that introducing the entire V region enabled considerable reproduction of the human repertoire in the rat spleen, which was further supported by Shannon-Weaver index of overall immune-repertoire diversity (Supplementary Table 8).

**Figure 5.**
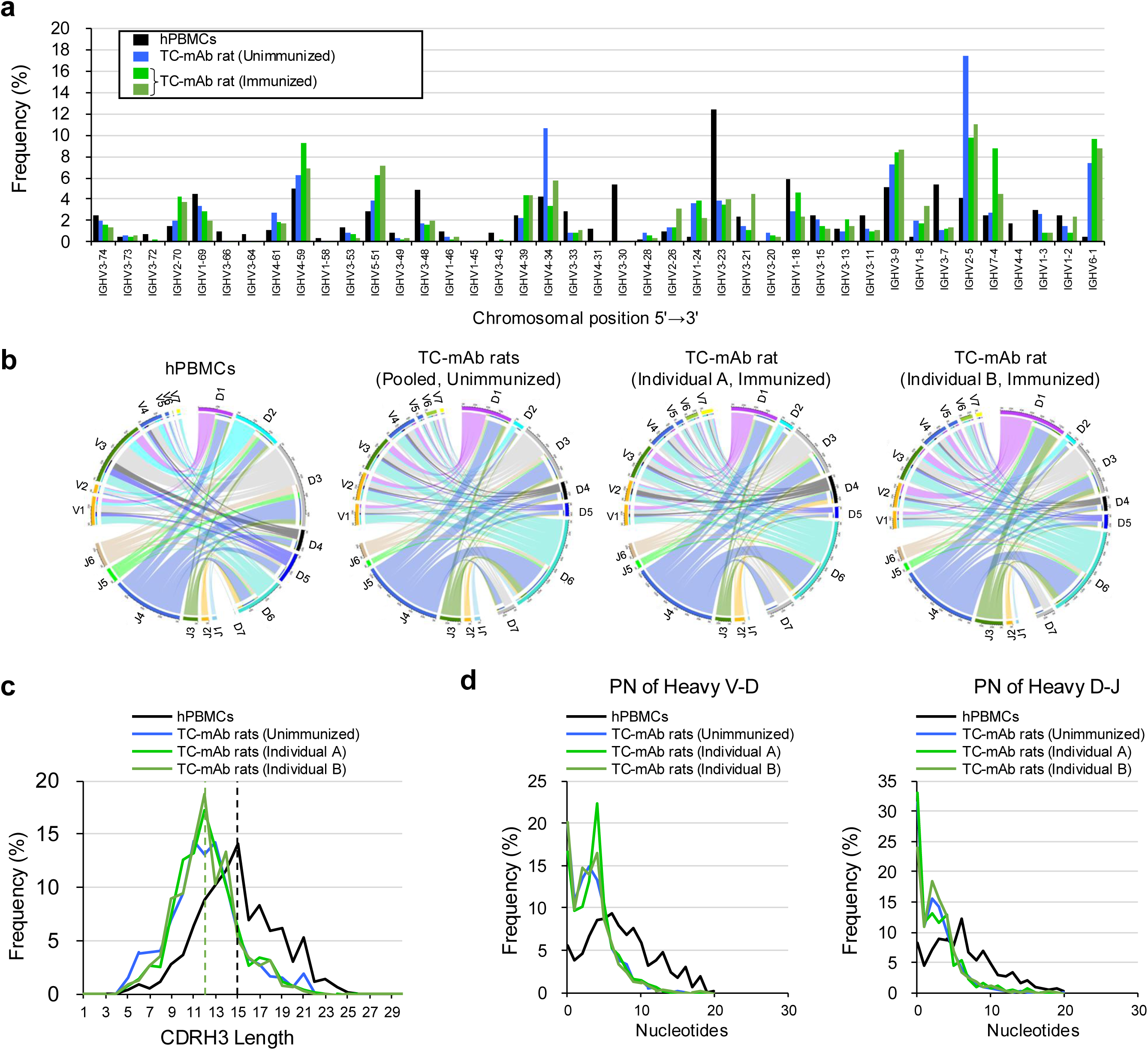
Repertoire analyses of heavy chain variable regions in TC-mAb rats. **(a)** Human heavy chain gene utilization in hPBMCs from healthy donors and in TC-mAb rats (female, 10-12 weeks old) with and without immunization (female, two individuals, 25-26 weeks old). Percentage frequency use of V gene segments in healthy human donors (grey), TC-mAb rats with (light and dark green), and without immunization (blue) are represented. **(b)** Circos plots comparing VDJ gene association. The gene segments are grouped as subfamilies and are shown together with the first digit of their allele name. Links indicated the relative frequencies of specific VDJ combinations, and wider links indicate higher frequencies of recombination. **(c)** The relative CDRH3 length. Black dot line indicates average length of CDRH3 in hPBMCs (14.98 amino acids), and blue (unimmunized, 12.01 amino acids), light green (OVA-immunized individual A, 12.12 amino acids), and dark green (OVA-immunized individual B, 12.23 amino acids) dot lines are also shown overlapping. **(d)** The relative P and N nucleotide addition at joining positions within the CDRH3 of productive rearrangements.

The use of D segment was mostly similar between TC-mAb rats and hPBMCs, although *IGHD6-19* gene (21 bp) segment is the most frequently used in TC-mAb rats (22.16-24.05%) compared to hPBMCs (5.99%). This finding is different from the most frequently use of *IGHD3-10* gene (31 bp) in hPBMCs (11.21%) and TC-mAb mice (30.38-33.46%)^7^ (Supplementary Table 7). All six J segments are used, and the *JH4* and *JH6* gene segments are dominantly used similar to TC-mAb mice and hPBMCs (Supplementary Table 7). Accordingly, circos plots of V, D, and J segment combinations in rearranged Ig heavy chains reveal that the frequency of V/D/J use and the combinations of VH regions are similar with hPBMCs and TC-mAb rats but the frequency of *DH2* and *DH3* segment families is decreased and that of *D6* segment family is increased (Fig. 5b and Supplementary Fig. 12). This result indicates that the V/D/J gene use and combination in TC-mAb rats is similar but slightly different to those in hPBMCs and TC-mAb mice.

### CDRH3 length distribution and amino acid composition

The length of complementary-determining region 3 of the heavy chain (CDRH3) is highly diverse and a key determinant of specificity in antigen recognition, and is one of the hallmarks of fully human Ab production in animals^20^. The distribution of CDRH3 length in TC-mAb rats is shorter than human with average length of 12.01 (unimmunized), 12.12 (OVA-immunized individual A), and 12.23 amino acids (OVA-immunized, individual B) in TC-mAb rats instead of 14.98 of humans (Fig. 5c). The length of CDRH3 is dependent on the usage frequency of the DH segment families because there are segments of varying length in the DH region (Supplementary Table 9). The *DH2* and *DH3* segment families have larger lengths (average 30.25 and 32.20 bp, respectively) and are frequently used in human and TC-mAb mice^7^, contributing to the long CDRH3 length. However, TC-mAb rats instead use the *DH6* segment family frequently, which has an average length of 19.5 bp (Fig. 5b and Supplementary Fig. 12 and 13). Calculated from the length and frequency use, the average length of the DH region is decreased by 3.1 bp, consistent with TC-mAb rats (unimmunized, 18.6 bp) and hPBMCs (21.7 bp).

Shorter distributions of PN nucleic acid addition at V-D and D-J junctions were observed in TC-mAb rats compared with hPBMCs (Fig. 5d and Supplementary Table 10), resulting in decreased average length of CDRH3 to 8.86 bp in TC-mAb rats (unimmunized, 5.78 bp) compared to hPBMCs (14.64 bp). The same reduction is also observed in genetically engineered rats^19^ and TC-mAb mice^7^, suggesting that the nucleotide addition in V(D)J recombination is regulated in a host-dependent manner. Altogether, the shorter CDRH3 region of TC-mAb rats is due to higher frequency use of the *DH6* segment family and reduced PN nucleotide addition (Fig. 5c). However, this reduction does not affect the amino acid composition, which is comparable between hPBMCs and TC-mAb rats (Supplementary Fig. 14a).

### Repertoire analysis of human IGK chains in TC-mAb rats

The NGS data indicated that all of the 18 potentially functional proximal V segments, as well as 11 functional distal V segments in TC-mAb rats and 9 out of 16 functional distal V segments in hPBMCs were used (Fig. 6a and Supplementary Table 11), closely consistent with the previous analyses of TC-mAb mice^5^. The Circos plots of V-J segments show that the frequency use and assembled combinations of the VK region are highly similar between hPBMCs and TC-mAb rats (Fig. 6b, Supplementary Fig. 15, and Supplementary Table 12). The Shannon-Weaver index also supports this diversity (Supplementary Table 8). No significant difference exists in J segment use with or without immunization. The CDRL3 length, the number of nucleotide additions, and amino acid composition are all comparable between hPBMCs and TC-mAb rats (Fig. 6c and Supplementary Fig. 14b).

**Figure 6.**
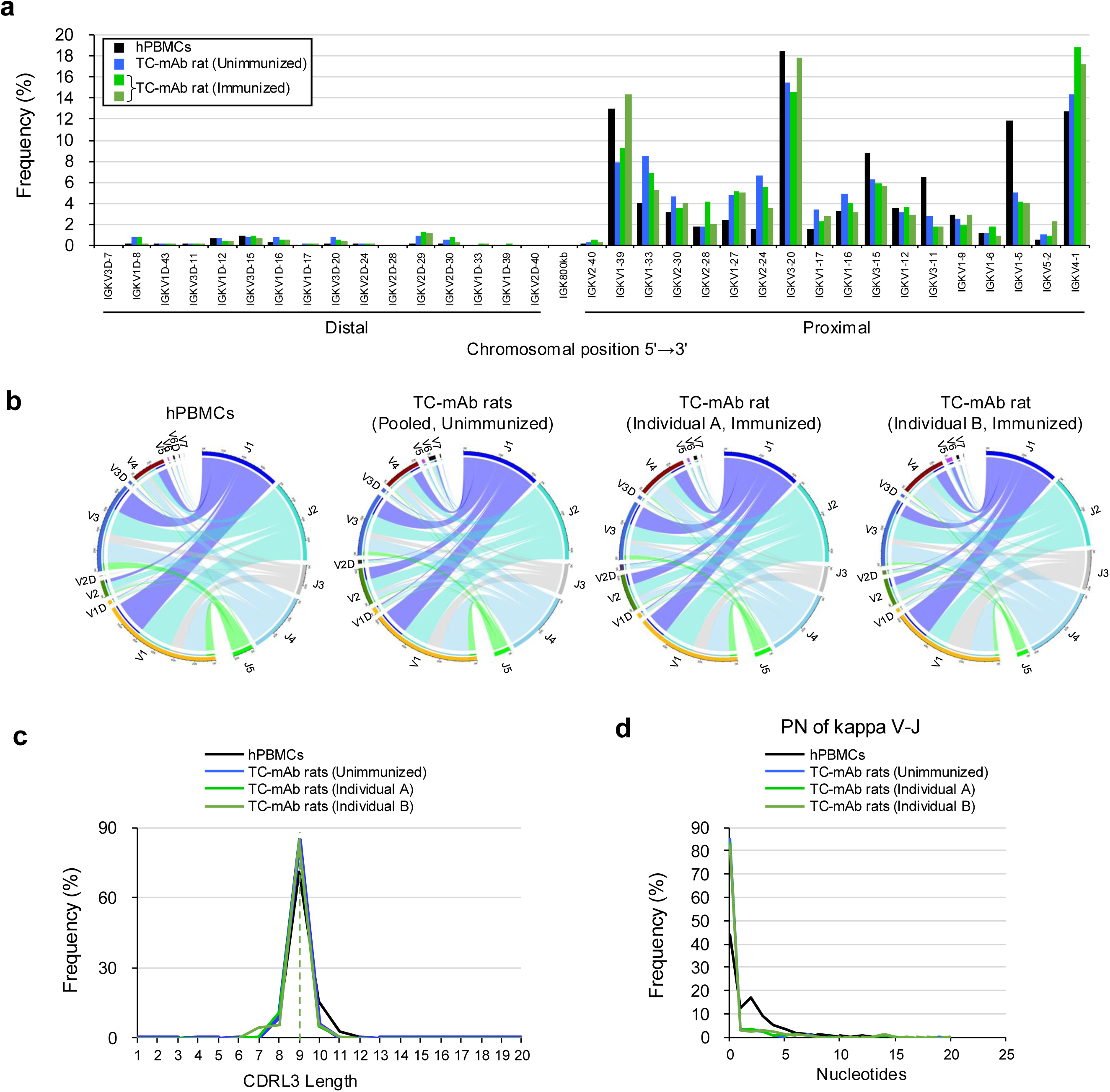
Repertoire analyses of kappa chain variable regions in TC-mAb rats. **(a)** Human kappa chain gene utilization in hPBMCs from healthy donors and in TC-mAb rats with and without immunization (Individuals are same as Fig. 5). Percentage frequency use of V gene segments in healthy human donors (grey), TC-mAb rats with (light and dark green), and without immunization (blue) are represented. **(b)** Circos plots comparing VJ gene association. The gene segments are grouped as subfamilies and are shown together with the first digit of their allele name. Links indicated the relative frequencies of specific VJ combinations, and wider links indicate higher frequencies of recombination. **(c)** The relative CDRL3 length. Black dot line indicates average length of CDRL3 in hPBMCs (9.09 amino acids), and Blue (unimmunized, 8.97 amino acids), light green (OVA-immunized individual A, 8.94 amino acids), and dark green (OVA-immunized individual A, 8.92 amino acids) dot lines are also shown overlapping. **(d)** The relative P and N nucleotide addition at joining positions within the CDRL3 of productive rearrangements.

### Somatic hypermutation in TC-mAb rats

SHM is a necessary process to produce high-affinity Abs against antigens and the mutations accumulate in variable regions. The variable region of heavy (VH) and kappa (VK) regions of TC-mAb rats are analysed, and the top 50 frequently used clone lineages of VH and VK sequences are compared in unimmunized pools (n=5) and immunized (two individuals) TC-mAb rats (Fig. 7). Contrasting the unimmunized and immunized results, mutations are clearly detected within the germline sequence in the entire region of VH and VK including complementary determining region (CDR) 1, 2, 3 and frame work region (FR) 3. This result is reasonable because not only the CDR regions but also the FR3 region containing the D-E loop has been reported to be involved in Ab-antigen binding^21^. The expected value of SHM; i.e., the expected number of nucleotide mutations in the VH of TC-mAb rats is 2.21 (unimmunized pools, n=5), 3.68 (immunized individual A), and 3.70 (immunized individual B) (Supplementary Table 13). The expected number of nucleotide mutations of the VK is 2.23 (unimmunized pools, n=5), 3.75 (immunized individual A), and 3.89 (immunized individual B). Intriguingly, in the TC-mAb mice, this VH value is 0.78 (unimmunized pools, n=5) and 0.90 (immunized, n=1), and that of VK is 1.15 (unimmunized pools, n=5) and 1.47 (immunized, n=1), suggesting that SHM frequency is clearly higher in TC-mAb rats than in TC-mAb mice^7^.

**Figure 7.**
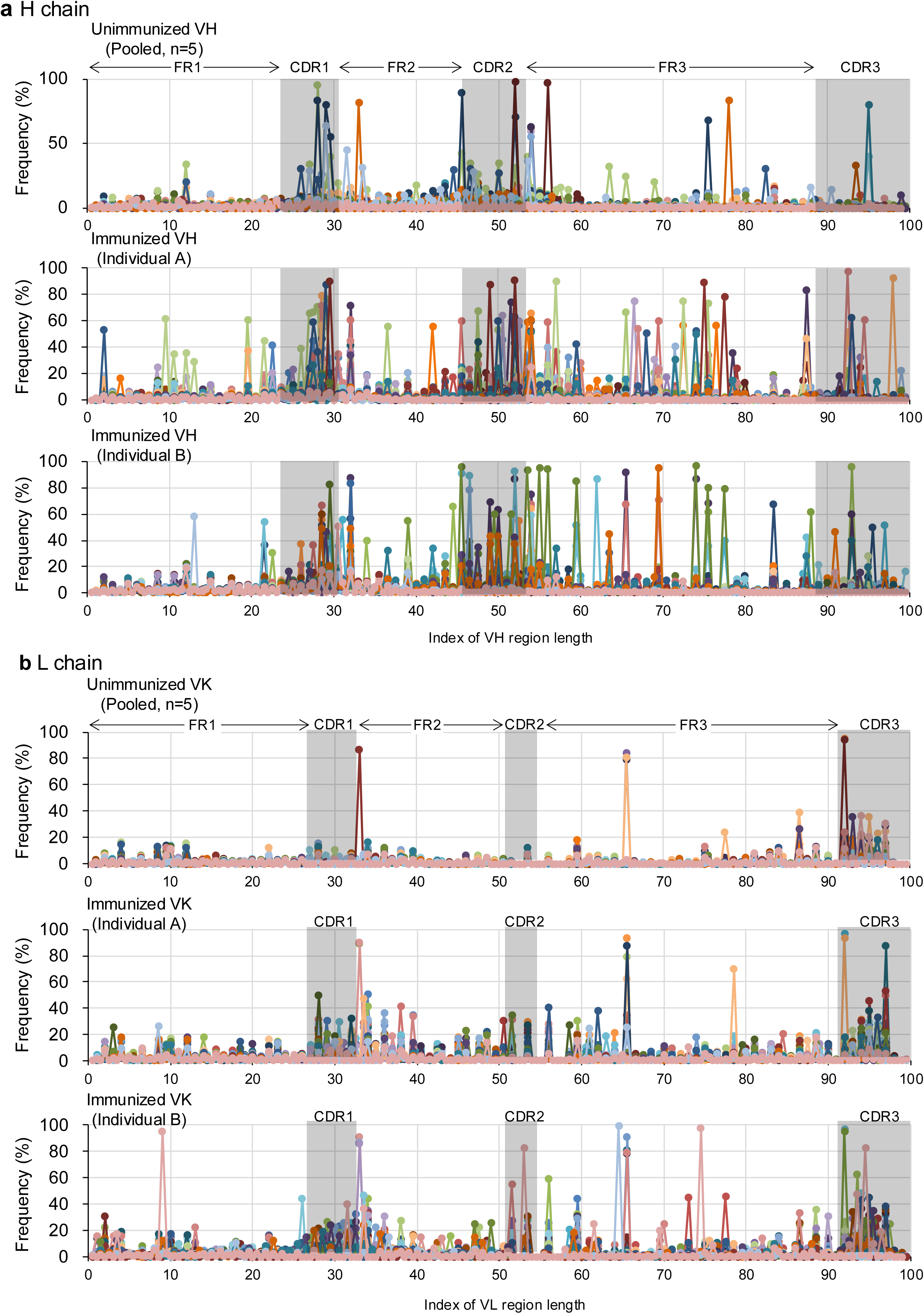
Somatic hypermutation analysis. The frequency of SHM at every position of the variable region. It was calculated as described in the Methods and is represented for VH and VK chains in OVA-immunized TC-mAb rats **(a, b)** and unimmunized TC-mAb mice **(c, d)** (Individuals are same as Fig. 5).

Circular dendrograms of phylogenetic trees are drawn to show the mutation mapping and expansion of clonotypes based on their amino acid sequence in each clone lineage according to the same definition and classification as reported in the previous study^7^. The circular dendrogram of the 10 most frequent clonal lineages of the VH (Supplementary Fig. 16) and VK (Supplementary Fig. 17) is presented. Similar with TC-mAb mice, expansion of clonotypes with SHMs was induced by immunization of TC-mAb rats, leading to antigen-driven VH and VK sequence diversification, which should contribute to the generation of high affinity Abs.

### Producing antigen-specific fully human Abs in TC-mAb rats

To evaluate the immune response for producing human Abs in TC-mAb rats, the age-matched TC-mAb (n=2) and Wistar rats (n=4) were immunized with OVA. The OVA-specific human Igγ response in TC-mAb rats was robustly elevated by immunization and was comparable to that of WT rats until it reached plateau (Fig. 8 a, b and Supplementary Fig. 18). This immune response is not similar with TC-mAb mice because TC-mAb mice require additional two or three booster steps^7^, and is consistent with the Ig level in anti-sera and increased CSR and SHM frequency in TC-mAb rats (Fig. 3 and 7).

**Figure 8.**
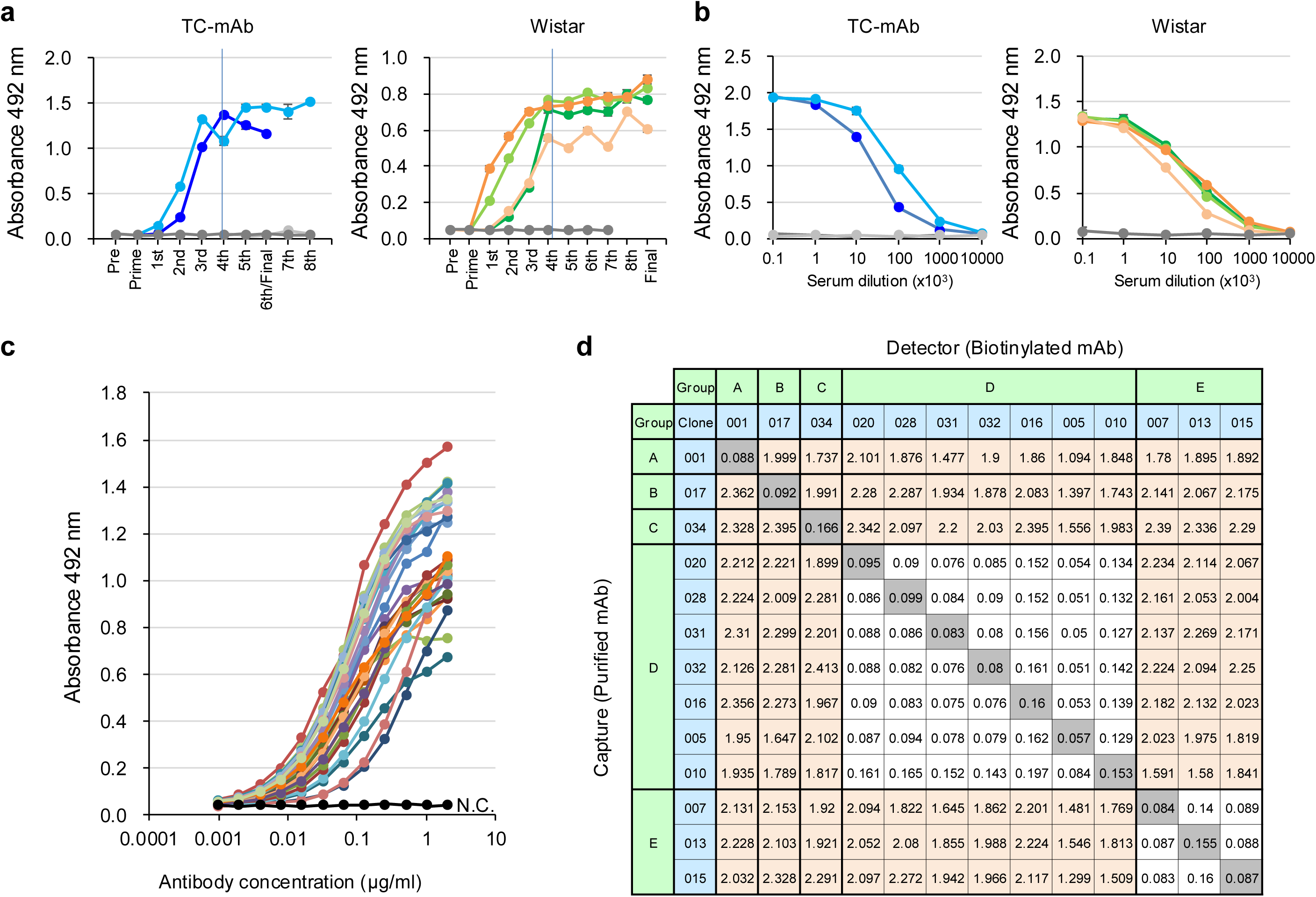
Analysis of antigen-specific mAbs obtained from TC-mAb rats. **(a)** OVA-specific Ab titre. The titres of Abs against OVA were measured by ELISA using a species-match secondary Ab. An anti-human IgG-Fc-HRP conjugate was used for TC-mAb rats, and an anti-rat IgG-Fc HRP-conjugate for Wistar rats. Error bars indicate the standard deviation of triplicate measurements. **(b)** Comparison of the anti-sera titres. The anti-sera samples were collected from TC-mAb rats after the 5 and final (dark blue) or 8 and final (light blue) booster, and from Wistar rats after the 8 boosters. Samples were serially diluted by 1/10 from 10^2^ to 10^7^ and used to determine the OVA-specific titre by ELISA. Error bars indicate the standard deviation of triplicate measurements. **(c)** The antigen-specific titre of the 27 mAbs obtained. Error bars indicate the standard deviation of triplicate measurements. N.C. indicates negative control using Ab-free sample. **(d)** Epitope distribution analysis by competitive assay. Clone names (sky blue columns) indicate the mAbs used in this analysis. Clear signals are observed for combinations that bind to different epitopes (highlighted as the orange column). The mAbs bind to the same epitope are indicated by groups (green columns). The 13 mAbs analyzed are classified into 5 groups, A to E.

Hybridoma cells were generated from OVA-immunized TC-mAb rats, and 30 OVA-specific fully human mAbs were successfully obtained (Supplementary Table 14). Antigen binding analysis by ELISA (Fig. 8c) and epitope distribution analysis by competitive assay (Fig. 8d) for 27 mAbs, IgG1 and IgG2 subclasses, showed that various kinds of antigen-specific Abs were obtained from immunized TC-mAb rat. Sequence analysis of the obtained mAb variable regions indicates a high degree of humanness determined using T20 score analyser^22^ and a high frequency of SHM in CDR regions and FR3 (Supplementary Fig. 19 and 20). Furthermore, most of the mAbs obtained are IgG1 subclass (27/30, 90.0%), also supporting the improved frequency of CSR from IgM to IgG in TC-mAb rats. Altogether, TC-mAb rats efficiently produce a variety of mAbs, consistent with immune responses similar to that of Wister rats (Fig. 3). The large number of lymphocytes obtained for rats due to their large individual size allowed for a series of analyses from the same individuals.

## Discussion

Advantage of the artificial chromosome vector is that the same vector can be utilized in various species if chromosome transfer is possible. In fact, TC-mAb rats were generated by introducing the IGHK-NAC only once without multiple genetic modification of embryonic stem cells (ESCs), reducing the cost and time required for establishing genetically engineered rat strain. In particular, Tc animals are readily produced by combining with the recently developed gene KO method of fertilised egg using genome editing technology. This approach is highly efficient when compared with a genetically engineered rat chimeric strain Omni rat^18^ that utilized repeated chromosomal modifications by DNA microinjection into oocytes^18^. Additionally, in contrast with our TC-mAb rats, since Omni rats do not contain the full-length human Ig gene locus, human V(D)J recombination is different from humans^19^. The generation of TC-mAb rats described herein demonstrates the practicality of combining these genomic engineering techniques, and represents the feasibility of introducing exogeneous large genes with the same structure into rodents for comparison.

A summary of TC-mAb rats, TC-mAb mice, Omni rats, and Wistar rats is indicated elsewhere (Table 1). Multiple biological processes are involved with efficient Ab expression; most notably, CSR and SHM efficiencies. In this study, we observed markedly different concentrations of IgG subclasses and induction patterns of antigen-specific IgGs between TC-mAb rats and mice. Specifically, higher concentrations of human Igγ, Igα, and Igε but lower concentration of Igμ suggest the improved CSR frequency of the human Ig gene in TC-mAb rats, although Ig levels of TC-mAb rats are not completely recovered to the levels reported in humans^14^. Another important factor to consider is SHM efficiency, and our NGS analysis revealed a higher frequency of SHM in TC-mAb rats including a 4.1-fold higher frequency in the VH region than TC-mAb mice^7^. Consistent with these findings, we subsequently determined the efficient production of hybridomas, and the generation of various human anti-OVA IgGs. Collectively, compared to TC-mAb mice, while TC-mAb rats have a conserved human Ig repertoire, significant improvements are observed in CSR and SHM frequencies, although a shortening of PN addition is still retained. These results suggest that the V(D)J recombination process depends on the structure of Ig gene loci, and this machinery is encoded on genome structure, but CSR and SHM frequency, and PN addition are regulated by host proteins.

**Table 1.** Characteristics of TC-mAb animals for antibody production.

| Table 1\| Characteristics of TC-mAb animals for antibody production |  |  |  |  |
| --- | --- | --- | --- | --- |
|  | Wistar rats | TC-mAb rats | TC-mAb mice | Omni rats |
| Carrier of human Ig | – | IGHK-NAC | IGHK-NAC | Targeting insertion |
| human Ig-genes | – | Full length<br>*IGH* and *IGK* | Full length<br>*IGH* and *IGK* | Partial length<br>*IGH* and *IGK* |
| Stability of Ig-genes | Stable<br>(Endogenous) | Stable | Stable | Stable |
| Spleen size and number of lymphocytes | Normal | Normal | Decreased by half | Normal |
| IgG concentration in anti-sera | Normal | Decreased | Significantly decreased | Normal |
| B cell development | Normal | Adverse in bone marrow | Adverse in bone marrow<br>and spleen | Normal |
| Immune response | Normal | Normal | Delayed | – |
| Antibody titre | Normal | Normal | Normal | – |
| Plasma cell / plasmablast differentiation | Normal | – | High | – |
| Antigen-specific B cell production | Normal | High | – | – |
| Hybridoma production frequency | Normal | No adverse | High | – |
| Somatic hypermutation frequency | Normal | *IgH*: similar to humans<br>*IgK*: about half for humans | Adverse<br>(40–50% of TC-mAb rats) | – |
| Class-switch recombination | Normal | IgG>IgM | IgM>IgG | – |
| IgG subclass (unimmunized) | Normal | IgG1>2>4>3 | IgG1≈2>3,4 | – |
| Human Ig repertoires | – | Closely similar to hPBMCs | Closely similar to hPBMCs | Different from hPBMCs |

–: Indications are unknown

CSR and SHM functions are widely known to be regulated by several proteins, including activation-induced cytidine deaminase (AID)^23^. Mutation study of AID indicates that separate domains of AID are required for CSR and SHM, respectively, and cofactors with different domains of AID have been responsible for their regulation^24^. In particular, a strict regulation machinery that causes mutations in CDR1, 2, and 3 regions has been proposed to be regulated by a licensing mechanism^25^ with several proteins such as RNA polymerase II and SPT5. To clarify whether these genes affect the frequency of CSR and SHM, it is necessary to analyse whether increased efficiency can be observed by humanizing the host proteins associated with CSR and SHM or genetic murinization of Ig gene constant and regulatory regions in TC-mAb mice.

In TC-mAb rats, GC B cell formation is possibly improved because the population of mature B cells in the spleen was recovered to the same level as Wistar rats. Furthermore, analysis of antigen-specific B cells using biotinylated OVA showed that significant increase of OVA-specific surface IgG expressing cells in the spleens of TC-mAb rats immunized with OVA, supporting the efficient differentiation of B cells to antigen-specific Abs production. The adverse effects are detected in B cell differentiation of CD45R^+^ B cells and developing B cells in PBMCs, bone marrow, and spleen, indicating the inefficient B cell maturation process in the bone marrow of TC-mAb animals, possibly the result of having only one copy of the human Ig locus on the IGHK-NAC. TC-mAb animals have only one set of *IGH* and *IGK* on their artificial chromosomes and no alleles. This may cause B cells that are unable to produce usable Abs to rapidly undergo apoptotic cell death through allelic exclusion process^26^, possibly leading to a decrease in developing B cells in TC-mAb rats.

In conclusion, although TC-mAb rats and mice show different parts of the Ab-producing capability, both are effective animals for the development of human Ab therapeutics. Furthermore, this study allows us to estimate the part of the Ig gene loci where information is conserved in its structure and the part that is modified by the animal host protein during the process of Ab production. Regarding the differences between Tc mice and rats, it is expected that more detailed analysis will be possible by using genetically engineered Tc mice, such as humanized AID genes with amino acid differences to mouse counterparts^27^ and analysing CSR and SHM efficiencies. Therefore, the strategic approach we have developed here using Tc animals contributes significantly to advance functional analysis of human genes.

## Methods

### Ethics declarations

All animal experiments were approved by the Animal Care and Use Committee of Tottori University (Permit Numbers: 14-Y-23, 15-Y-31, 16-Y-20, 17-Y-28, 19-Y-22, 20-Y-13, 20-Y-31, 21-Y-26, and 22-Y-36) and the National Institutes of Natural Sciences (16A027). All experiments were carried out in compliance with the ARRIVE guidelines, other relevant guidelines and regulations.

### Cell culture

CHO K1 cells (Riken BRC, Ibaraki, Japan: RCB0285) carrying the IGHK-NAC were maintained at 37℃ in Ham’s F12 nutrient mixture (Wako, Tokyo, Japan) supplemented with 10% FBS and 800 μg/ml G418. A parental rat ES cell line (BLK2i-1, RGD ID:10054010) and the BLK2i-1 microcell hybrid clones were maintained on mitomycin C-treated neomycin-resistant MEFs (Oriental Yeast Co., Ltd., Tokyo, Japan) as feeder layer, as described previously^28^.

### MMCT

MMCT was performed as described previously^29^. Briefly, rat ES cells were fused with microcells prepared from the donor CHO cells carrying the IGHK-NAC, and selected with 150 μg/ml G418.

### Chimeric rat production

Chimeric rats were produced as reported previously^30^. BLK2i-1 microcell hybrid clones were microinjected into the blastocoel cavity of Crlj:WI E4.5 blastocysts (9-11 ES cells per blastocyst). Then, the blastocysts were transferred into ureti of pseudopregnant recipient rats at 3.5 d.p.c. (12-15 blastocysts per recipient). The contribution of the ES cells in chimeric rats was determined by coat color and monitoring EGFP.

### TALEN mRNA preparation

Platinum TALEN construction and analysis of their activity were performed as described previously^31^. Each TALEN mRNA was prepared with mMESSAGE mMACHINE T7 Ultra Kit (Thermo Fisher Scientific, Waltham, MA, USA), following the manufacturer’s instructions. Synthesized poly(A)-tailed TALEN mRNA was cleaned up using the RNeasy Mini kit (Qiagen, Hilden, Germany). Each TALEN target sequence is left 5’-TCC CCC TCG TCT CCT GCG-3’ and right 5’-TGG CCA CCA AAT TCT CAT-3’ for *Igh*, left 5’-TAC AGC ATG AGC AGC AC-3’ and right 5’-TGA CTT TCA TAG TCA GCC-3’ for *Igκ*, left 5’-TAA TTT GAT CCA GCC CA-3’ and right 5’-TGG GTT CAG CAC CTG GGC-3’ for *Igλ* up, and left 5’-TCA CGA AGA GAA CAC TGT-3’ and right 5’-TAG GAA CAC TCAGCA CGG-3’ for Igλ down.

### Genome editing by TALEN

For genome editing, pronuclear-stage zygotes were collected from oviductal ampullae of Crlj:WI female rats superovulated and mated with male rats^32,33^. They were injected with each TALEN mRNA solution (10-20 ng/μl) and cultured for 24 h in mKRB medium at 37 ℃ under 5% CO_2_. All surviving embryos were transferred into oviductal ampullae of pseudopregnant rats at 0.5 days postcoitum (24-48 embryos per recipient). Successful genome editing was confirmed by Cel-1 assay and sequencing of Ig genes.

### FISH analysis

Cultured cells, lymphocytes and homogenized tissue samples were fixed with Carnoy’s solution (methanol and acetic acid ratio 3:1). The slides were prepared by standard methods. FISH analyses were carried out by using digoxigenin-labelled DNA [human COT-1 DNA/mouse COT-1 DNA (Invitrogen, Carlsbad, CA, USA) and IGK-BAC (CH17-216K2)] and biotin-labelled DNA [human COT-1 DNA/mouse COT-1 DNA and IGH-BAC (CH17-212P11)] as described previously^29^. Chromosomal DNA was counterstained with DAPI (Sigma-Aldrich, St. Louis, MO, USA). Images were obtained by using an AxioImagerZ2 fluorescence microscope (Carl Zeiss GmbH, Jena, Germany).

### RT-PCR analysis

RT-PCR analysis was performed as described previously^6,7^.

### Serum concentration of Abs in TC-mAb and Wistar rats

Abs concentration in serum of Ig KO rats was analyzed by ELISA. Plates were coated with mouse anti-rat IgM antibody-biotin and detection was carried out by mouse anti-rat kappa/lambda light chain-HRP (Bio-Rad, Hercules, CA, USA: MCA1296P), mouse monoclonal K4F5 anti-rat kappa light chain-HRP (Abcam, Cambridge, UK: ab99692) and mouse anti-rat lambda light chain-HRP (Bio-Rad: MCA2366P), respectively. Purified rat IgM kappa (Bio-Rad: PRP08) and purified rat IgMλ (Southern Biotech, Birmingham, AL, USA: 0120-01) were used as standard. 3,3′,5,5′-tetramethylbenzidine (TMB) (Nacalai Tesque, Kyoto, Japan) was used as substrate, and absorbance at 450 nm was measured by a spectrophotometer (BioTek instruments, Winooski, VT, USA).

Abs concentration in serum of human Abs was measured using the same procedure. Goat polyclonal IgG specific to each Ig class (Bethyl laboratories) as capture (unconjugated) and detection (HRP-conjugated) Abs were used in measurements of hIgγ, hIgμ, hIgα, hIgε, and hIgκ (Capture and detector Abs are listed in Supplementary Table 1). 1-Step™ TMB ELISA Substrate Solutions (#34029, Thermo Fisher Scientific) was used as substrate. All steps were performed at room temperature. Samples and standards were triplicated and averages and standard deviations were calculated. The concentration of hIgγ subclasses, Igγ1, Igγ2, Igγ3, and Igγ4, were measured using IgG Subclass Human ELISA Kit (# 99-1000, Thermo Fisher Scientific) following the product’s instruction.

### Immunization

Ovalbumin (OVA) was obtained from Sigma (A7641) and dissolved in PBS or deionized water at 1 mg/ml. OVA (250 μg/250 μl) was then mixed with an equal volume of either Freund’s or Sigma adjuvant (Sigma Adjuvant S6322; Sigma CFA F5881, Sigma) prepared following the manufacturer’s instructions. Prime and booster injections were given intraperitoneally (*i.p.*) every 2 weeks. Doses varied depending on the administration route and experimental requirements and were determined according to the relevant JP Home Office animal license for the procedure. Final boosters were delivered without adjuvant intravenously (*i.v.*) via the tail vein.

### Cell staining and Flow cytometry

Analyses of B cell development in Wistar and TC-mAb rats were performed by flow cytometry. For PBMC samples, 50 µl peripheral blood was mixed with 50 µl anti-rat CD45R Ab diluted in staining buffer (BD Biosciences Brilliant stain buffer) and incubated on ice for 30 min before hemolysis treatment. Isolated cells from bone marrow and spleen were pre-incubated with FcR blocker (anti-rat CD32) in 50 µl FCM buffer (PBS with 5% FBS) followed by the addition of 50 µl Abs mixture (Supplementary Table 17) diluted in staining buffer and incubation under the condition described above. All samples were analyzed by flow cytometer CytoFLEX S (Beckman Coulter, Brea, CA, USA), acquisition software Cytexpert (Beckman Coulter) and data analysis software Kaluza ver 2.1 (Beckman Coulter).

### Deep sequencing analysis of Ab-coding transcripts

An NGS analysis was performed using the unbiased TCR/BCR repertoire analysis technology developed by Repertoire Genesis (Osaka, Japan) according to the same procedure from the previous study^7^. Sequencing was performed using the Illumina MiSeq paired-end platform (2 × 300 bp). The human PBMC total RNA (Takara Bio USA, San Jose, CA, USA) used in this study was derived from normal human peripheral leukocytes pooled from 426 male/female Asians aged 18–54 years old.

All paired-end reads were classified by index sequences. Sequence assignment was conducted by determining the most identical sequence in a dataset of reference sequences from the international ImMunoGeneTics information system (IMGT) database (https://www.imgt.org). Sequence data were automatically processed, assigned, and combined using repertoire analysis software originally developed by DNA Chip Research (Tokyo, Japan).

### Circos analysis

Circos software was adopted in this study for its high data-to-ink ratio and for its ability to clearly display relational data. Circos open-source software was obtained from https://circos.ca. The V(D)J region recombination data were reformatted to conform to the data file requirements using R, the statistical programming language.

### Detection of somatic hypermutations

The definition of the clone lineage referred to a set of B cells that were related by descent, which arose from the same V(D)J rearrangement event. The definition of the clonotype refers to a single Ab sequence (CDR1, 2, and 3-joined unique sequence)^34^. NGS reads with identical CDR1-2-3 sequences were grouped into a single clonotype. Mutations were detected at all nucleotide position (approximately 315 nucleotides) by comparison with germline sequences and were calculated as a percentage. The mutation rate at each position in the same clone lineage was combined as a clone linage mutation rate. In addition, different lengths of Ab variable regions in annotated reads existed; therefore, mutated positions were standardized by converting the average variable region length of the clonal lineage as 100. The index was similar to the amino acid position in the variable region and clone lineages could be compared at the same magnitude.

### Expected value of somatic mutation

The expected value (*E*) of somatic mutation is calculated using the following equation (1) and (2):

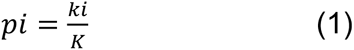

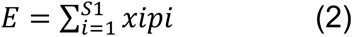

where *K* is the total number of the sequence reads, *ki* is the number of *i*th clone lineage reads. *S1* is the species number of clone lineage reads, and *xi* is the average number of SHM frequency in clone lineage. The expected value (*E*) indicates the expected number of SHM inserted per antibody sequence.

### Diversity index

To estimate BCR diversity in deep sequence data, the Shannon-Weaver index (H′) was calculated using the following equation (3):

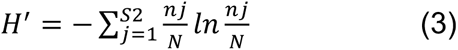

where N is the total number of sequence reads, *nj* is the number of *j*th annotated reads, and S2 is the species number of annotated reads^35^. A greater H′ value reflects greater sample diversity.

### Phylogenetic analysis of the human Ab repertoire

Phylogenetic trees (circular dendrograms) were created by alignment of CDRH3 and CDRL3 amino acid sequences using the multiple sequence alignment program and the Neighbour-Joining method^36^. Furthermore, two phylograms of unimmunized and OVA-immunized TC-mAb mice were assembled within one phylogram based on their amino acid sequences. Additionally, copies comprising the same CDRH3 sequences were counted and overlaid on the leaves of circular dendrograms and are shown with a maximum of 50 reads; thereby, as the number of reads increased, the circle in the leaves increased.

### Hybridoma generation

Immunized TC-mAb rats were euthanized and their spleens were harvested, homogenized to single cell suspensions, and fused with myeloma P3X63Ag8.653 cells using an electro-cell-fusion generator (ECFG21) (Nepagene, Chiba, Japan). Fused hybridoma cells were seeded in 96-well plates. After approximately 14 days of culture, a primary screen of supernatants was performed by an ELISA for detecting wells in which clones that produced OVA-specific Abs. Cells in the positive wells were picked up and passaged in 96-well plates and the supernatant was again analysed by the ELISA. Hybridoma cells were established by two or more limiting dilutions. The subclasses of obtained human mAbs were determined using the Iso-Gold™ Rapid Human Antibody Isotyping Kit (BioAssay Works, Ijamsville, MD, USA) according to the manufacturer’s instructions.

### ELISA

The antibody titters were measured using ELISA method. In brief, 96-well immunoassay plates (Nunc Maxisorp, Thermo Fisher Scientific) were coated with 100 µl/well of antigen overnight and blocked with PBS containing 5% skim milk (Difco Laboratories, Detroit, MI, USA) for 30 min at room temperature. The antigen-coated plates were incubated with 100 µl of samples for 1 h at room temperature. The plates were incubated with 100 µl of goat anti-human IgG (H+L) cross-adsorbed secondary Ab (Abcam, Cambridge, UK) at 1/10,000 dilution in TBS-T for 30 min at room temperature. The plates were washed after each step.

The colorimetric reaction was performed using 100 µl of *o*-phenylenediamine dihydrochloride and stopped using 25 µl of 1 M H_2_SO_4_. After developing for 15 min, the absorbance was read at 492 nm.

### Determination of epitope distribution

The distribution of epitope to which the hybridoma cells bind was analysed by competitive assay. In brief, unlabelled purified Abs were used as capture Abs and biotinylated Abs were used as detector Abs. The 96-well immunoassay plates (Nunc Maxisorp) were coated by each capture Ab at 100 ng/well for 1 hr blocked with PBS containing 5% skim milk (Difco) for 30 min at room temperature. Biotinylating of Ab was performed using EZ-link NHS Biotin kit (Thermo Fisher Scientific) according to the manufacturer’s instructions. After washing, 100 µl biotinylated antibody samples were added to wells and incubated for 1 hr at room temperature. The plates were washed again and incubated with 100 µl of Streptavidiin-HRP conjugate (Proteintech, Tokyo, JP) at 1/3,000 dilution in TBS-T for 30 min at room temperature. The plates were washed once again and developed using *o*-phenylenediamine dihydrochloride, with the reaction being stopped using 1 M H_2_SO_4_. After developing, absorbance was read at 492 nm.

### Humanness score of mAbs

The amino acid sequence of obtained Abs was analysed using the T20 scoring method, which was developed to calculate the humanness of mAb variable region sequences^22^. A Blast search of the variable region was performed against the T20 Cutoff Human Database available at http://abanalyzer.lakepharma.com. The T20 score for an Ab is based on the average of percent identities to the top 20 matched human sequences. To be considered non- or low immunogenic, T20 scores must be above 79 for the FR and CDR sequences and 86 for the FR sequences only.

### Data availability

Source data are provided with this article. The repertoire analysis data generated in this study have been deposited in the DDBJ database under accession code PRJDB18074. SAMD00787159 (TCK052hG_S25), SAMD00787160 (TCK052hK_S29), and SAMD00787152 (TCK052hM_S24) are the data of OVA-immunized TC-mAb rat individual #1. SAMD00787153 (TCK053hG_S27), SAMD00787154 (TCK053hK_S30), and SAMD00787155 (TCK053hM_S26) are the data of OVA-immunized TC-mAb rat individual #2. SAMD00787156 (TCK051hG_S23), SAMD00787157 (TCK051hK_S28), and SAMD00787158 (TCK051hM_S22) are the data of unimmunized TC-mAb rats (pooled).

## Acknowledgements

We thank Y. Sato (DNA Chip Research Inc.) for assistance with generating Circos plots, frequency analyses of VH and VK, and circular dendrograms. We thank Y. Sumida, E. Kaneda, A. Ashiba, M. Morimura, K. Yoshida, M. Fukino, F. Adachi, and T. Kurosaki at Tottori University for assistance with generating and maintaining TC-mAb rats. We thank Mr H. Sugihara of the Technical Department, Tottori University, for his technical support. We also thank Dr. H. Kugoh, Dr. T. Ohbayashi, Dr. Y Nakayama, Dr. T. Ohira, and Dr. Y. Hiramuki at Tottori University. This study was supported in part by the Japan Agency for Medical Research and Development (AMED) under Grant Number JP18am0301009 (Y.K.), JP21am0101124 (Y.K.), JP24ama121046 (Y.K.), JP24gm1610006 (Y.K. and K.T.), JP24gm0010010 (Y.K. and K.T.), JP23am0401002 (Y.K. and K.T.), JP24gm1810008 (Y.K. and Y.B.), Joint Research of the Exploratory Research Center on Life and Living Systems (ExCELLS) (ExCELLS program No. 21-101) (Y.K.), JST CREST Grant Number JPMJCR18S4, Japan (Y.K. and K.T.) and JSPS KAKENHI Grant Number JP 24K11652 (H.S.). This research was partly performed at the Tottori Bio Frontier managed by Tottori prefecture.

## Author contribution statement

H.S., S.A., T.M., and Y.K. planned the study; S.A., A.O., K.K., M. Hirabayashi, and K.N. performed cell culture and rat production experiments; S.T. and T.Y. constructed and provided TALEN vectors.; H.S., S.A., T.M., K.K., and M. Hiratsuka performed animal analyses; S.H., Y.B., and K.T. contributed to analyses and discussion of the data; H.S., S.A., T.M. and Y.K. analysed the results; H.S., S.A., T.M., and Y.K. wrote the manuscript with contributions from each author; K.T. supervised the study.

## Competing interest statement

The authors declare no conflicts of interest.

## Data availability statement

All data are included in the Supplemental Information or available from the authors upon reasonable requests as are unique reagents used in this study.

## Notes

### Competing Interest Statement

The authors have declared no competing interest.

