## Supplementary Figures and Tables for "Comparative analysis of trans-chromosomic rodent models reveals improved somatic hypermutation and class-switch recombination in rats"

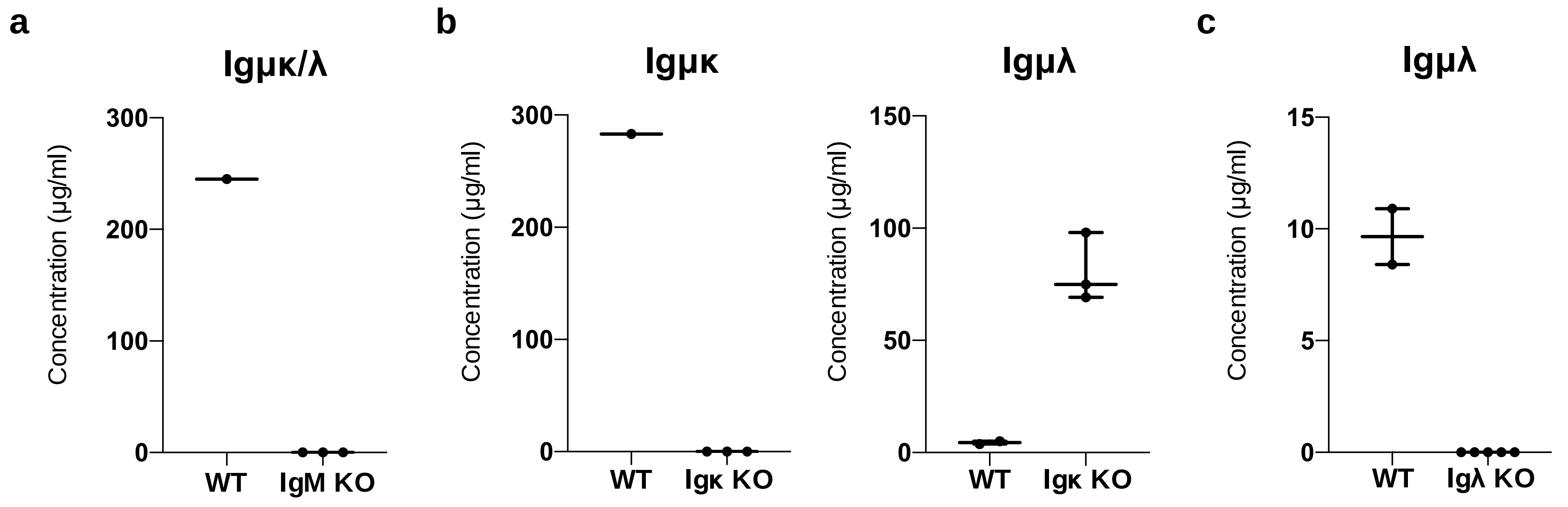

**Supplementary Figure 1| Serum Abs in WT and Ig genes KO rats.**
**(a)** Serum Igμκ/λ concentration in IgM KO and WT rats. **(b)** Serum Igμκ and Igμλ concentrations in WT and Igκ KO rats. **(c)** Serum Igμλ concentration in WT and Igλ KO rats. These rats were crossed to produce Ig genes KO (HKLD) rats, which express no functional protein from IgM, Igκ, and Igλ genes.

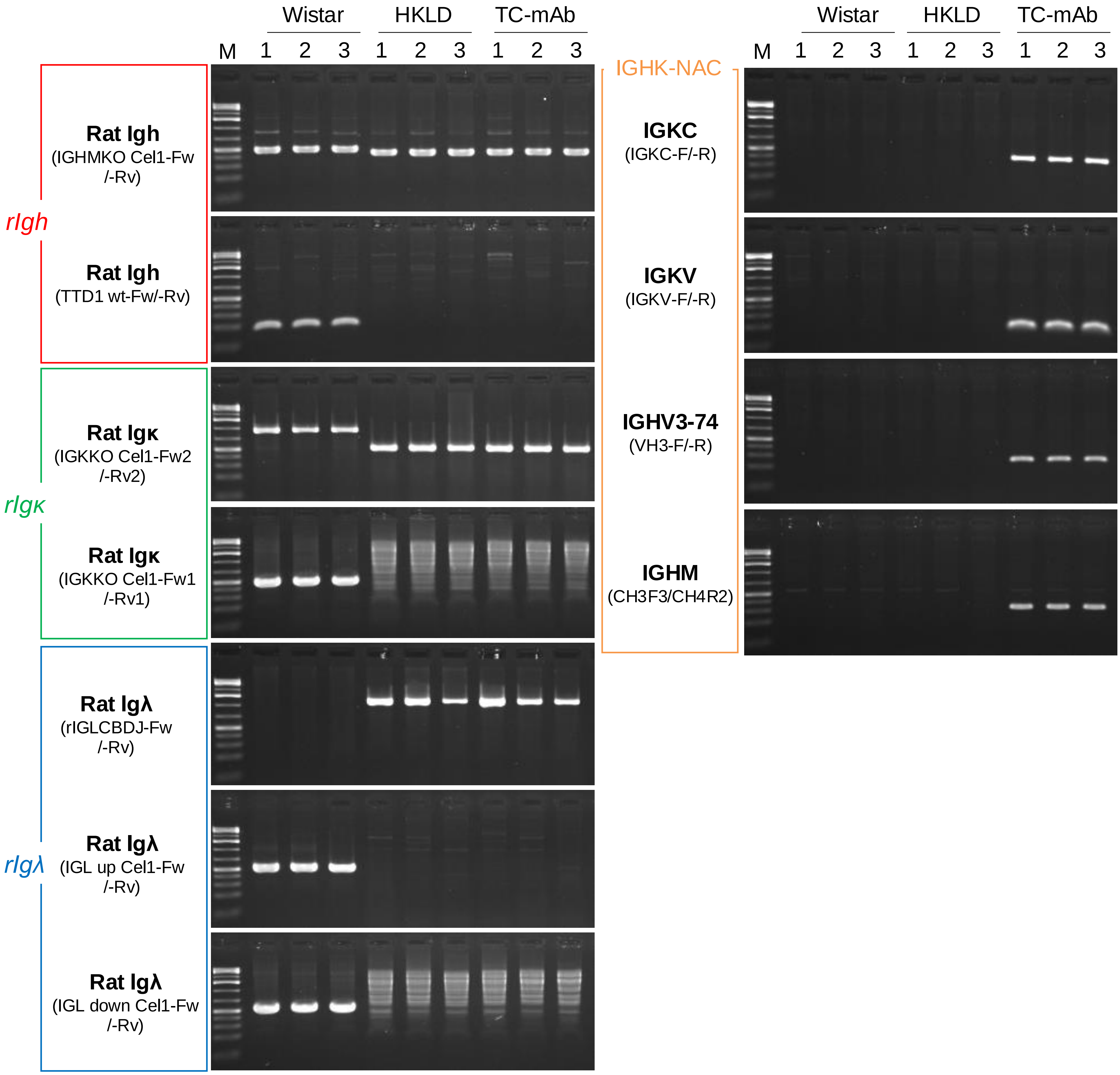

**Supplementary Figure 2| Genotyping results of KO and TC-mAb rats.**
Each gene KO and the presence of IGHK-NAC were confirmed by genomic PCR. The amplified gene fragments and primer sets used are indicated on the left side of each panel. Primers are listed in Supplementary Table 18. Original gel images are shown in Supplementary Figure 27.

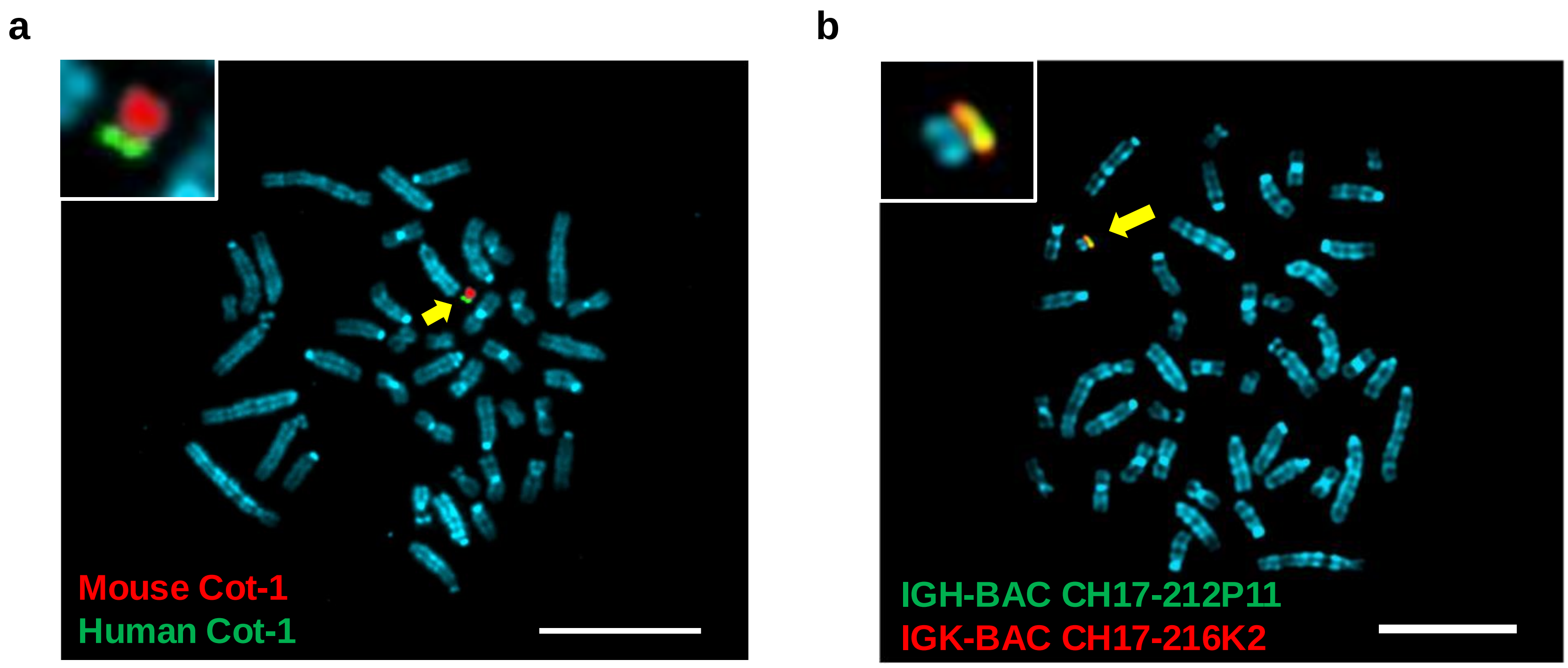

**Supplementary Figure 3| Independent maintenance of the IGHK-NAC in TC-mAb rat.**
Representative FISH images of lymphocytes in TC-mAb rat. (a) FISH image for mouse Cot-1 and human Cot-1 as probe. (b) FISH image for IGK-BAC and IGH-BAC as probe. The image is original of Fig. 1c. Scale bars indicate 10 μm.

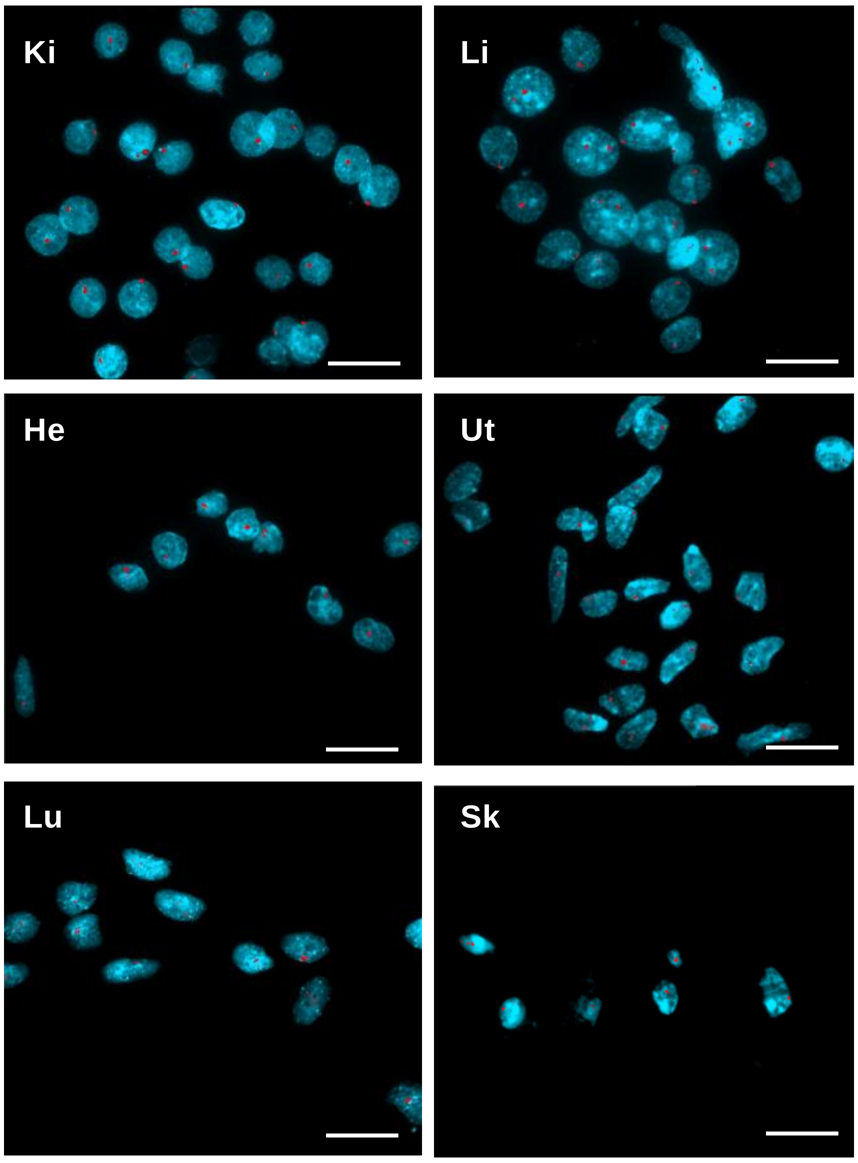

**Supplementary Figure 4| Representative FISH images of each tissue from TC-mAb rat.**
The IGHK-NAC is detected in red by using human Cot-1 DNA as probe. Ki kidney, Li liver, He heart, Ut uterus, Lu lung, Sk skeletal muscle. Scale bars indicate 20μm.

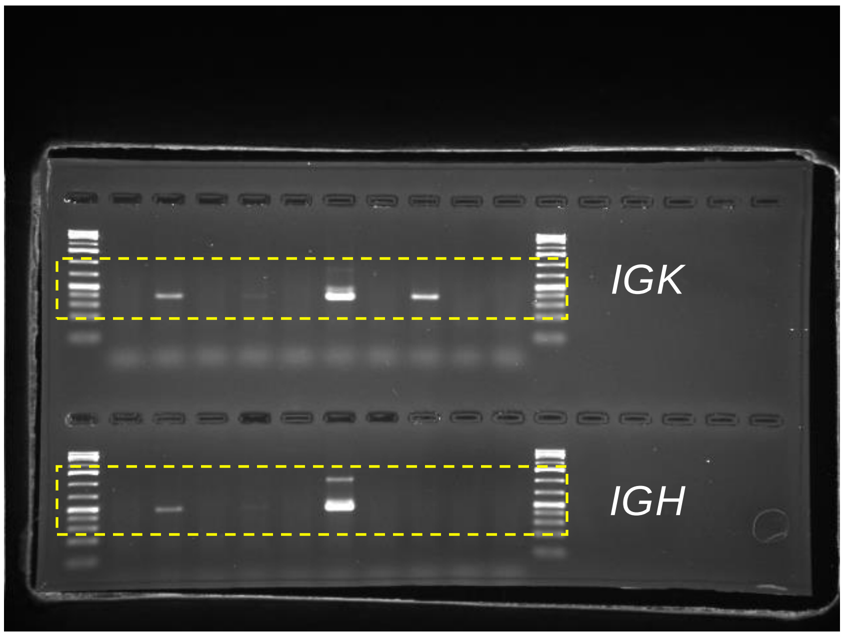

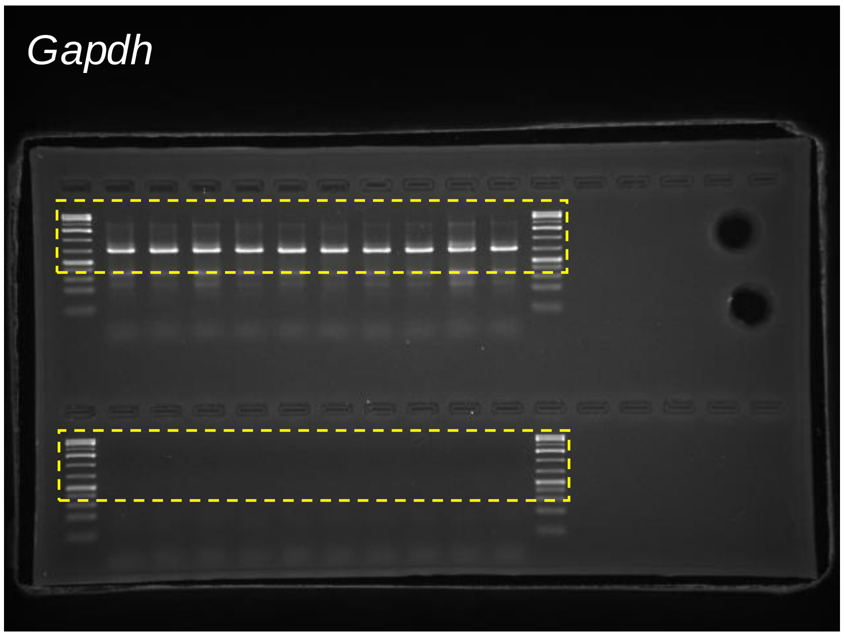

**Supplementary Figure 5| Expression of human genes in various tissues of TC-mAb rat.**
Uncropped images of electrophoresis are presented, and the dashed squares indicate the images used in Figure 2e.

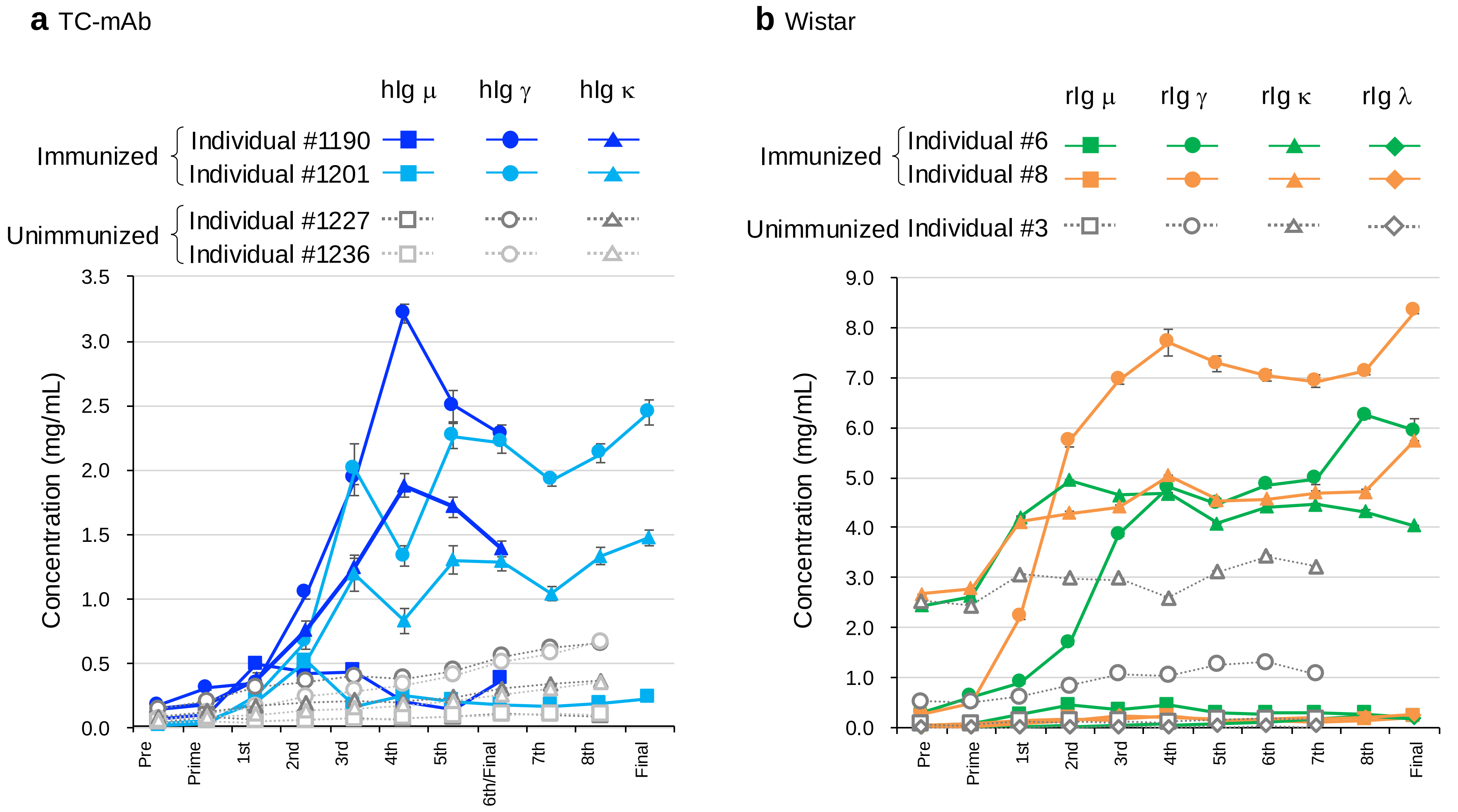

**Supplementary Figure 6| Comparison of Ig concentration between TC-mAb and Wistar rats.**
The serum concentration of Igμ, Igγ, Igκ and Igλ during OVA immunization of TC-mAb **(a)** and Wistar rats **(b)**. The individuals #1201 and #1236 of TC-mAb rats and individuals #6 and #8 of Wistar rats were used in Figure 3. Human Igs are represented as hIgμ (square), hIgγ (round), and hIgκ (triangle) and rats Igs are represented as rIgμ (square), rIgγ (round), rIgκ (triangle), and rIgλ (diamond). Filled symbols and solid lines indicate immunized concentrations; unfilled symbols and dashed lines indicate non-immunized concentrations. Error bars indicate the standard deviation of triplicate measurements.

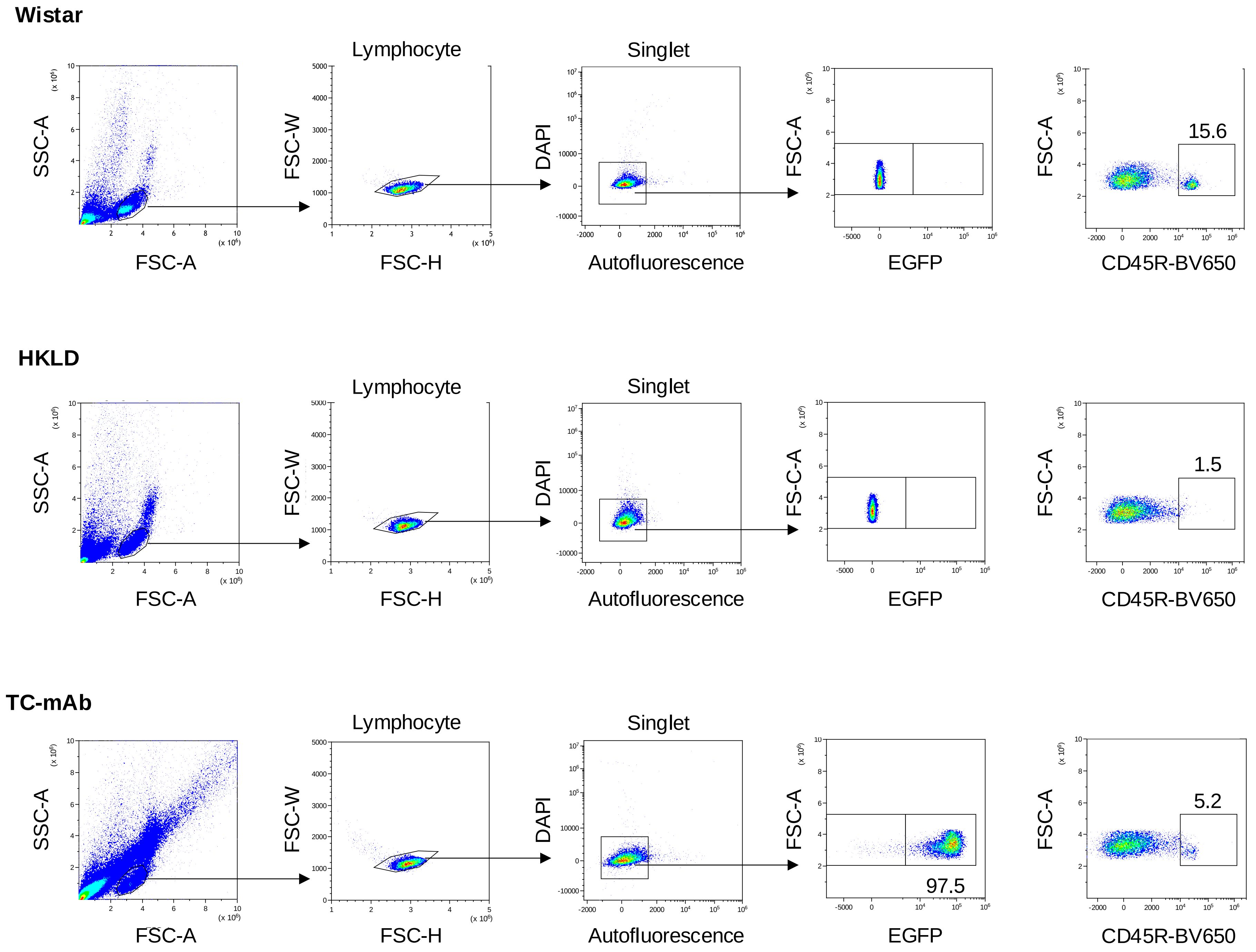

**Supplementary Figure 7| Flow cytometry analysis of B cells ($CD45R^+$) in PBMCs.**
FCM gating strategies for B cells ($CD45R^+$) in PBMCs. In the case of TC-mAb rats, since lymphocyte express EGFP derived from the IGHK-NAC, B cells ($CD45R^+$) is fractionated as $EGFP^+CD45R^+$ double positive cells. The distributions were analysed using 4-6 weak-age of Wistar (Wild-type), HKLD (rat Ig-KO), TC-mAb rat.

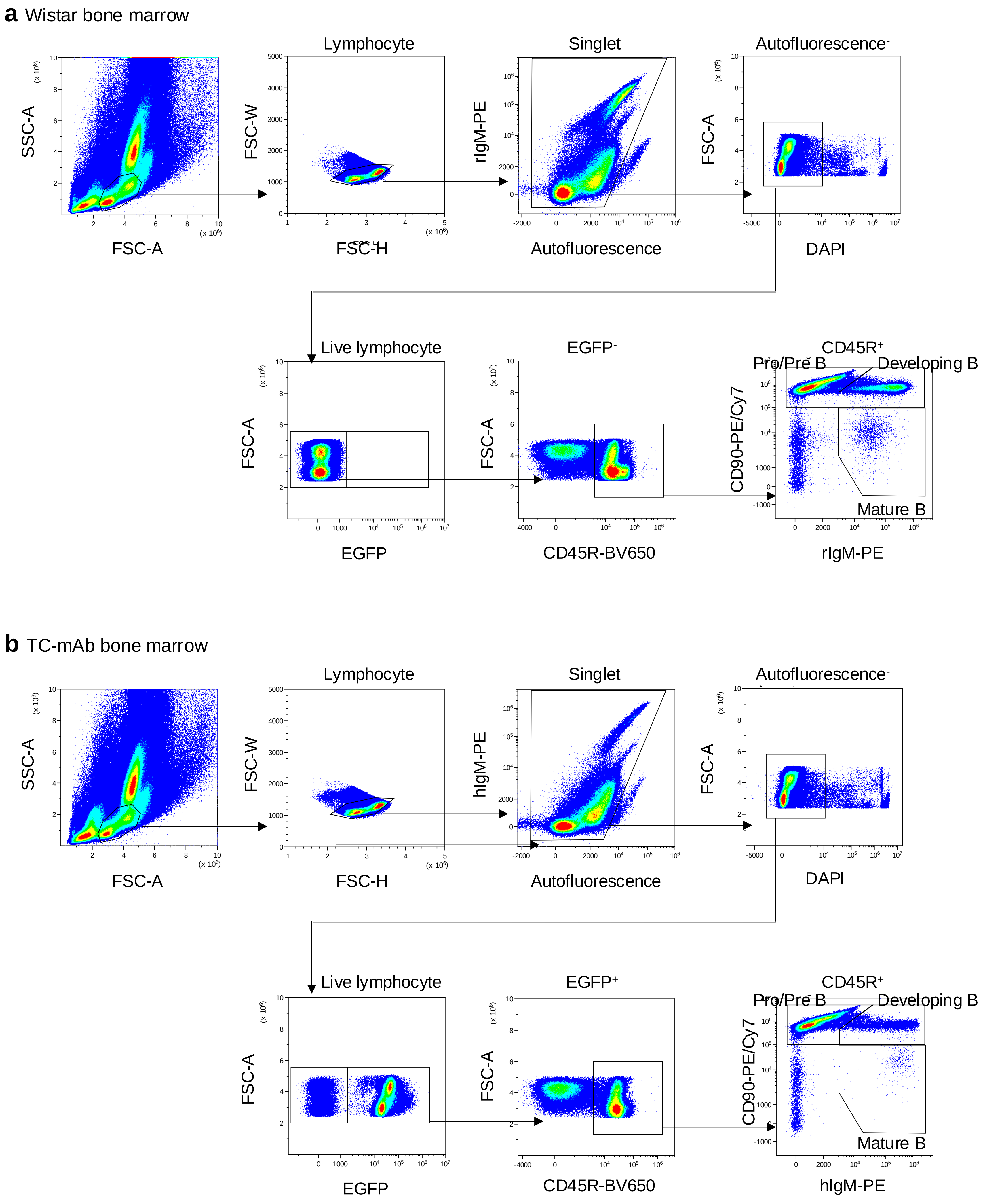

**Supplementary Figure 8| Flow cytometry identification of Pro/Pre B, developing B, and mature B cells in bone marrow.** Flow cytometry gating strategies for Pro/Pre B, developing B, mature B cells in bone marrow. The distributions were analysed using 10-12 weak-age of Wistar **(a)** and TC-mAb **(b)** rats.

**a** Wistar bone marrow

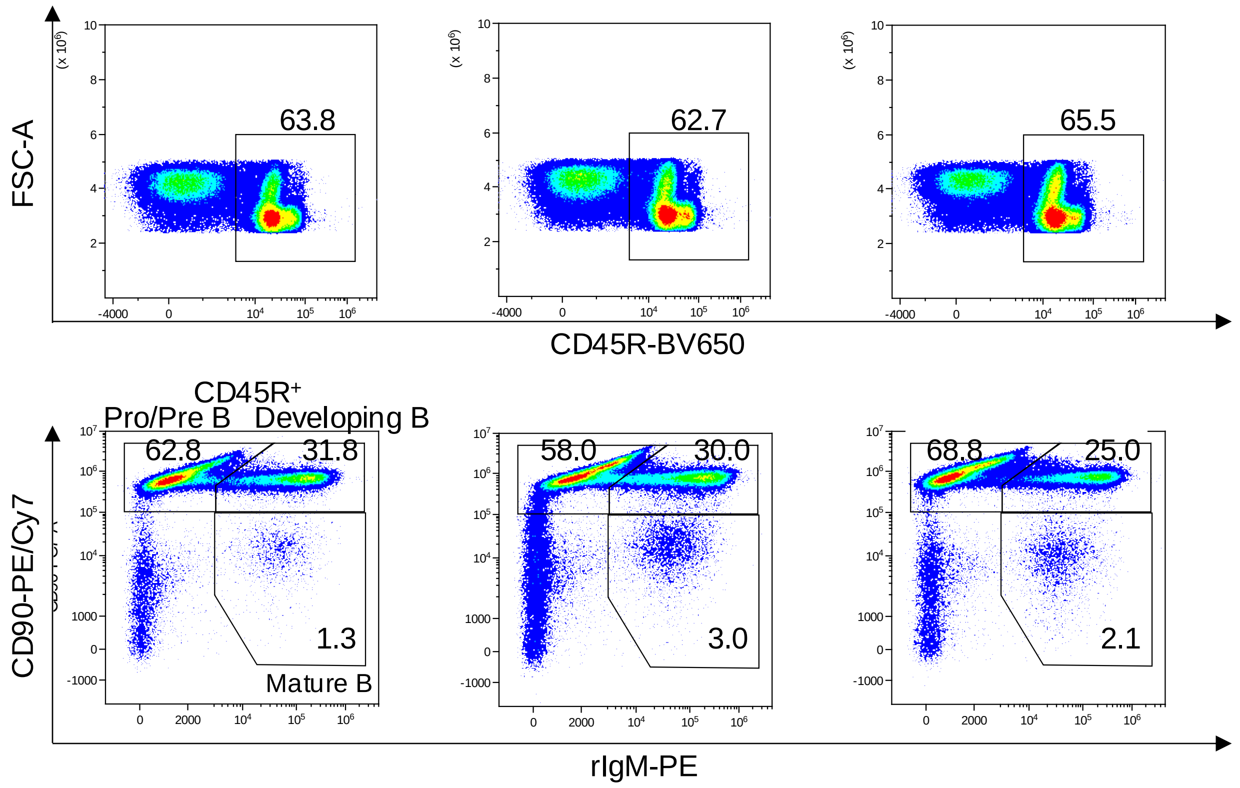

**b** TC-mAb bone marrow

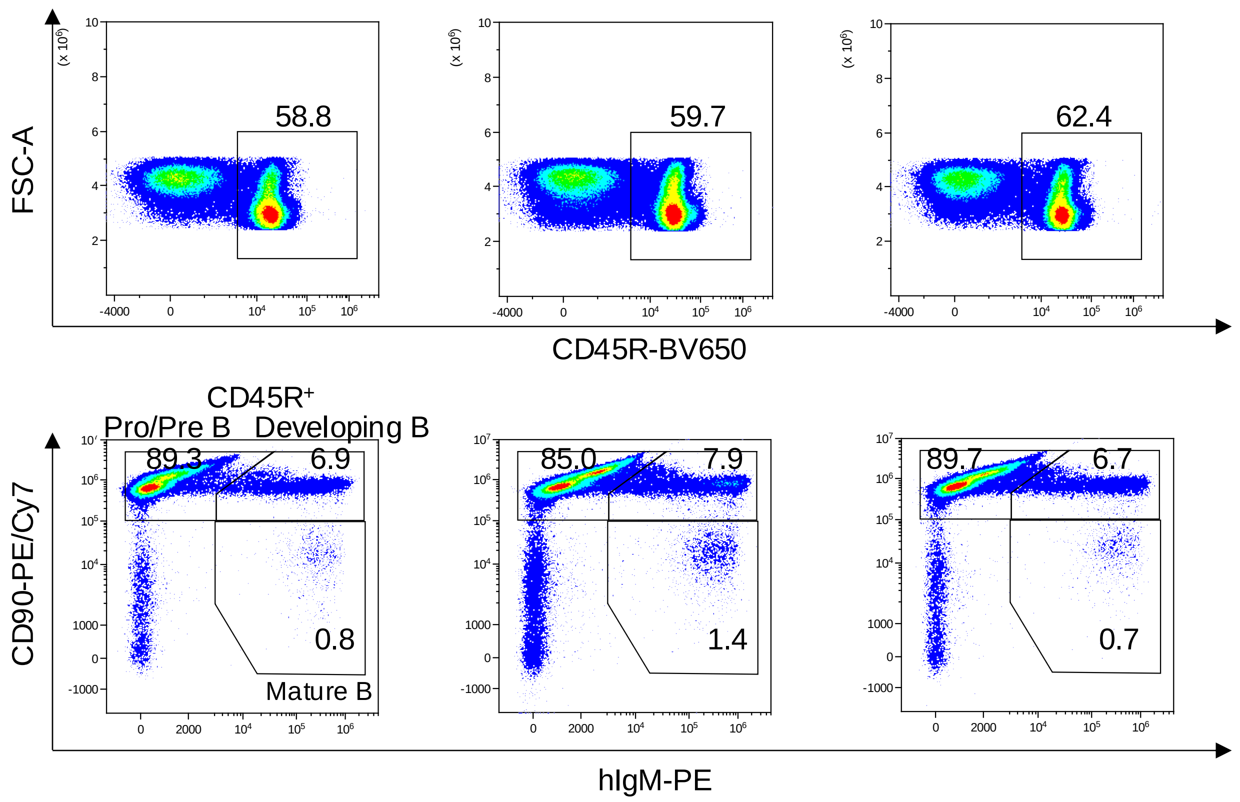

**Supplementary Figure 9| Flow cytometry identification of Pro/Pre B, developing B, and mature B cells in bone marrow.** The compartments of Pro/Pre B (CD45R+CD90+IgM-), developing B (CD45R+CD90+IgM+), mature B (CD45R+CD90-IgM+) cells in bone marrow of Wistar rats (n=3) and TC-mAb rats (n=3). Numbers in the flow cytometry results indicate the percentage of each B cell subset(s). The data collected from Wistar rats (**a**, three panels of upper row) and TC-mAb rats (**b**, three panels of lower row) are indicated, respectively.

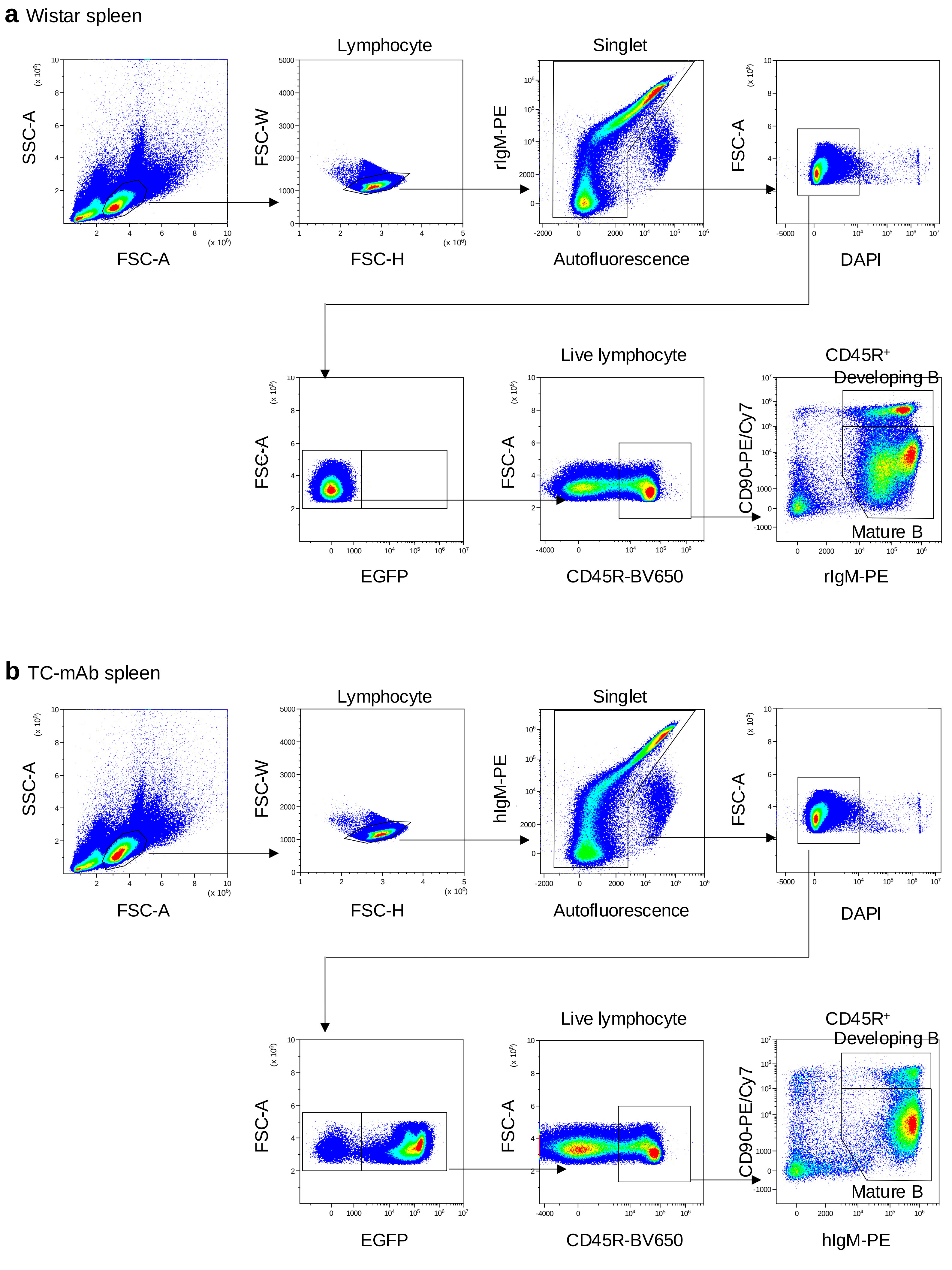

**Supplementary Figure 10| Flow cytometry identification of developing B and mature B cells in spleen.**
Flow cytometry gating strategies for developing B and mature B cells in spleen. The distributions were analysed using 10-12 weak-age of Wistar **(a)** and TC-mAb **(b)** rats.

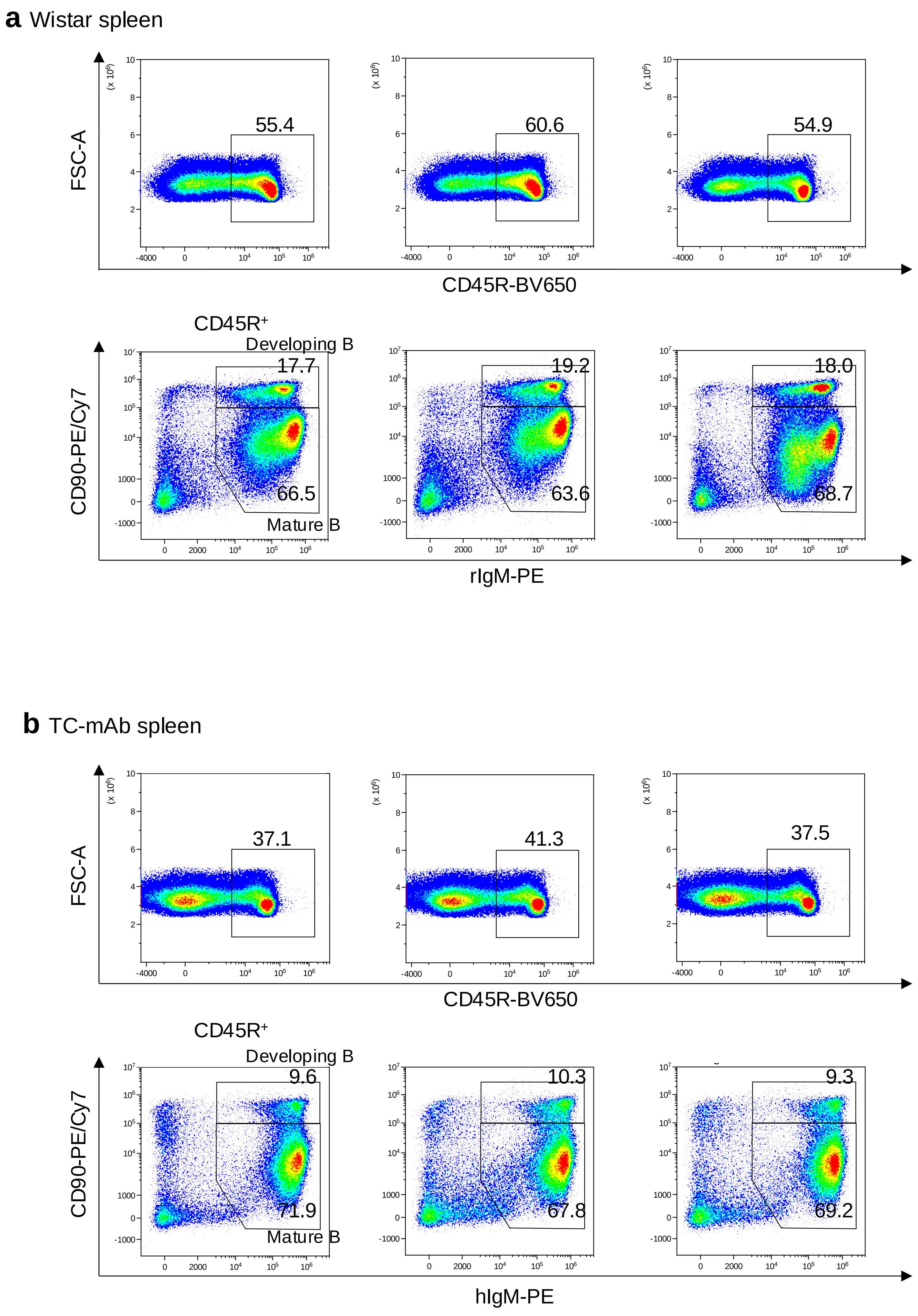

**Supplementary Figure 11| Flow cytometry identification of developing B and mature B cells in spleen.**
The fraction of developing B ($CD45R^+CD90^+IgM^+$) and mature B ($CD45R^+CD90^-IgM^+$) cells in spleen of Wistar rats (n=3) **(a)** and TC-mAb rats (n=3) **(b)**. Numbers in the flow cytometry results indicate the percentage of each B cell subsets.

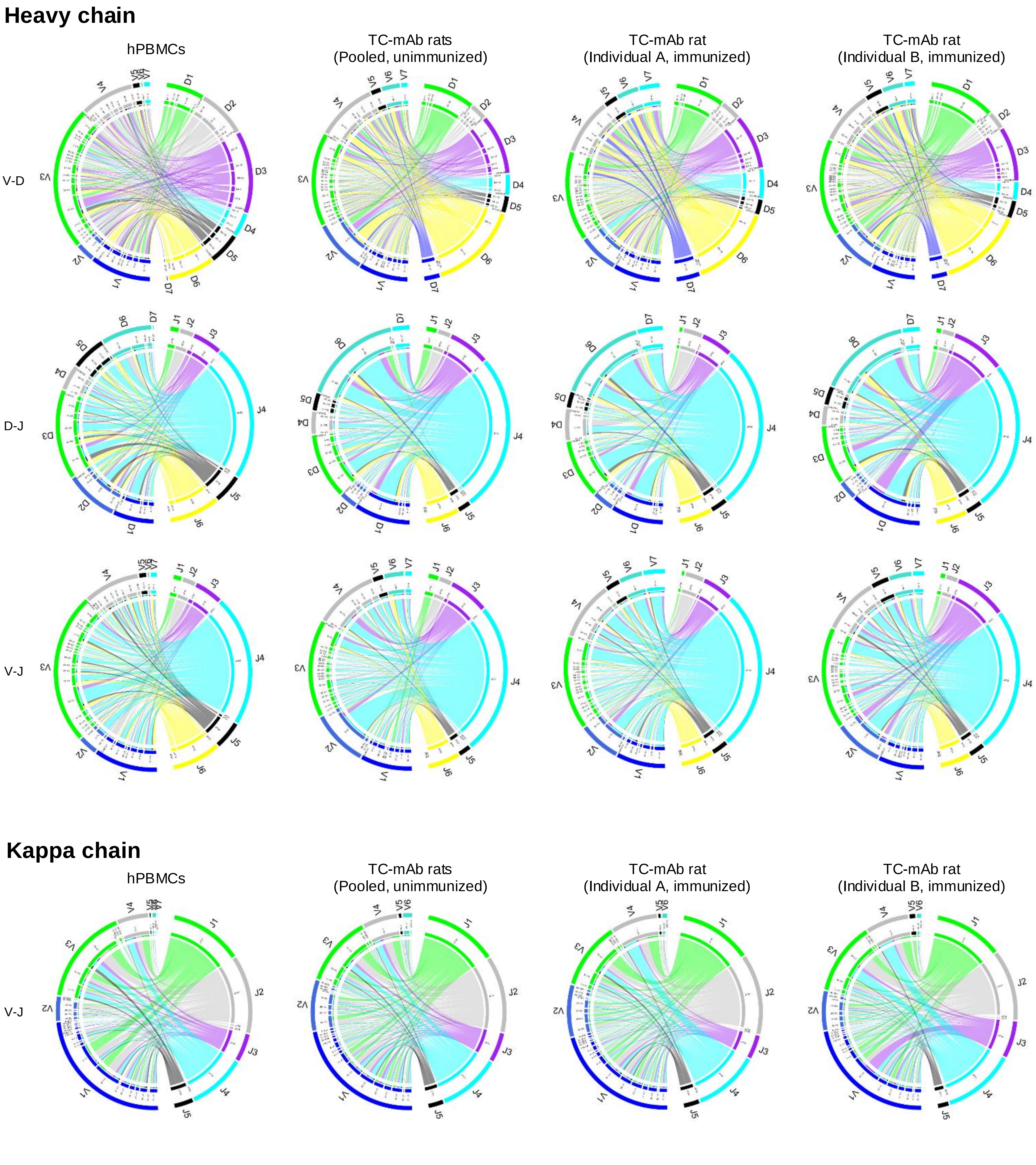

**Supplementary Figure 12| Comparison of V(D)J gene association in TC-mAb rats.**

The circos plots were represented to compare of V(D)J genes detailed association. Outermost tracks mark the boundaries of each V, D, or J region subfamily in the circos plot. Internal tracks indicate the relative frequencies of subgroups within subfamilies. Highlighted links indicate combinations that constitute 1% or more of all sequences observed. Links indicate the relative frequencies of specific V-D, D-J, and V-J combinations of heavy chain and V-J combinations of light chain. The wider links indicate higher frequencies of recombination. The circos plots of hPBMCs, unimmunized TC-mAb rats, and OVA-immunized TC-mAb rats (Individual A and B) were represented.

**a** DH segment usage

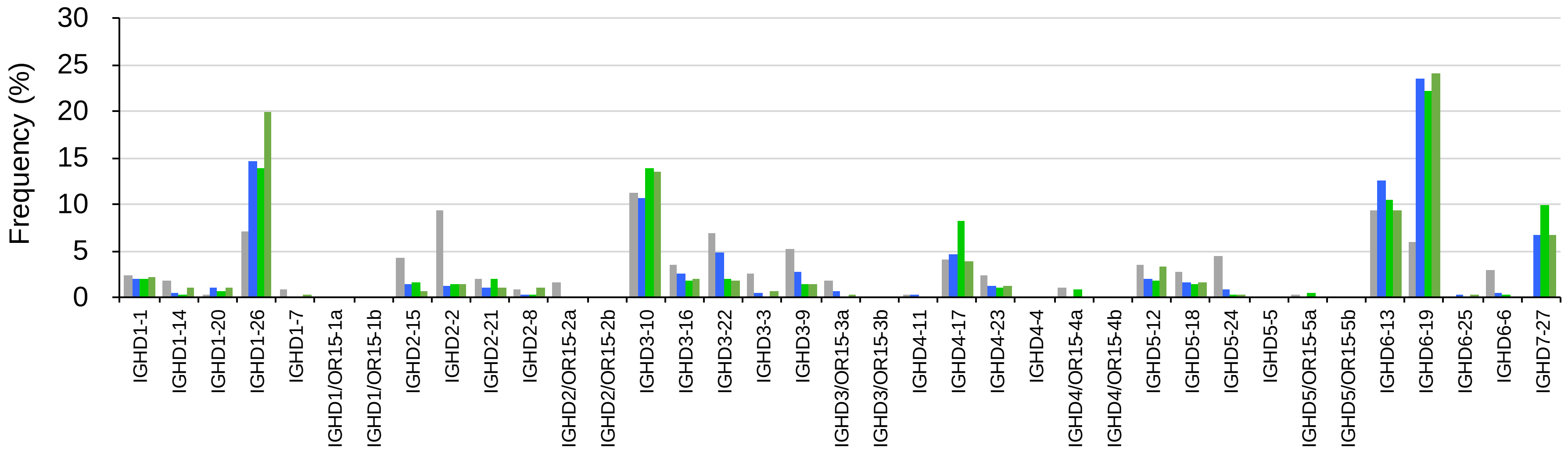

**b** JH segment usage

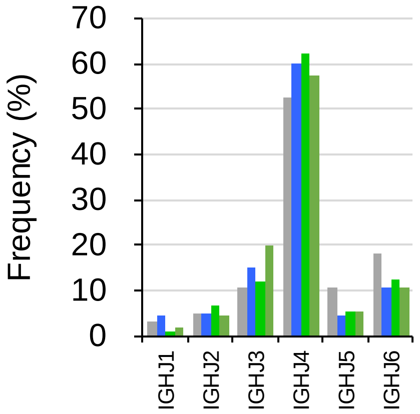

**c**

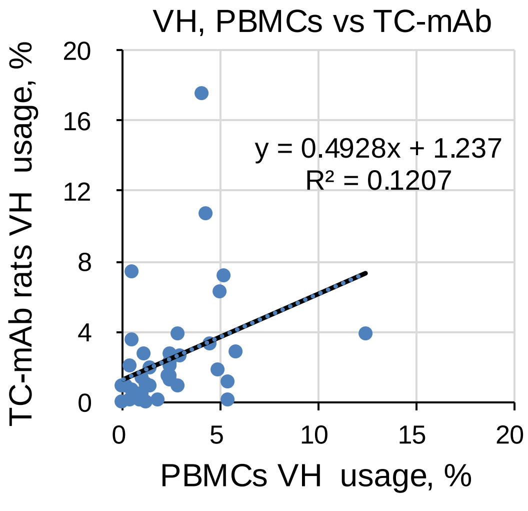

**d**

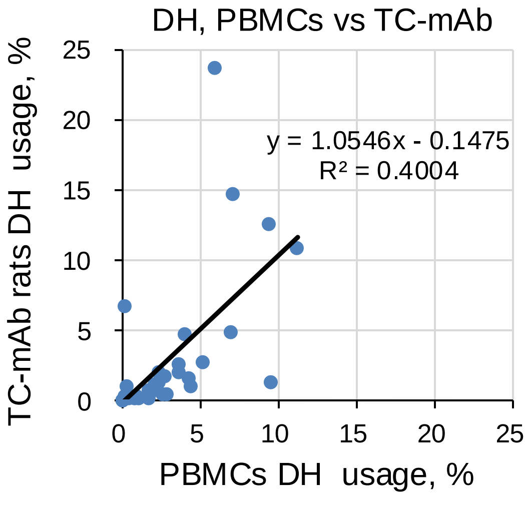

**e**

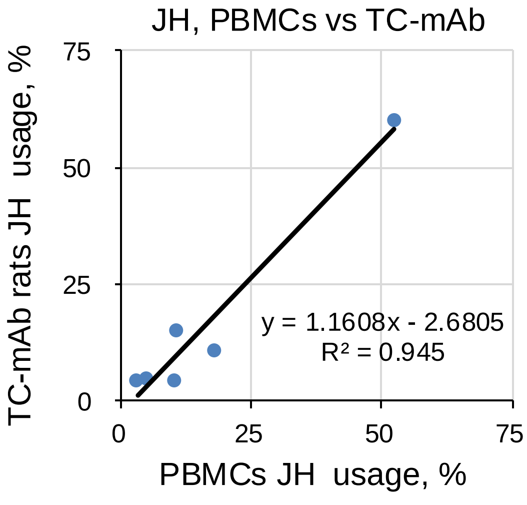

**Supplementary Figure 13| Comparison with the D and J usage between hPBMCs and TC-mAb rats.**
The frequency use of DH **(a)** and JH **(b)** compared between hPBMCs and unimmunized and OVA-immunized TC-mAb rats. The frequency of VH **(c)**, DH **(d)**, and JH **(e)** element is plotted and correlated between unimmunized TC-mAb rats (X-axis) and hPBMCs (Y-axis). R2 values are exhibited as correlation coefficient.

a VH

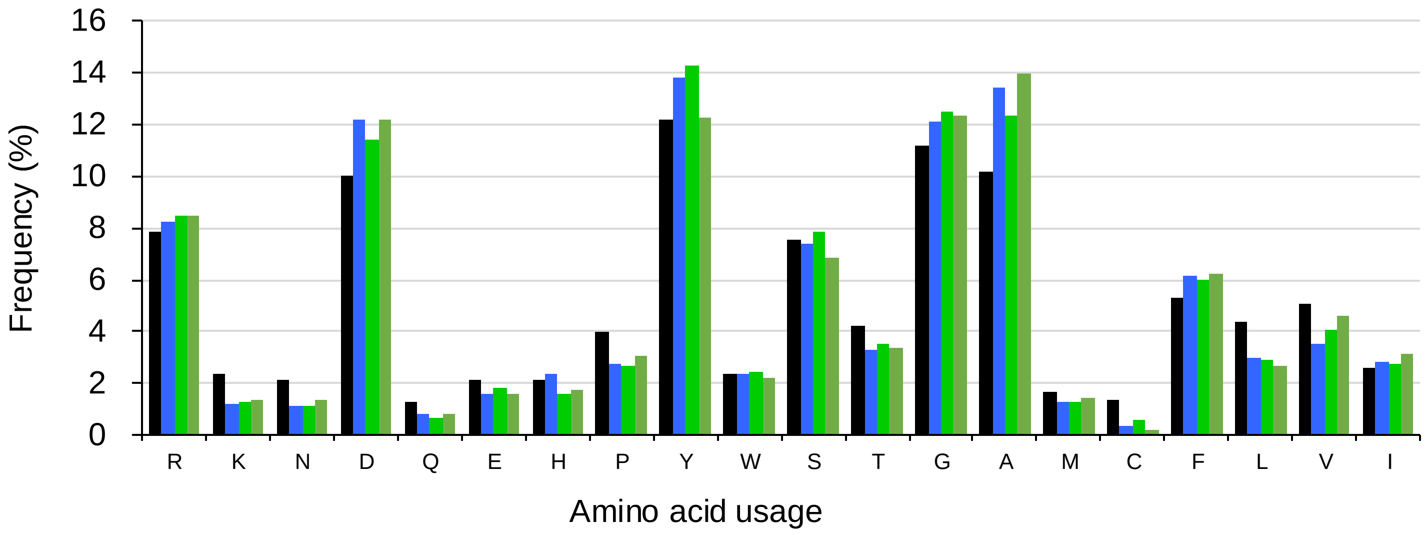

b VK

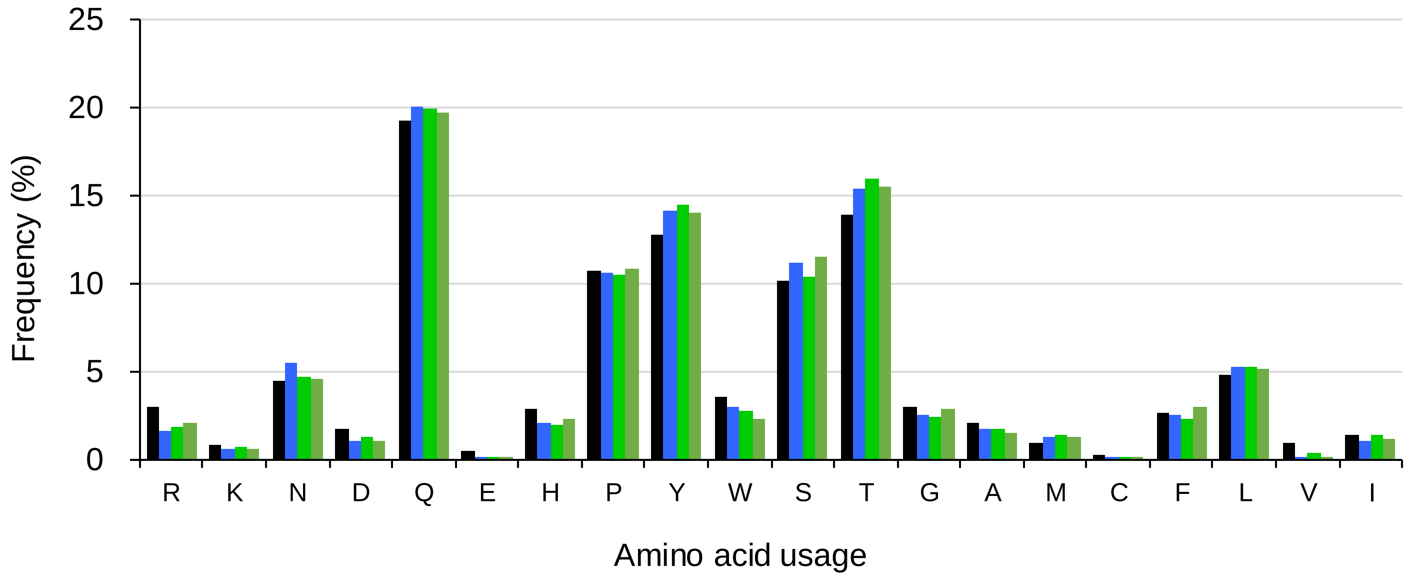

**Supplementary Figure 14| Amino acid usage frequency in TC-mAb rats.**
Amino acid usage within the CDRH3 (a) and CDRL3 (b) of productive rearrangements in healthy human donors (grey), TC-mAb rats with (light and dark green), and without immunization (blue) are represented.

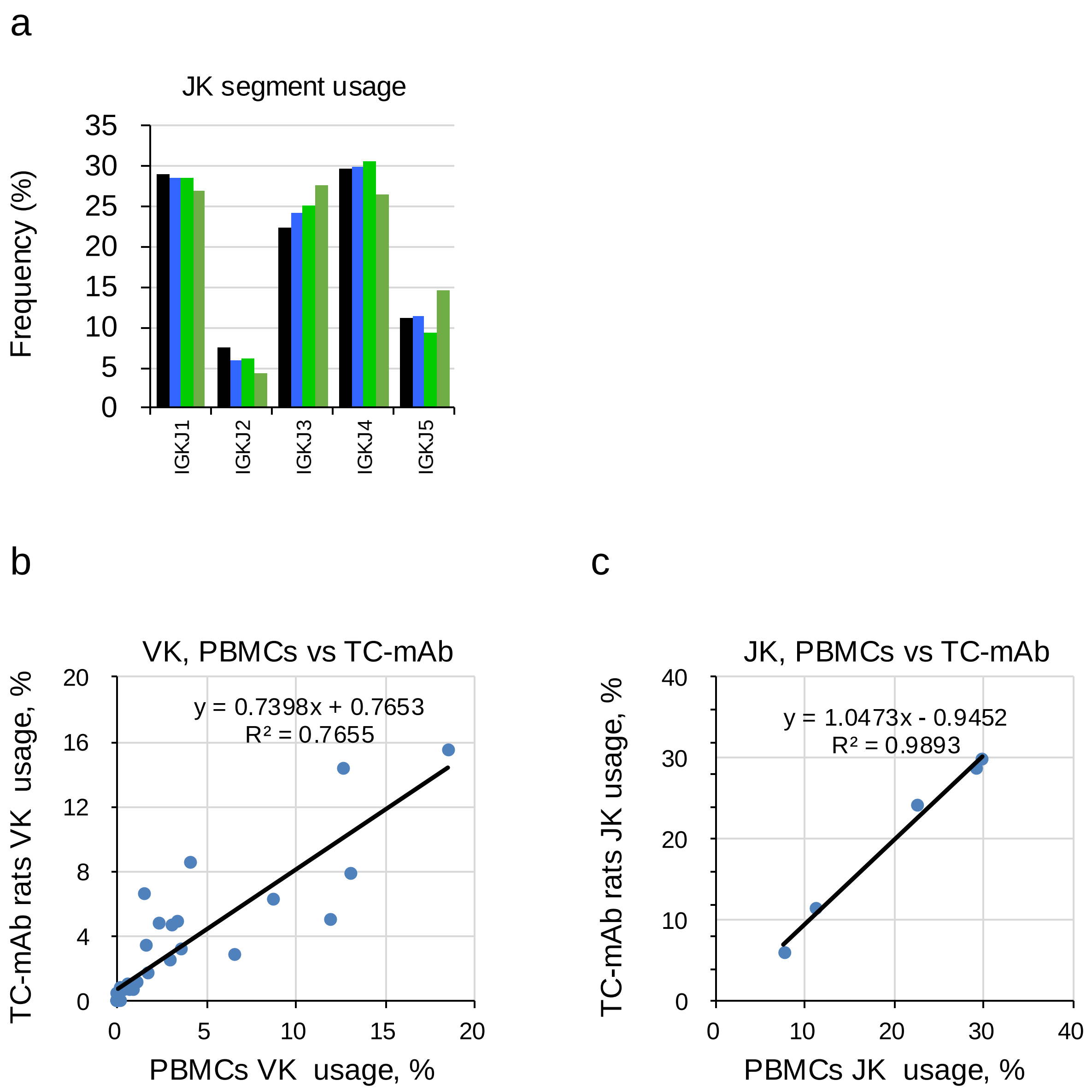

**Supplementary Figure 15| Comparison with the J usage between hPBMCs and TC-mAb rats.**
**(a)** The frequency use of JK compared between hPBMCs and unimmunized and OVA-immunized TC-mAb rats. The frequency of VK **(b)** and JK **(c)** element is plotted and correlated between unimmunized TC-mAb rats (X-axis) and hPBMCs (Y-axis). R2 values are exhibited as correlation coefficient.

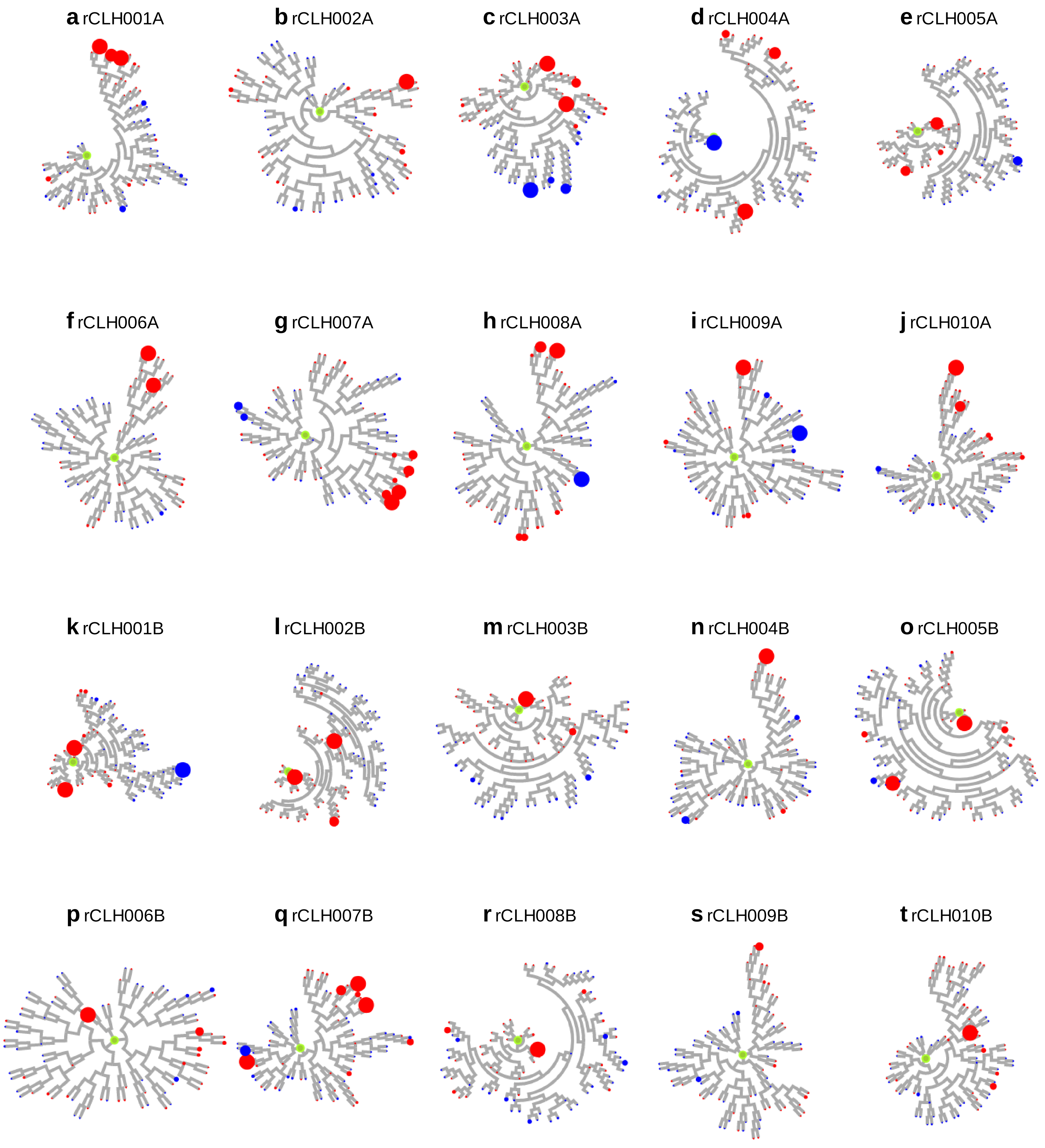

**Supplementary Figure 16| Cricular dendrogram and expanded clonotypes.**
The phylogenetic trees of the CDRH3 and expanded clonotypes were represented. The circular dendrograms of top 10 frequently used clone lineages were over-rayed with the number of copies having the same CDR3 sequences in VH. The results of rCLH001A-010A of heavy chain from individual A **(a-j)** and rCLH001B-010B of heavy chain from individual B **(k-t)** were represented. The leaves of circular dendrograms with a maximum of 50 reads, thereby the larger number drew the larger circle in the leaves. The color of the circule was indicated with (red) or without (blue) immunization in TC-mAb rats.

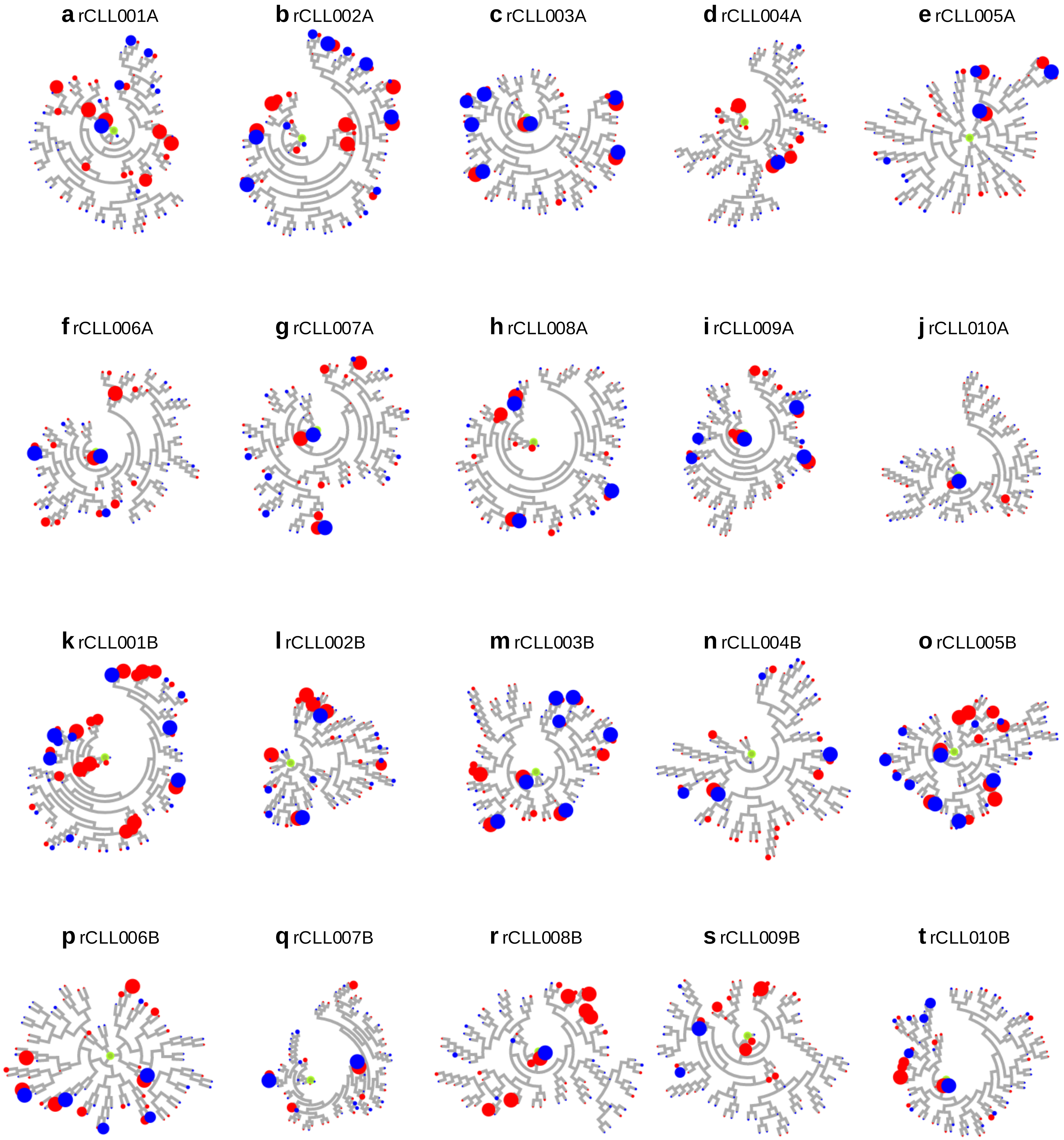

**Supplementary Figure 17| Cricular dendrogram and expanded clonotypes.**
The phylogenetic trees of the CDRL3 and expanded clonotypes were represented. The circular dendrograms of top 10 frequently used clone lineages were over-rayed with the number of copies having the same CDR3 sequences in VK. The results of rCLL001A-010A of heavy chain from individual A **(a-j)** and rCLL001B-010B of kappa chain from individual B **(k-t)** were represented. The leaves of circular dendrograms with a maximum of 50 reads, thereby the larger number drew the larger circle in the leaves. The color of the circule was indicated with (red) or without (blue) immunization in TC-mAb rats.

**a** TC-mAb rats

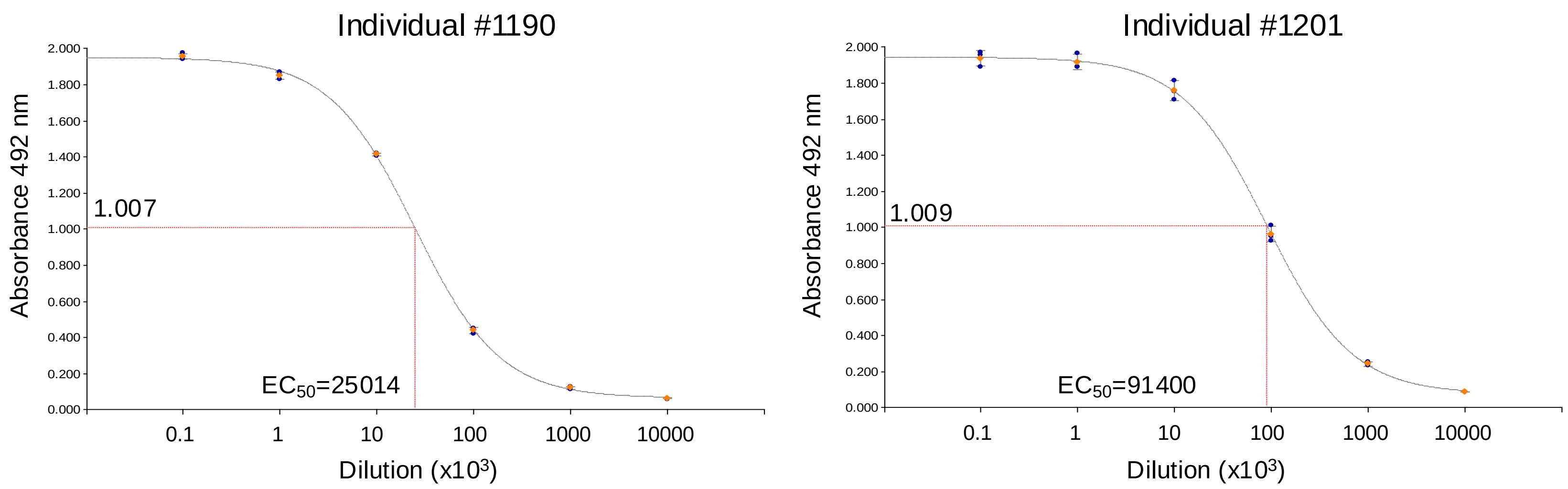

**b** Wistar rats

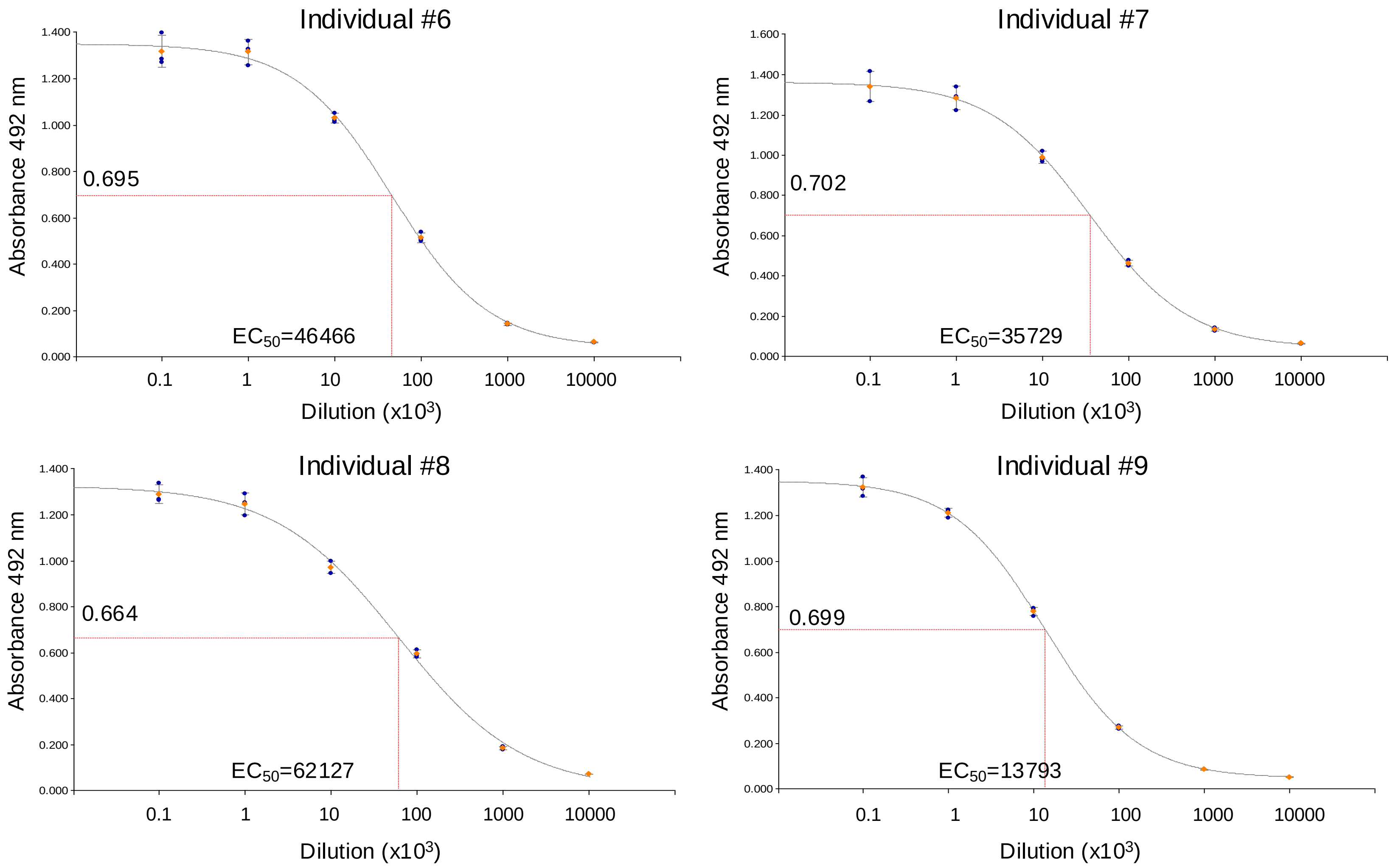

**c**

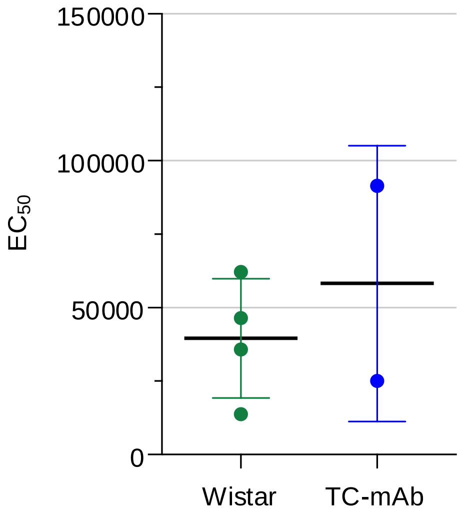

| Curve name | Y (Absorbance 492 nm) | X (Dilution of Anti-sera) |
|---|---|---|
| TC-mAb rat, individual #1190 | 1.007 | 25014 |
| TC-mAb rat, individual #1201 | 1.009 | 91400 |
| Wistar, individual #6 | 0.695 | 46466 |
| Wistar, individual #7 | 0.702 | 35729 |
| Wistar, individual #8 | 0.664 | 62127 |
| Wistar, individual #9 | 0.699 | 13793 |

**Supplementary Figure 18| Comparison of antisera affinities.**
The effective concentration 50 (EC$_{50}$) of anti-sera obtained from two individual TC-mAb **(a)** and four individual Wistar rats **(b)** are represented. The titres of serially diluted OVA-specific anti-sera indicated that the EC$_{50}$ of respective anti-sera was calculated at a dilution of 1.3−15.4 in $10^4$ by a 4-parameter logistic curve fitting model. The table is a summary of each absorbance and Ab concentration at EC$_{50}$. **(c)** Beeswarm plot of EC$_{50}$ for TC-mAb and Wistar rats. Error bars indicate the standard deviation of measurements.

#### **a** Variable region sequences of the obtained mAbs

| Clone | | Variable region | SHMs | Top V gene match | Top D gene match | Top J gene match |
|---|---|---|---|---|---|---|
| #001 | H chain | QVQLQESGPGLVKPSETLSLTCTVSGGSISSNYWSWIRQPPGKGLEWIGYIYYSGTTNYNPSLKSRVTISVDTSKNQFSLKLSSVTAADTAVYYCARERSGSYLFDYWGQGTLVTVSS | 3 | IGHV4-59*01 | IGHD1-26*01 | IGHJ4*02 |
| | κ chain | DIRMTQSPSSLSASVGDRVTITCRASQGISNSLAWYQQRPGKSPNLLLSAASRLGTGVPSRFSGSGSGTDYTLTISSLQPEDFATYYCQQYSSTPLTFGGGTKVEIK | 10 | IGKV1-NL1*01 | - | IGKJ4*01 |
| #006 | H chain | EVQLVESGGGLVKPGGSLRLSCAASGFTFSNAWMNWVRQPPGKGLEWVGRIKRKTDGGTTDYAAPVKGRFTISRDDSKNTLYLQMNSLKTEDTAVYYCTTEGYSYWGQGSLVTVSS | 3 | IGHV3-15*01 | IGHD3-16*02 | IGHJ4*02 |
| | κ chain | DLQMTQSPSAMSASVGDSVTITCRTSQGINNYLAWFQQKPGKVPKRLIYAASSLHSGVPSRFSGSGSGTEFTLTISSLQPEDFATYYCLQHATYPHTFGPGTKVDIK | 8 | IGKV1-17*03 | - | IGKJ3*01 |
| #007 | H chain | QVQLVQSGAEVKKPGASVKVACKASGYTFTSYGISWVRQGPGQGLEWMGWISADNDNTNYAQKFQGRVIMTTDTSTSTAYMELRSLRSDDTAVYYCARDPGIAVIPFDYWGQGTLVAVSS | 9 | IGHV1-18*01 | IGHD6-19*01 | IGHJ4*02 |
| | κ chain | EIVLTQSPGTLSLSPGERATLSCRASQIVSSSYLAWFQQKPGQAPRLLIYGASSRATGIPDRFSGSGSGTDFTLTISRLEPEDFAVYYCQQFVTSPWTFGQGTKVEIK | 7 | IGKV3-20*01 | - | IGKJ1*01 |
| #016 | H chain | EVQLLESGGGLVQPGGSLRLSCAASGFTFSIYAMSWVRQAPGKGLEWVSTISDFGDTTHHADSVKGRFIISRDNSKNTLYLQMNSLRAEDTAVYYCAKPYSSGWPAPLDYWGQGTLVTVSS | 11 | IGHV3-23*01 | IGHD6-19*01 | IGHJ4*02 |
| | κ chain | DIQMTQSPSSLSASVGDRVTITCRASQDTSSSLAWYQQKPGKAPKLLVNAASRLESGVPSRFSGSGSGTDFTLTISSLQPEDFATYYCQRYYTPPYTFGQGTKLEIK | 12 | IGKV1-NL1*01 | - | IGKJ2*01 |
| #023 | H chain | QVQLQQSGPGLVKPSQTLSLTCAISGDSVSRNGAVWNWIRQSPSRGLEWLGRTYYRSKWYNDYAESVKSRIAIKPDTSKNQFSLQLNSVTPEDTAVYYCARDFDYWGQGTLVTVSS | 8 | IGHV6-1*01 | N/A | IGHJ4*02 |
| | κ chain | EIVLTQSPGTLSLSPGERATLSCRVSQSVSTYLAWYQQKPGQAPRLLIYGASNRATGIPDRFSGSGSGTDFTLTISRLEPEDFAVYYCQQYNSSPLTFGGGTKVEIK | 8 | IGKV3-20*01 | - | IGKJ4*01 |

N/A: Sequences not available in the database

#### **b** Mutated positions in the nucleotide sequences

##### H chain variable region

```
001    CAGGTGCAGCTGCAGGAGTCGGGCCCAGGACTGGTGAAGCCTTCGGAGACCCTGTCCCTC
006    GAGGTGCAGCTGGTGGAGTCTGGGGGAGGCTTGGTAAAGCCTGGGGGGTCCCTTAGACTC
007    CAGGTTCAATTGGTGCAGTCTGGAGCTGAGGTGAAGAAGCCTGGGGCCTCAGTGAAGGTC
016    GAGGTGCAGCTGTTGGAGTCTGGGGGAGGCTTGGTACAGCCTGGGGGGTCCCTGAGACTC
023    CAGGTACAGCTGCAGCAGTCAGGTCCAGGACTGGTGAAGCCCTCGCAGACCCTCTCACTC
        **** **  **  * **** **    *  **    ****  *    *  *     **

                          CDR1
001    ACCTGCACTGTCTCT------GGTGGCTCCATCAGTAGTAACTACTGGAGCTGGATCCGG
006    TCCTGTGCAGCCTCT------GGATTCACTTTCAGTAACGCCTGGATGAACTGGGTCCGC
007    GCCTGCAAGGCTTCT------GGTTACACCTTTACCAGCTATGGTATCAGCTGGGTGCGA
016    TCCTGTGCAGCCTCT------GGATTCACCTTTAGCATCTATGCCATGAGCTGGGTCCGC
023    ACCTGTGCCATCTCCGGGGACAGTGTCTCTAGAAATGGTGCTGTTTGGAACTGGATCAGG
        ****        **          *   * *    *         * **** *   *

                                                         CDR2
001    CAGCCCCCAGGGAAGGGACTGGAGTGGATTGGGTATATTTATTACAG---TG------GG
006    CAGCCTCCAGGGAAGGGGCTGGAGTGGGTTGGCCGTATTAAAAGAAAAACTGATGGTGGG
007    CAGGGCCCTGGACAAGGGCTTGAGTGGATGGGATGGATCAGCGC------TGACAATGAT
016    CAGGCTCCAGGGAAGGGGCTGGAGTGGGTCTCAACCATTAGTGA------TTTGGTGAT
023    CAGTCCCCATCGAGAGGCCTTGAGTGGCTGGGAAGGACATACTACAG---GTCCAAGTGG
       ***   **        ** ** ****** *       *

001    ACCACCAACTACAACCCCTCCCTCAAGAGTCGAGTCACCATATCAGTAGACACGTCCAAG
006    ACAACAGACTACGCTGCACCCGTGAAAGGCAGATTCACCATCTCAAGAGATGATTCAAAA
007    AACACAAACTATGCACAGAAATTCCAGGGCAGAGTCATCATGACCACAGACACATCCACG
016    ACCACACACCACGCAGACTCCGTGAAGGGCCGGTTCATCATCTCCAGAGACAATTCCAAG
023    TATAATGATTATGCAGAATCTGTGAAAAGTCGAATAGCCATCAAACCAGACACATCCAAG
           *   *  *          *  *  *  *  *   ***        ***     ** *

001    AACCAGTTCTCCCTGAAGCTGAGCTCTGTGACCGCTGCGGACACGGCCGTGTATTACTGT
006    AACACGCTGTATCTGCAAATGAACAGCCTGAAAACCGAGGACACAGCCGTGTATTACTGT
007    AGCACAGCCTACATGGAGCTGAGGAGCCTGAGATCTGACGACACGGCCGTGTATTACTGT
016    AACACGCTGTATCTGCAAATGAACAGCCTGAGAGCCGAGGACACGGCCGTATATTACTGT
023    AACCAGTTCTCCCTGCAGCTGAATTCTGTGACTCCCGAGGACACGGCTGTGTATTACTGT
       * *      *    ** *  ***       ***    * *  ***** ** ** *********

                     CDR3
001    GCGAGAGAGAGGAGTGGGAGCTAC------CTCTTTGACTACTGGGGCCAGGGAACCCTG
006    ACCA----------CAGAGGGTTA-----------TAGCTACTGGGGCCAGGGATCCCTG
007    GCGAGAGACCCGGGTATAGCAGTG---ATTCCCTTTGACTACTGGGGCCAGGGAACCCTG
016    GCGAAGCCGTATAGCAGTGGCTGGCCCGCCCCTCTTGACTACTGGGGCCAGGGAACCCTG
023    GCAAGAGA-----------------------TTTTGACTACTGGGGCCAGGGAACCCTG
        * *                                  *   ***************** *****

001    GTCACCGTCTCCTCAG
006    GTCACCGTCTCCTCAG
007    GTCGCCGTCTCCTCAG
016    GTCACCGTCTCCTCAG
023    GTCACCGTCTCCTCAG
       *** ************
```

##### κ chain variable region

```
006    GACCTCCAGATGACCCAGTCTCCATCTGCCATGTCTGCATCTGTAGGAGACAGCGTCACC
016    GACATCCAGATGACCCAGTCTCCATCCTCCCTGTCTGCATCTGTAGGAGACAGAGTCACC
001    GACATCCGGATGACCCAGTCTCCATCCTCCCTGTCTGCATCTGTAGGAGACAGAGTCACC
007    GAGATTGTGTTGACGCAGTCTCCAGGCACCCTGTCTTTGTCTCCAGGGGAAAGAGCCACC
023    GAAATTGTGTTGACGCAGTCTCCAGGCACCCTGTCTTTGTCTCCAGGGGAAAGAGCCACC
       **  *   * **** *********    ** *****    ***  *** ** ** * ****

                         CDR1
006    ATCACTTGTCGGACGAGTCAGGGCATTAACAAT---TATTTAGCCTGGTTTCAGCAGAAA
016    ATCACTTGCCGGGCGAGTCAGGACACTAGCAGT---TCTTTAGCCTGGTATCAGCAGAAA
001    ATCACTTGCCGGGCGAGTCAGGGCATTAGCAAT---TCTTTAGCCTGGTATCAGCAGAGA
007    CTCTCCTGCAGGGCCAGTCAGATTGTCAGCAGCAGCTACTTAGCCTGGTTCCAGCAGAAA
023    CTCTCCTGCAGGGTCAGTCAGAGTGTTAGCACC---TACTTAGCCTGGTACCAGCAGAAA
        ** * **  **    ******      *  **     * **********   ******* *

                                       CDR2
006    CCAGGGAAAGTCCCTAAGCGCCTGATCTATGCTGCATCCAGTTTGCACAGTGGGGTCCCA
016    CCAGGCAAAGCCCCAAAGCTCCTGGTCAATGCTGCATCCAGATTGGAAAGTGGGGTCCCA
001    CCAGGGAAATCCCCTAACCTCCTGCTCTCTGCTGCATCCAGATTGGGAACTGGGGTCCCA
007    CCTGGCCAGGCTCCCAGGCTCCTCATCTATGGTGCATCCAGCAGGGCCACTGGCATCCCA
023    CCTGGCCAGGCTCCCAGGCTCCTCATCTATGGTGCATCCAACAGGGCCACTGGCATCCCA
       ** **  *    ** *  * ***  **  ** ********     *    * ***  *****

006    TCAAGGTTCAGCGGCAGTGGATCTGGGACAGAATTCACTCTCACAATCAGCAGCCTGCAG
016    TCCAGGTTCAGTGGCAGTGGATCTGGGACGGATTTCACTCTCACCATCAGCAGCCTGCAG
001    TCCAGGTTCAGTGGCAGTGGGTCTGGGACGGATTACACTCTCACCATCAGCAGCCTGCAG
007    GACAGGTTCAGTGGCAGTGGGTCTGGGACAGACTTCACTCTCACCATCAGCAGACTGGAG
023    GACAGGTTCAGTGGCAGTGGGTCTGGGACAGACTTCACTCTCACCATCAGCAGACTGGAG
          ******** ******** ******** ** * ********* ******** *** **

                                       CDR3
006    CCTGAAGATTTTGCAACTTATTACTGTCTACAGCATGCTACTTACCCTCACACTTTCGGC
016    CCTGAAGATTTTGCAACTTATTACTGTCAACGATATTATACTCCCCCGTACACTTTTGGC
001    CCTGAAGATTTTGCAACTTATTACTGTCAGCAGTATTCTAGTACCCCGCTCACTTTCGGC
007    CCTGAAGATTTTGCAGTGTATTACTGTCAGCAGTTGTTACCTCACCGTGGACGTTCGGC
023    CCTGAAGATTTTGCAGTGTATTACTGTCAGCAGTATAATAGCTCACCGCTCACTTTCGGC
       ***************   **********  *    *  **      **    ** ** ***

006    CCTGGGACCAAAGTGGATATCAAAC--
016    CAGGGGACCAAGCTGGAGATCAAAC--
001    GGTGGGACCAAGGTGGAGATCAAACGA
007    CAAGGGACCAAGGTGGAAATCAAAC--
023    GGAGGGACCAAGGTGGAGATCAAACGA
          ********  **** *******
```

**Supplementary Figure 19| Sequence analysis of mAbs obtained from TC-mAb rats.**
The variable region sequences of the heavy and light chain sequences obtained from mAbs were represented. **(a)** The amino acid sequences, SHM frequencies, and top-matched V(D)J segments of five clones were represented. **(b)** The sequence alignment of the variable region heavy chain (left side) and kappa chain (right side) of five clones were represented. The section flanked by arrows indicates the complementarity-determining region (CDR).

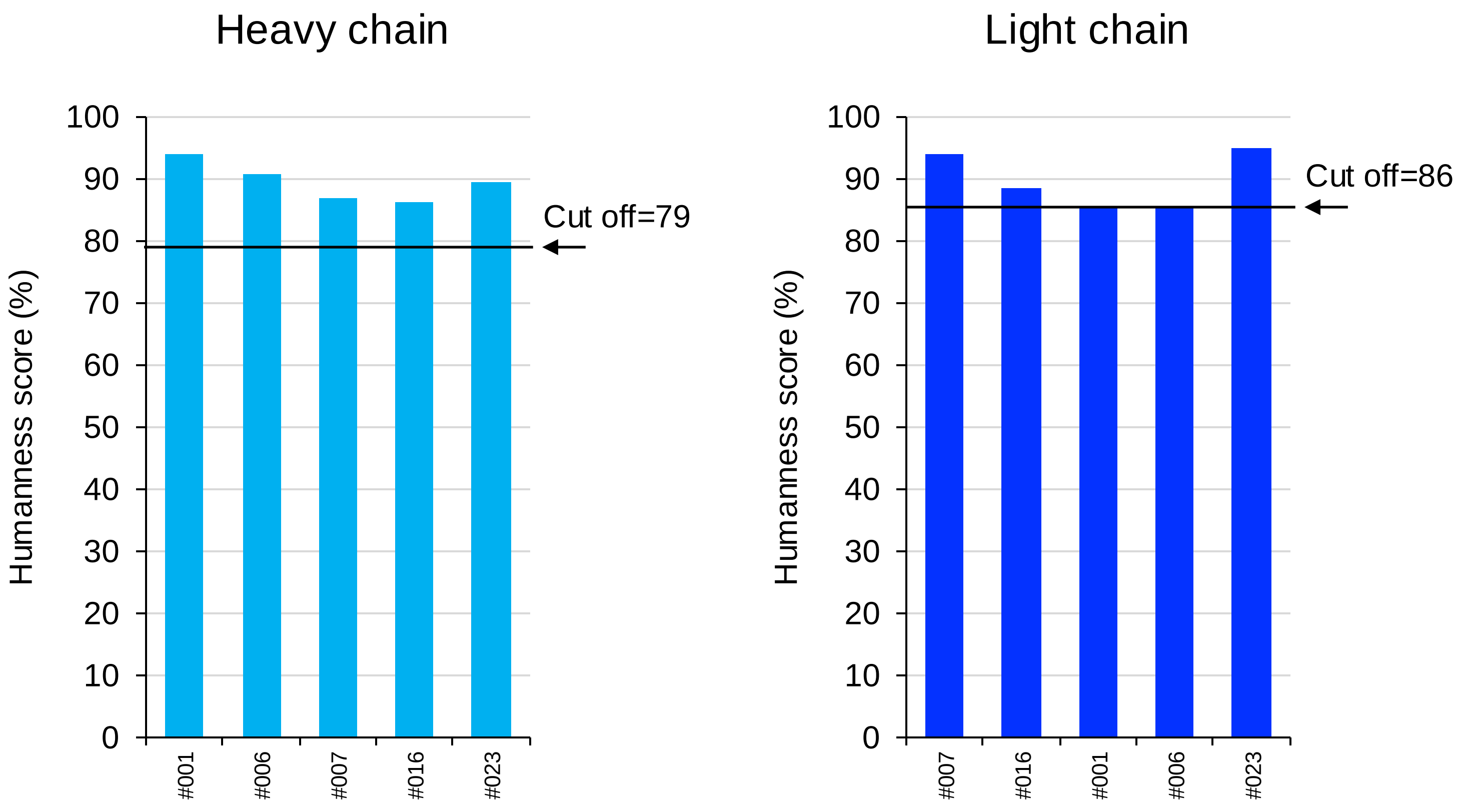

**Supplementary Figure 20| Humanness score of mAbs.**
Humanness score of mAb isolated from TC-mAb rats were represented. The nucleotide sequence of mAbs used is the same as in Supplementary Fig. 19. The T20 scores were determined using the T20 score analyzer. To make a judgement as human sequence, cut off values (Heavy chain was 79 and κ chain was 86) was used according to the threshold set by previous report[23]. The thresholds were indicated as arrows.

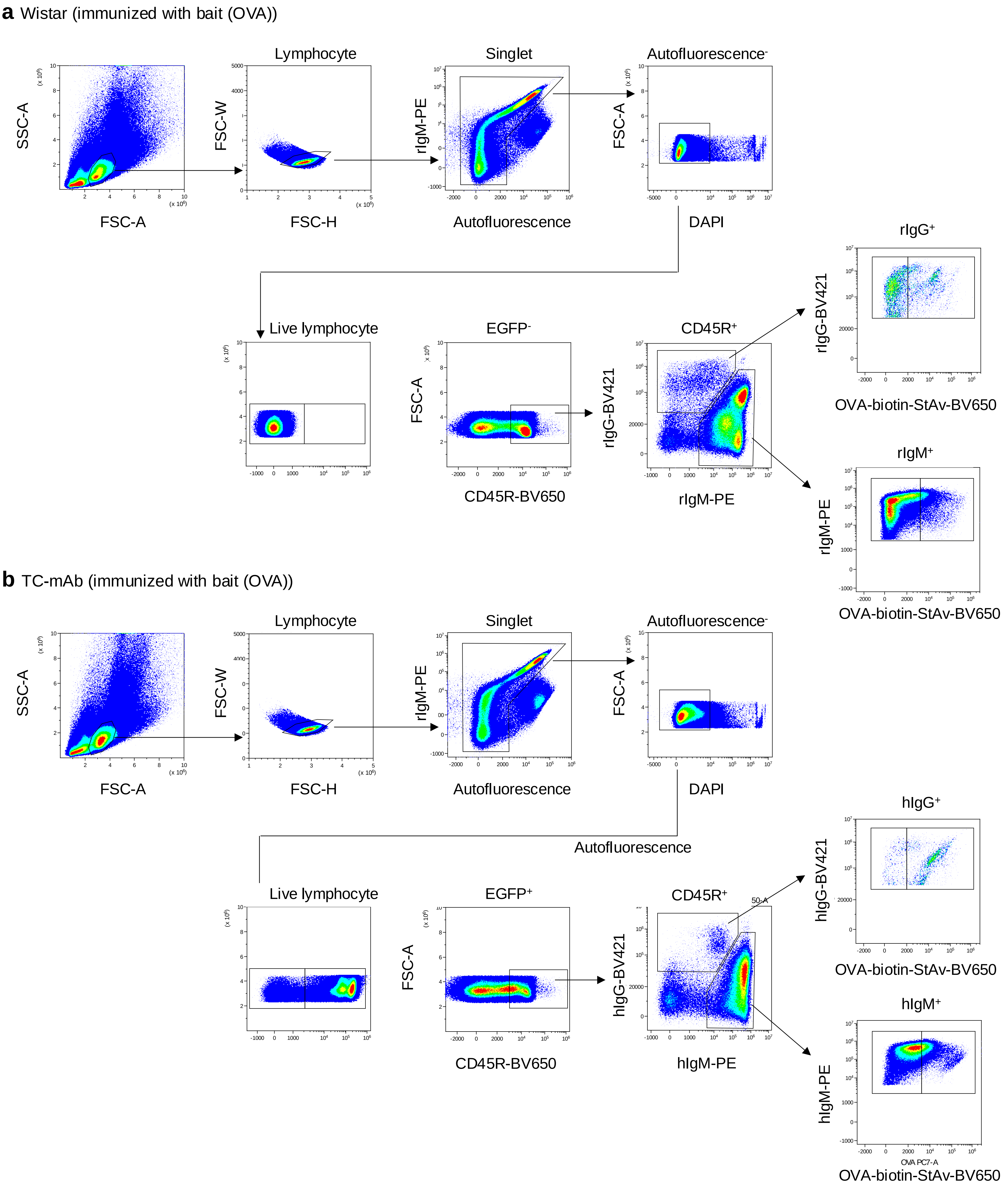

**Supplementary Figure 21| Flow cytometry identification of antigen-specific Ab expressing B cells in spleen.**
Flow cytometry gating strategies for antigen-specific Ab expressing B cells in spleen. The distributions were analysed using OVA-immunized or unimmunized Wistar (**a**) and TC-mAb (**b**) rats.

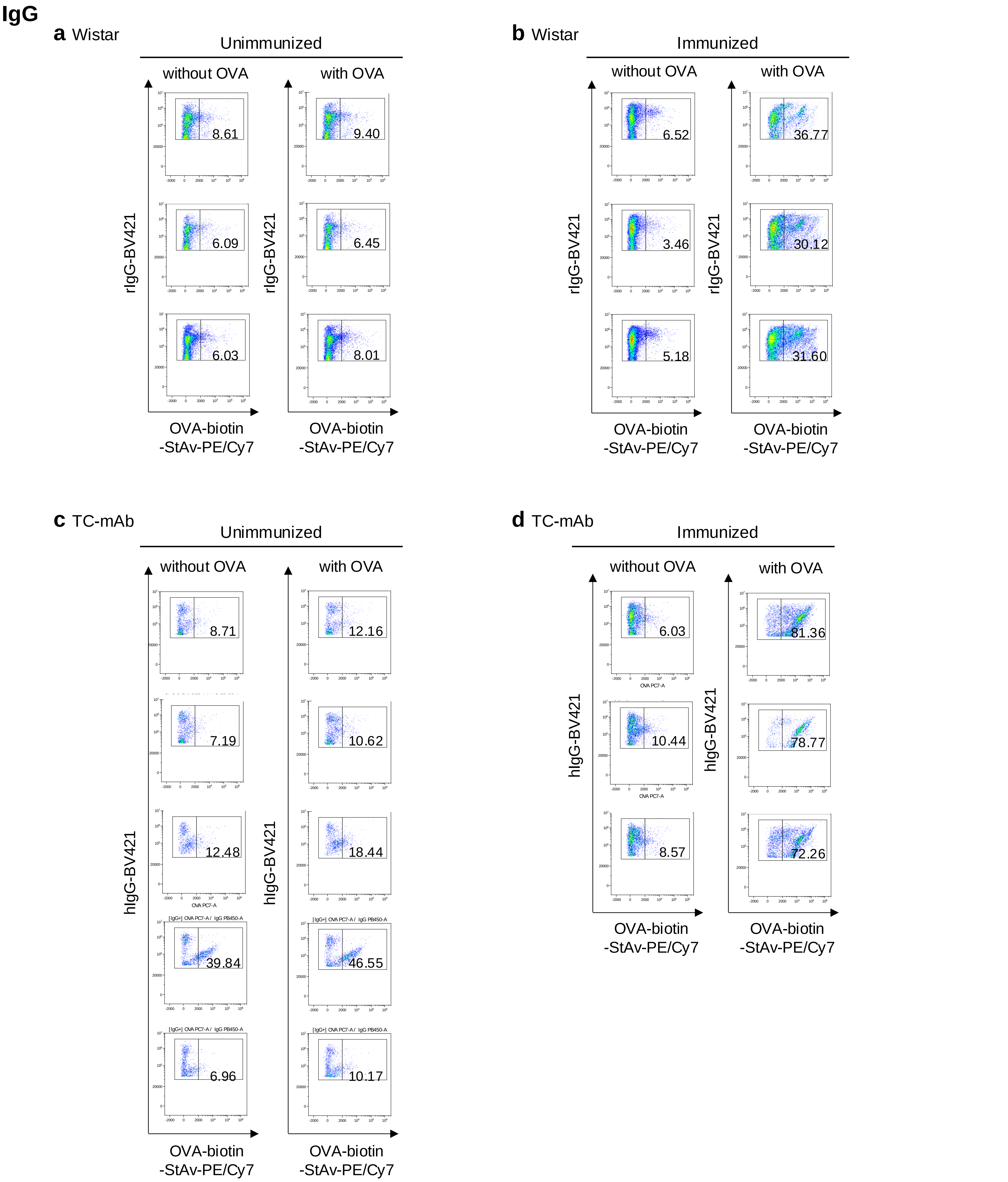

**Supplementary Figure 22| Representation of flow cytometry of antigen-specific Ab (IgG) expressing B cells in the spleen.**

The fractions of antigen-specific Ab (IgG) expressing B cells in spleen of Wistar (n=3) and TC-mAb rats (n=3). Numbers in the flow cytometry results indicate the percentage of each B cell subset. The data collected from 20-22 weeks-age Wistar (**a**: unimmunized, **b**: OVA-immunized) and TC-mAb rats (**c**: unimmunized, **d**: OVA-immunized) are indicated, respectively. OVA: Biotinylated OVA is used as bait protein.

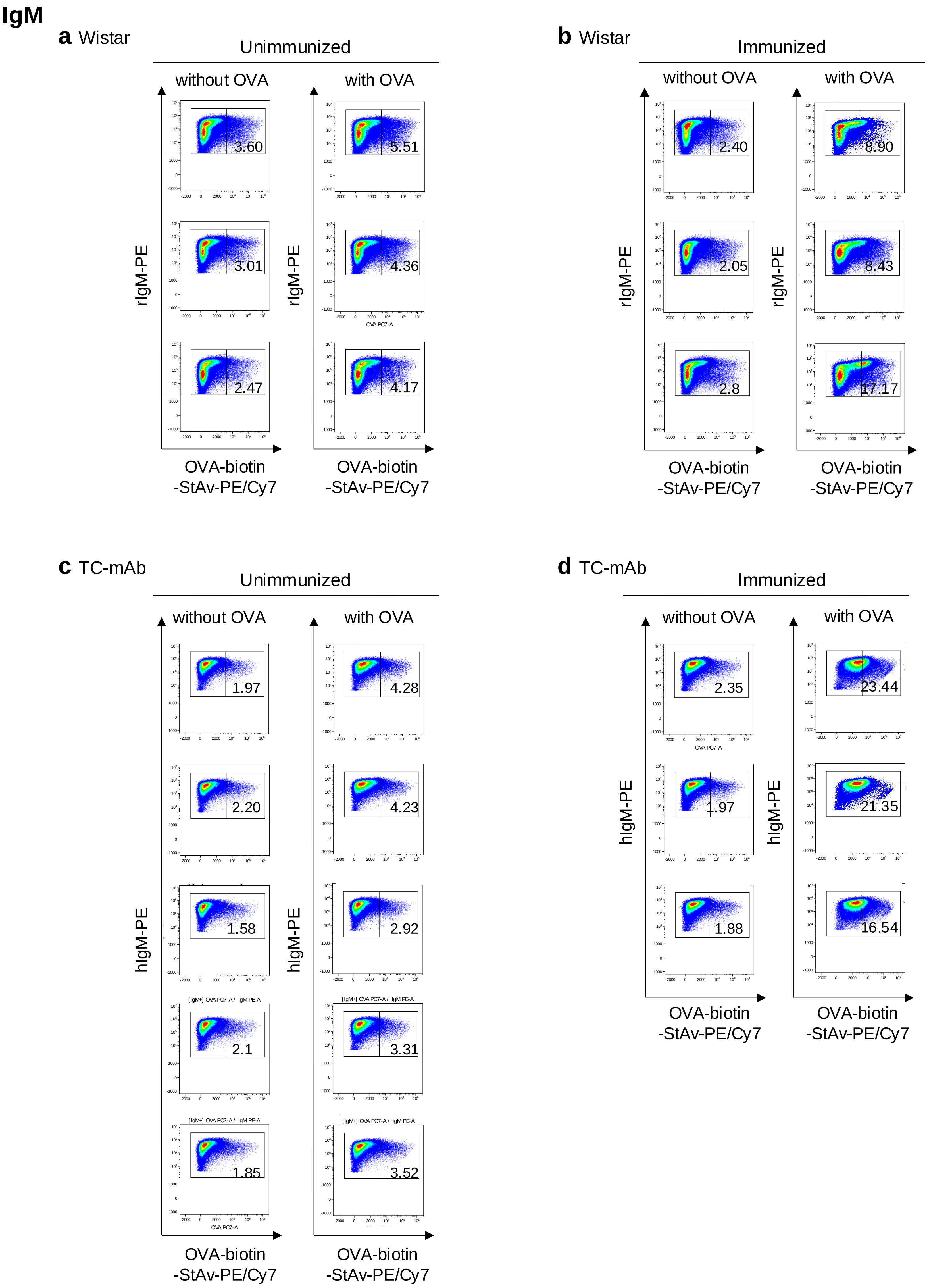

**Supplementary Figure 23| Representation of flow cytometry of antigen-specific Ab (IgM) expressing B cells in the spleen.**

The fractions of antigen-specific Ab (IgM) expressing B cells in spleen of Wistar (n=3) and TC-mAb rats (n=3). Numbers in the flow cytometry results indicate the percentage of each B cell subset. The data collected from 20-22 weeks-age Wistar (**a**: unimmunized, **b**: OVA-immunized) and TC-mAb rats (**c**: unimmunized, **d**: OVA-immunized) are indicated, respectively. OVA: Biotinylated OVA is used as bait protein.

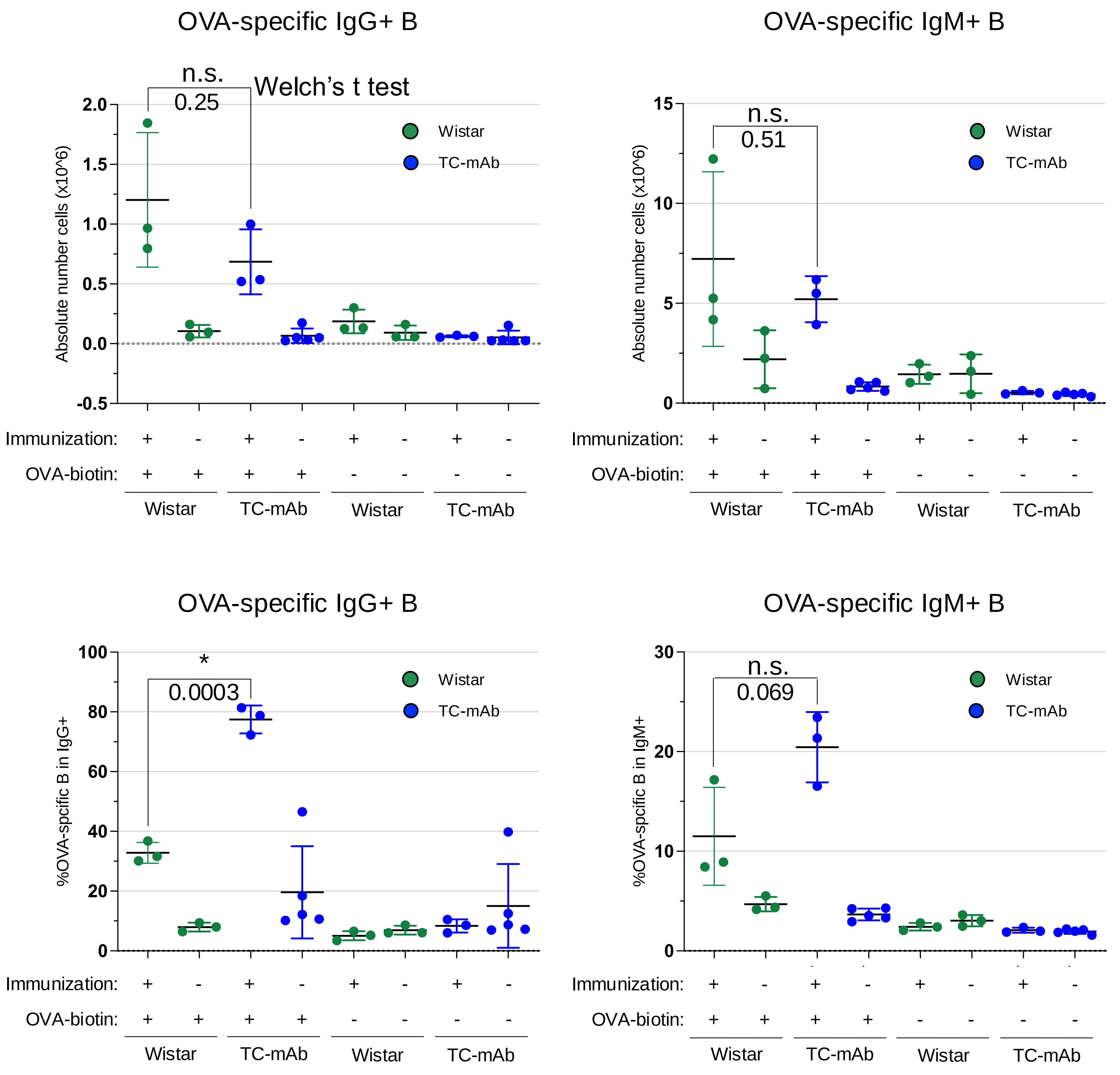

**Supplementary Figure 24| Percentage of antigen-specific Ab expressing B cells in the spleen.**
From the Flow cytometry results in Supplementary Fig. 22,23, the values detected by adding OVA (with OVA) minus the values not added (without OVA) are shown. If the value was negative, it was shown as zero. Error bars indicate the standard deviation of measurements.

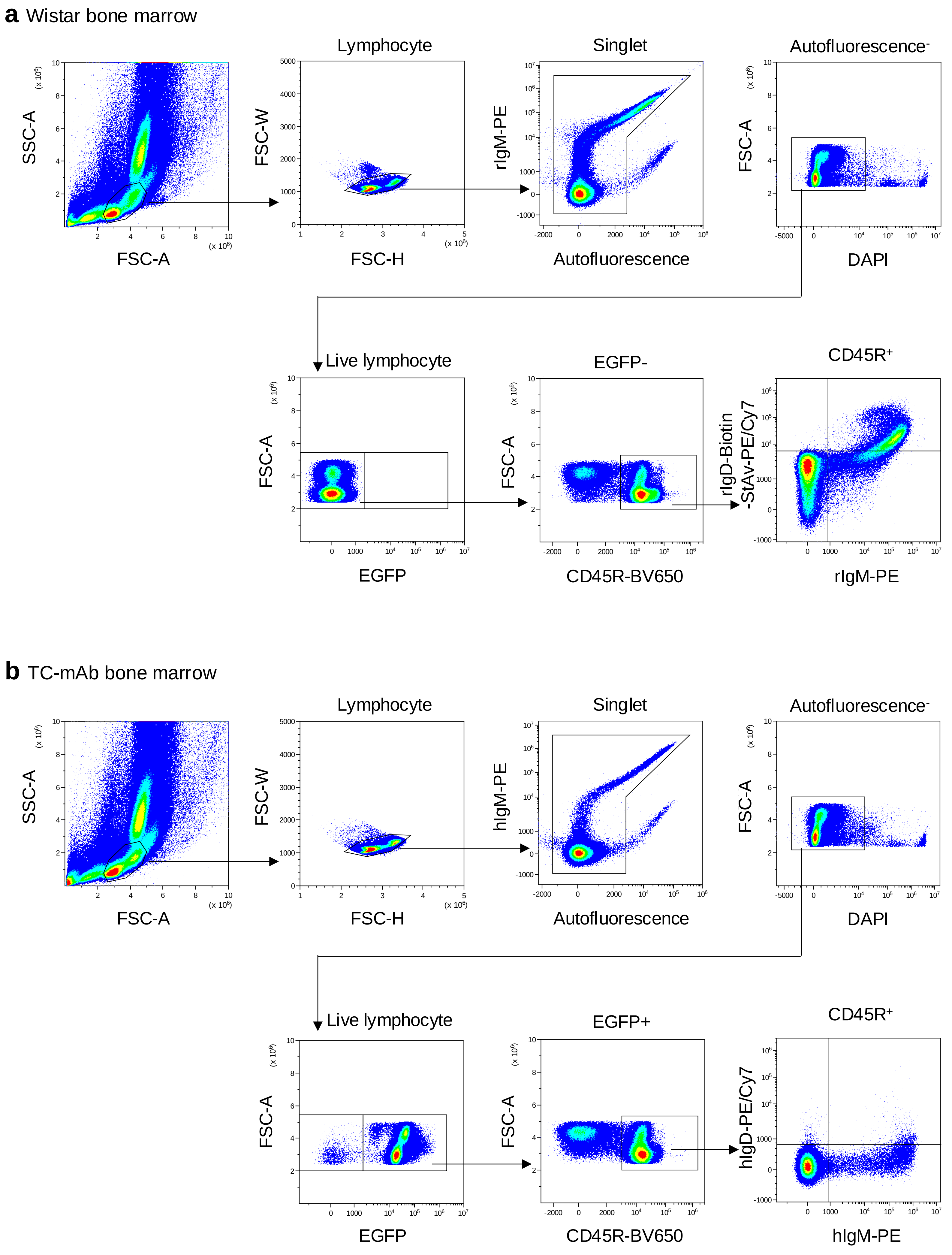

**Supplementary Figure 25| Flow cytometry identification of IgM and IgD expressing B cells in bone marrow.**
Flow cytometry gating strategies for IgM/IgD co-expressing B cells in spleen of Wistar and TC-mAb rats. The distributions were analysed using 10-12 weak-age of Wistar (**a**) and TC-mAb (**b**) rats

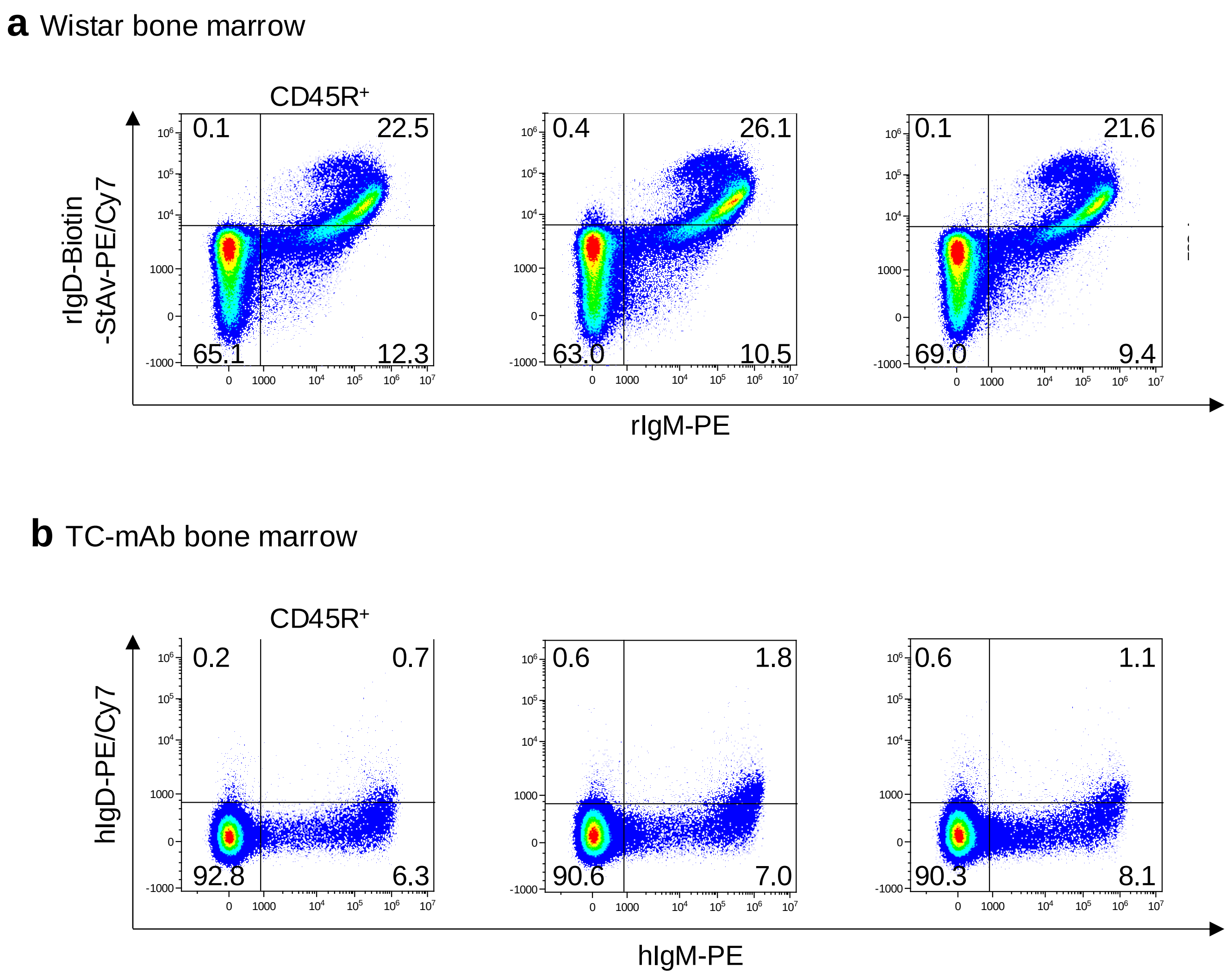

**Supplementary Figure 26| Flow cytometry identification of IgM and IgD expressing B cells in bone marrow.**
Analysis of the IgM and IgD expression of B cells in the bone marrow of Wistar (n=3) and TC-mAb rats (n=3). Numbers in the flow cytometry results indicate the percentage of each B cell subsets. The raw data are represented from Wistar **(a)** and TC-mAb rats **(b)**, respectively.

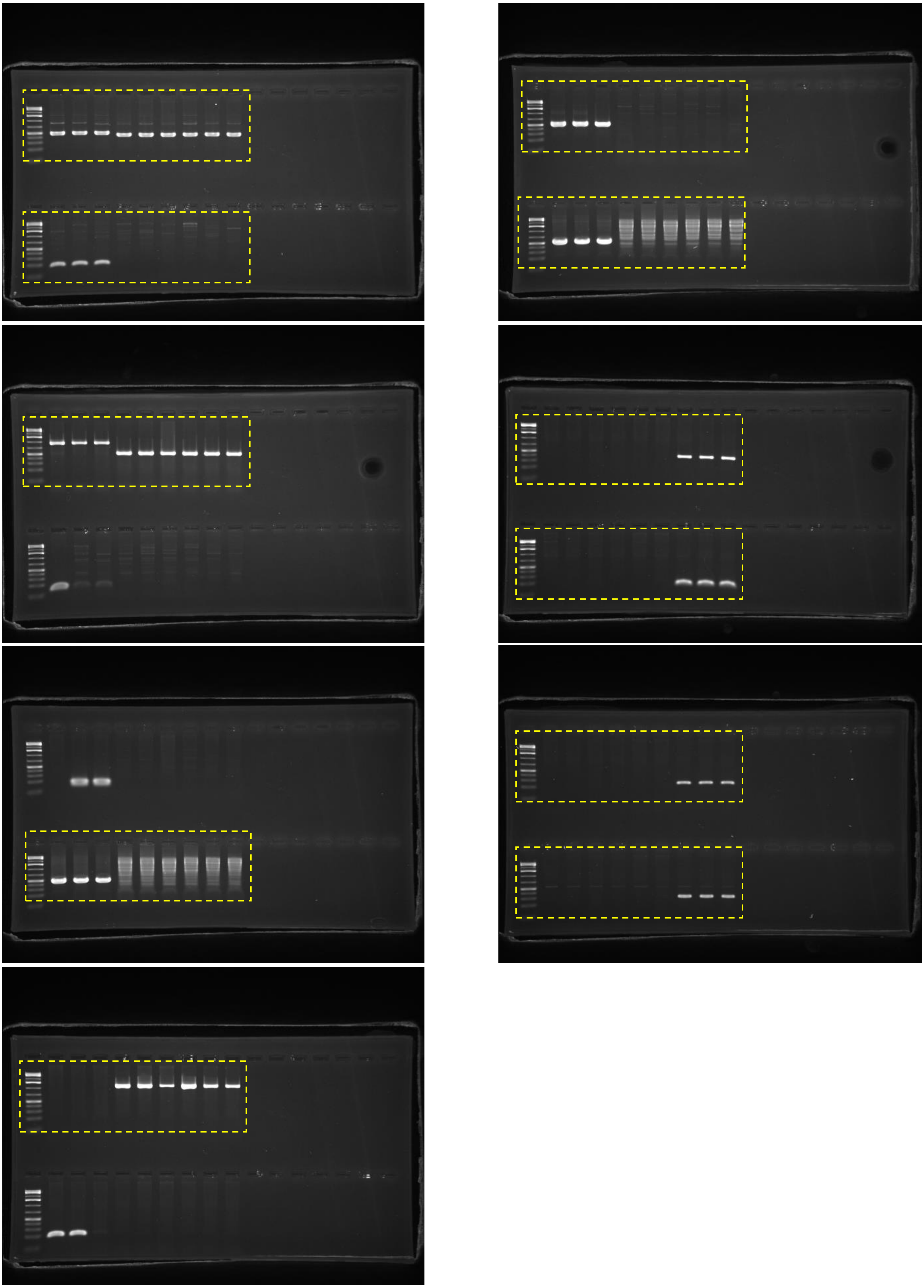

**Supplementary Figure 27| KO of endogenous Ig genes and carriage of IGHK-NAC in TC-mAb rat.**
Uncropped images of electrophoresis are presented, and the dashed squares indicate the images used in Supplementary Fig.2 .

### Supplementary Tables

**Supplementary Table 1| Antibodies for determining serum immunoglobulin concentrations.**

| | | Antibody | Labelling | Dilution | Supplier | Clone |
|---|---|---|---|---|---|---|
| **hIg μ** | Capture | Goat anti-Human IgM | - | 1:100 | Bethyl Laboratories | Goat polyclonal A80-200A |
| | Detector | Goat anti-Human IgM | HRP | 1:50000 | Bethyl Laboratories | Goat polyclonal A80-200P |
| **hIg γ** | Capture | Goat anti-human IgG-Fc | - | 1:100 | Bethyl Laboratories | Goat polyclonal A80-304A |
| | Detector | Goat anti-human IgG-Fc | HRP | 1:100000 | Bethyl Laboratories | Goat polyclonal A80-304P |
| **hIg κ** | Capture | Goat anti-Human Ig kappa | - | 1:100 | Bethyl Laboratories | Goat polyclonal A80-115A |
| | Detector | Goat anti-Human Ig kappa | HRP | 1:150000 | Bethyl Laboratories | Goat polyclonal A80-115P |
| **hIg α** | Capture | Goat anti-Human IgA | - | 1:100 | Bethyl Laboratories | Goat polyclonal A80-202A |
| | Detector | Goat anti-Human IgA | HRP | 1:75000 | Bethyl Laboratories | Goat polyclonal A80-202P |
| **hIg ε** | Capture | Goat anti-Human IgE | - | 1:100 | Bethyl Laboratories | Goat polyclonal A80-108A |
| | Detector | Goat anti-Human IgE | HRP | 1:150000 | Bethyl Laboratories | Goat polyclonal A80-108P |
| **hIg γ subclass** | Kit (991000 ) | | | | Thermo Fisher Scientific | |
| **rIg μ** | Capture | Goat anti-Rat IgM | - | 1:100 | Bethyl Laboratories | Goat polyclonal A110-200A |
| | Detector | Goat anti-Rat IgM | HRP | 1:50000 | Bethyl Laboratories | Goat polyclonal A110-200P |
| **rIg γ** | Capture | Goat anti-Rat IgG-Fc | - | 1:100 | Bethyl Laboratories | Goat polyclonal A110-236A |
| | Detector | Goat anti-Rat IgG-Fc | HRP | 1:100000 | Bethyl Laboratories | Goat polyclonal A110-236P |
| **rIg κ** | Capture | Mouse anti-Rat Ig kappa | - | 1:100 | Biolegend | MRK-81 |
| | Detector | Mouse anti-Rat Ig kappa | HRP | 1:100000 | SouthernBiotech | K4F5 |
| **rIg λ** | Capture | Mouse anti-Rat Ig lambda | - | 1:100 | Biolegend | MRL-61 |
| | Detector | Mouse anti-Rat Ig lambda | HRP | 1:25000 | Bio-Rad | MARL-15 |

| Supplementary Table 2\| Serum Ig-levels in TC-mAb rats. | | Average (μg/ml) | SD | Range |
|---|---|---|---|---|
| **Unimmunized TC-mAb rats** | | | | |
| **Number of rats=38** | | | | |
| **hIgμ** | | 33.1 | 6.4 | 23.1-48.0 |
| **hIgγ** | | 421.7 | 84.2 | 252.9-596.4 |
| **hIgκ** | | 250.0 | 37.3 | 181.8-313.5 |
| **Number of rats=38** | | | | |
| **hIgα** | | 9.6 | 3.9 | 4.9-20.1 |
| **hIgε** | | 1.6 | 0.4 | 1.2-2.8 |
| **Number of rats=18** | **Average(±SD) % in IgG** | | | |
| **hIgγ1** | 50.3 (±7.0) | 223.7 | 63.1 | 140.5-405.0 |
| **hIgγ2** | 30.7 (±8.2) | 134.4 | 40.0 | 62.8-244.2 |
| **hIgγ3** | 6.2 (±3.1) | 28.2 | 17.0 | 8.1-73.1 |
| **hIgγ4** | 12.8 (±5.1) | 55.7 | 24.6 | 19.5-128.1 |
| **Unimmunized TC-mAb rats** | | | | |
| **Number of rats= 2** | | | | |
| **6 week-age** | | X, Y | X, Y (triplicate) | |
| **hIgμ** | | 40.3, 45.9 | 1.2, 1.7 | 40.3-45.9 |
| **hIgγ** | | 144.7, 67.8 | 3.7, 7.1 | 67.8-144.7 |
| **hIgκ** | | 85.1, 58.4 | 2.9, 1.5 | 58.4-85.1 |
| **21 week-age** | | | | |
| **hIgμ** | | 96.2, 102.6 | 5.3, 4.7 | 96.2-102.6 |
| **hIgγ** | | 657.8, 667.7 | 9.4, 14.3 | 657.8-667.7 |
| **hIgκ** | | 364.5, 354.7 | 7.0, 3.4 | 354.7-364.5 |
| **OVA-immunized TC-mAb rats** | | | | |
| **Number of rats= 2** | | | | |
| **Before immunization (6 week-age)** | | A, B | A, B (triplicate) | |
| **hIgμ** | | 64.4, 34.3 | 0.1, 1.1 | 34.3-64.4 |
| **hIgγ** | | 177.6, 31.4 | 7.8, 1.3 | 31.4-177.6 |
| **hIgκ** | | 142.7, 30.7 | 9.3, 2.4 | 30.7-142.7 |
| **After immunization (19 or 23 week-age)** | | | | |
| **hIgμ** | | 375.2, 232.7 | 5.4, 3.8 | 232.7-375.2 |
| **hIgγ** | | 2276.6, 2448.8 | 77.6, 98.2 | 2276.6-2448.8 |
| **hIgκ** | | 1391.7, 1475.8 | 63.5, 61.8 | 1391.7-1475.8 |
| **Unimmunized Wistar rats** | | | | |
| **Number of rats= 1** | | | | |
| **6 week-age** | | Z | Z (triplicate) | |
| **rIgμ** | | 72.4 | 1.3 | |
| **rIgγ** | | 519.6 | 6.6 | |
| **rIgκ** | | 2538.0 | 14.1 | |
| **rIgλ** | | 11.3 | 0.3 | |
| **21 week-age** | | | | |
| **rIgμ** | | 178.3 | 4.7 | |
| **rIgγ** | | 1073.5 | 5.9 | |
| **rIgκ** | | 3222.7 | 0.0 | |
| **rIgλ** | | 31.0 | 0.2 | |
| **OVA-immunized Wistar rats** | | | | |
| **Number of rats= 2** | | | | |
| **Before immunization (6 week-age)** | | #6, 8 | #6, 8 (triplicate) | |
| **rIgμ** | | 41.6, 49.4 | 0.6, 1.0 | 41.6-49.4 |
| **rIgγ** | | 299.0, 273.2 | 2.7, 1.2 | 273.2-299.0 |
| **rIgκ** | | 2433.1, 2668.0 | 11.7, 0.0 | 2433.1-2668.0 |
| **rIgλ** | | 13.2, 12.6 | 0.4, 0.03 | 12.6-13.2 |
| **After immunization (24 week-age)** | | | | |
| **rIgμ** | | 211.2, 214.1 | 4.1, 2.5 | 211.2-214.1 |
| **rIgγ** | | 5933.1, 8343.2 | 233.3, 61.3 | 5933.1-8343.2 |
| **rIgκ** | | 4045.2, 5743.8 | 13.6, 5.5 | 4045.2-5743.8 |
| **rIgλ** | | 175.0, 249.7 | 2.2, 16.8 | 175.0-249.7 |

**Supplementary Table 3| Data collection of B cell development (Peripheral blood).**

| | | Wistar rat | HKLD rat | TC-mAb rat |
|---|---|---|---|---|
| **Rats** | **Gating conditions** | **Unimmunized (n=6)** | **Unimmunized (n=16)** | **Unimmunized (n=166)** |
| | | **%Total** | | |
| **Live lymphocyte** | FSC, SSC, Autofluorescence, DAPI | 20.5 (±18.22) | 6.0 (±8.0) | 7.5 (±7.6) |
| | | **%Live lymphocyte** | | |
| **IGHK-NAC-positive cells** | $EGFP^+$ | | - | 96.3 (±7.5) |
| | | **%Live lymphocyte** | | |
| **B cells** | $CD45R^+$ | 15.8 (±4.6) | 1.4 (±0.8) | 4.8 (±2.2) |

**Supplementary Table 4| Data collection of B cell development (Bone marrow).**

| Rats | Gating conditions | Wistar rat | | TC-mAb rat | |
|---|---|---|---|---|---|
| | | Unimmunized (n=3) | | Unimmunized (n=3) | |
| | | Absolute number | %Subset | Absolute number | %Subset |
| | | (×$10^6$) | %Lymphocyte | (×$10^6$) | %Lymphocyte |
| Total cells | | 50.5 (±29.6) | | 48.9 (±22.9) | |
| Live lymphocytes | FSC, SSC, Autofluorescence, DAPI | 10.2 (±6.3) | | 9.7 (±5.0) | |
| IGHK-NAC-positive cells | $EGFP^+$ | - | - | 9.5 (±5.1) | 95.8 (±3.5) |
| $CD45R^+$ B | $CD45R^+$ | 6.6 (±4.2) | 64.0 (±1.1) | 5.6 (±2.9) | 60.3 (±1.5) |
| | | | % $CD45R^+$ B | | % $CD45R^+$ B |
| Pro B, Pre B | $CD90^{hi}IgM^+$ | 4.3 (±3.0) | 63.2 (±4.4) | 5.0 (±2.6) | 88.0 (±2.1) |
| Developing B | $CD90^{hi}IgM^+$ | 1.8 (±1.0) | 28.9 (±2.9) | 0.4 (±0.2) | 7.1 (±0.5) |
| Mature B | $CD90^{lo}IgM^+$ | 0.15 (±0.09) | 2.1 (±0.7) | 0.05 (±0.03) | 1.0 (±0.3) |

Data represent the average ± SD of three independent experiments from different animals.

**Supplementary Table 5| Data collection of B cell development (Spleen).**

| Rats | Gating conditions | Wistar rat | | TC-mAb rat | |
|---|---|---|---|---|---|
| | | Unimmunized (n=3) | | Unimmunized (n=3) | |
| | | **Absolute number** | **%Subset** | **Absolute number** | **%Subset** |
| | | **(×$10^6$)** | **%Lymphocyte** | **(×$10^6$)** | **%Lymphocyte** |
| **Total cells** | | 314 (±89.19) | | 270.67 (±53.9) | |
| **Live lymphocytes** | FSC, SSC, Autofluorescence, DAPI | 123.5 (±63.5) | | 151.9 (±30.9) | |
| **IGHK-NAC-positive cells** | $EGFP^+$ | | | 146.0 (±32.3) | 95.8 (±1.9) |
| **$CD45R^+$ B** | $CD45R^+$ | 69.2 (±33.7) | 57.0 (±2.6) | 56.0 (±10.2) | 38.6 (±1.9) |
| | | | **% $CD45R^+$ B** | | **% $CD45R^+$ B** |
| **Developing B** | $CD90^{hi}IgM^+$ | 12.6 (±5.9) | 18.3 (±0.6) | 5.5 (±1.0) | 9.7 (±0.4) |
| **Mature B** | $CD90^{lo}IgM^+$ | 46.5 (±23.9) | 66.3 (±2.1) | 39.2 (±8.4) | 69.6 (±1.7) |

Data represent the average ± SD of three independent experiments from different animals.

**Supplementary Table 6| Data collection of NGS.**

| **Species** | **TC-mAb rats** | | | | | | **hPBMCs** | |
|---|---|---|---|---|---|---|---|---|
| **Sample collection** | Unimmunized | | Immunized | | | | Healthy donors (Asians) | |
| | *n=5* | | Individual A | | Individual B | | 426 male/female, ages: 18-54 | |
| **Analysed chain** | *IGH* (IgG and IgM) | *IGK* | *IGH* (IgG and IgM) | *IGK* | *IGH* (IgG and IgM) | *IGK* | *IGH* (IgG and IgM) | *IGK* |
| **Original reads** | 1,928,944 | 862,732 | 1,806,352 | 881,536 | 2,130,588 | 826,024 | 1,662,560 | 1,483,520 |
| **Merged reads** | 983,524 | 620,564 | 993,144 | 622,672 | 1,129,644 | 608,608 | 1,483,520 | 1,082,276 |
| **Annotated reads** | 245,531 | 155,097 | 247,969 | 155,614 | 281,868 | 152,125 | 156,798 | 270,485 |
| **CDR3 reads** | 225,923 | 149,593 | 239,991 | 148,867 | 274,787 | 147,029 | 139,745 | 262,936 |
| **Productive reads (%)** | 209,155 (92.6 %) | 141,949 (94.9 %) | 221,631 (92.3 %) | 142,064 (95.4 %) | 253,129 (92.1 %) | 139,546 (94.9 %) | 127,103 (91.0 %) | 248,872 (94.7 %) |

Merged reads represented the number of NGS reads recovered from preprocessing with merge paired-end reads of original reads. Annotated reads and CDR3 reads were determined as human Ig-sequence identified by IgBlast. Productive reads (%) indicated the percentage of productive amino acid sequence with in-frame and contained no stop codon.

### Heavy chain

**Supplementary Table 7| Frequency usage of V-gene segments.**

| | hPBMC | Unimmunized | Immunized Individual A | Immunized Individual B |
|---|---|---|---|---|
| IGHV6-1 | 0.53 | 7.40 | 9.61 | 8.80 |
| IGHV1-2 | 2.46 | 1.49 | 0.91 | 2.38 |
| IGHV1-3 | 3.00 | 2.61 | 0.90 | 0.85 |
| IGHV4-4 | 1.79 | 0.09 | 0.07 | 0.05 |
| IGHV7-4 | 2.47 | 2.74 | 8.74 | 4.51 |
| IGHV2-5 | 4.08 | 17.48 | 9.71 | 11.00 |
| IGHV3-7 | 5.42 | 1.11 | 1.18 | 1.32 |
| IGHV1-8 | 0.45 | 2.00 | 1.79 | 3.36 |
| IGHV3-9 | 5.17 | 7.20 | 8.39 | 8.66 |
| IGHV3-11 | 2.49 | 1.23 | 1.03 | 1.16 |
| IGHV3-13 | 1.20 | 1.02 | 2.09 | 1.46 |
| IGHV3-15 | 2.44 | 2.09 | 1.47 | 1.17 |
| IGHV1-18 | 5.83 | 2.81 | 4.64 | 2.32 |
| IGHV3-20 | 0.02 | 0.85 | 0.57 | 0.53 |
| IGHV3-21 | 2.36 | 1.51 | 1.16 | 4.51 |
| IGHV3-23 | 12.45 | 3.82 | 3.45 | 4.01 |
| IGHV1-24 | 0.54 | 3.56 | 3.83 | 2.28 |
| IGHV2-26 | 0.99 | 1.32 | 1.36 | 3.05 |
| IGHV4-28 | 0.21 | 0.87 | 0.54 | 0.35 |
| IGHV3-30 | 5.40 | 0.14 | 0.12 | 0.13 |
| IGHV4-31 | 1.20 | 0.02 | 0.01 | 0.02 |
| IGHV3-33 | 2.89 | 0.91 | 0.81 | 1.07 |
| IGHV4-34 | 4.27 | 10.68 | 3.37 | 5.72 |
| IGHV4-39 | 2.45 | 2.25 | 4.41 | 4.35 |
| IGHV3-43 | 0.92 | 0.09 | 0.19 | 0.08 |
| IGHV1-45 | 0.01 | 0.02 | 0.02 | 0.02 |
| IGHV1-46 | 1.01 | 0.42 | 0.23 | 0.43 |
| IGHV3-48 | 4.90 | 1.78 | 1.60 | 2.00 |
| IGHV3-49 | 0.90 | 0.38 | 0.27 | 0.37 |
| IGHV5-51 | 2.84 | 3.86 | 6.23 | 7.12 |
| IGHV3-53 | 1.40 | 0.90 | 0.68 | 0.31 |
| IGHV1-58 | 0.39 | 0.07 | 0.03 | 0.04 |
| IGHV4-59 | 5.00 | 6.28 | 9.24 | 6.88 |
| IGHV4-61 | 1.17 | 2.73 | 1.87 | 1.78 |
| IGHV3-64 | 0.73 | 0.16 | 0.05 | 0.05 |
| IGHV3-66 | 0.96 | 0.05 | 0.08 | 0.05 |
| IGHV1-69 | 4.49 | 3.30 | 2.84 | 1.96 |
| IGHV2-70 | 1.44 | 1.93 | 4.22 | 3.77 |
| IGHV3-72 | 0.75 | 0.16 | 0.18 | 0.06 |
| IGHV3-73 | 0.50 | 0.62 | 0.44 | 0.66 |
| IGHV3-74 | 2.49 | 2.02 | 1.67 | 1.37 |

**Supplementary Table 7| Frequency usage of D-gene segments.**

| | Length (bp) | hPBMC | Unimmunized | Immunized Individual A | Immunized Individual B |
|---|---|---|---|---|---|
| IGHD1-1 | 17 | 2.37 | 2.02 | 2.09 | 2.19 |
| IGHD2-2 | 31 | 9.47 | 1.26 | 1.37 | 1.41 |
| IGHD3-3 | 31 | 2.59 | 0.43 | 0.20 | 0.63 |
| IGHD4-4 | 16 | 0.00 | 0.00 | 0.00 | 0.00 |
| IGHD5-5 | 20 | 0.00 | 0.00 | 0.00 | 0.00 |
| IGHD6-6 | 18 | 2.88 | 0.46 | 0.26 | 0.12 |
| IGHD1-7 | 17 | 0.83 | 0.19 | 0.14 | 0.26 |
| IGHD2-8 | 31 | 0.93 | 0.32 | 0.27 | 1.14 |
| IGHD3-9 | 31 | 5.15 | 2.70 | 1.50 | 1.48 |
| IGHD3-10 | 31 | 11.21 | 10.78 | 13.87 | 13.53 |
| IGHD4-11 | 16 | 0.35 | 0.31 | 0.14 | 0.09 |
| IGHD5-12 | 23 | 3.62 | 2.00 | 1.77 | 3.28 |
| IGHD6-13 | 21 | 9.36 | 12.52 | 10.54 | 9.37 |
| IGHD1-14 | 17 | 1.88 | 0.55 | 0.29 | 1.09 |
| IGHD2-15 | 31 | 4.28 | 1.54 | 1.73 | 0.77 |
| IGHD3-16 | 37 | 3.63 | 2.59 | 1.90 | 1.98 |
| IGHD4-17 | 16 | 4.06 | 4.74 | 8.20 | 3.86 |
| IGHD5-18 | 20 | 2.78 | 1.71 | 1.53 | 1.59 |
| IGHD6-19 | 21 | 5.99 | 23.64 | 22.16 | 24.05 |
| IGHD1-20 | 17 | 0.35 | 1.03 | 0.71 | 1.04 |
| IGHD2-21 | 28 | 2.12 | 1.16 | 2.08 | 1.08 |
| IGHD3-22 | 31 | 6.95 | 4.90 | 1.95 | 1.87 |
| IGHD4-23 | 19 | 2.32 | 1.24 | 1.05 | 1.26 |
| IGHD5-24 | 20 | 4.41 | 0.96 | 0.30 | 0.28 |
| IGHD6-25 | 18 | 0.19 | 0.36 | 0.18 | 0.34 |
| IGHD1-26 | 20 | 7.11 | 14.72 | 13.98 | 20.01 |
| IGHD7-27 | 11 | 0.23 | 6.68 | 9.98 | 6.66 |

**Supplementary Table 7| Frequency usage of J-gene segments.**

| | hPBMC | Unimmunized | Immunized Individual A | Immunized Individual B |
|---|---|---|---|---|
| IGHJ1 | 3.17 | 4.35 | 1.20 | 1.91 |
| IGHJ2 | 5.10 | 4.87 | 6.91 | 4.48 |
| IGHJ3 | 10.76 | 15.29 | 11.81 | 20.05 |
| IGHJ4 | 52.35 | 60.22 | 62.09 | 57.34 |
| IGHJ5 | 10.47 | 4.42 | 5.46 | 5.37 |
| IGHJ6 | 18.15 | 10.85 | 12.52 | 10.84 |

**Supplementary Table 8| Data Shannon-Weaver Index (H').**

| | hPBMCs | Unimmunized | Immunized (Individual A) | Immunized (Individual B) |
|---|---|---|---|---|
| **HV** | 3.331 | 3.077 | 3.089 | 3.132 |
| **HD** | 3.018 | 2.545 | 2.483 | 2.426 |
| **HJ** | 1.386 | 1.255 | 1.205 | 1.254 |
| **KV** | 2.574 | 2.785 | 2.746 | 2.647 |
| **KJ** | 1.495 | 1.477 | 1.462 | 1.476 |

The Shannon-Weaver diversity indexes were represented based on communication theory. The indexes were calculated using productive reads indicated in Supplementary Table 1.

**Supplementary Table 9| Data collection of frequency use of DH segments.**

| | | | hPBMCs | | Unimmunized | | Immunized (individual A) | | Immunized (Individual B) | |
|---|---|---|---|---|---|---|---|---|---|---|
| | | average | frequency use (%) | bp | frequency use (%) | bp | frequency use (%) | bp | frequency use (%) | bp |
| IGHD1-1 | 17 | | 2.367 | 0.402 | 2.024 | 0.344 | 2.085 | 0.354 | 2.192 | 0.373 |
| IGHD1-7 | 17 | | 0.834 | 0.142 | 0.189 | 0.032 | 0.144 | 0.025 | 0.259 | 0.044 |
| IGHD1-14 | 17 | 17.6 | 1.881 | 0.320 | 0.551 | 0.094 | 0.293 | 0.050 | 1.090 | 0.185 |
| IGHD1-20 | 17 | | 0.352 | 0.060 | 1.032 | 0.175 | 0.712 | 0.121 | 1.043 | 0.177 |
| IGHD1-26 | 20 | | 7.114 | 1.423 | 14.717 | 2.943 | 13.978 | 2.796 | 20.008 | 4.002 |
| IGHD2-2 | 31 | | 9.474 | 2.937 | 1.258 | 0.390 | 1.374 | 0.426 | 1.410 | 0.437 |
| IGHD2-8 | 31 | 30.25 | 0.925 | 0.287 | 0.324 | 0.100 | 0.266 | 0.083 | 1.142 | 0.354 |
| IGHD2-15 | 31 | | 4.283 | 1.328 | 1.544 | 0.479 | 1.733 | 0.537 | 0.767 | 0.238 |
| IGHD2-21 | 28 | | 2.120 | 0.594 | 1.157 | 0.324 | 2.076 | 0.581 | 1.079 | 0.302 |
| IGHD3-3 | 31 | | 2.592 | 0.804 | 0.434 | 0.134 | 0.196 | 0.061 | 0.626 | 0.194 |
| IGHD3-9 | 31 | | 5.148 | 1.596 | 2.700 | 0.837 | 1.498 | 0.464 | 1.477 | 0.458 |
| IGHD3-10 | 31 | 32.2 | 11.215 | 3.477 | 10.783 | 3.343 | 13.869 | 4.299 | 13.527 | 4.193 |
| IGHD3-16 | 37 | | 3.634 | 1.344 | 2.589 | 0.958 | 1.897 | 0.702 | 1.977 | 0.731 |
| IGHD3-22 | 31 | | 6.946 | 2.153 | 4.900 | 1.519 | 1.950 | 0.604 | 1.872 | 0.580 |
| IGHD4-4 | 16 | | 0.000 | 0.000 | 0.000 | 0.000 | 0.000 | 0.000 | 0.000 | 0.000 |
| IGHD4-11 | 16 | 16.75 | 0.352 | 0.056 | 0.309 | 0.049 | 0.136 | 0.022 | 0.094 | 0.015 |
| IGHD4-23 | 19 | | 2.321 | 0.441 | 1.238 | 0.235 | 1.048 | 0.199 | 1.258 | 0.239 |
| IGHD4-17 | 16 | | 4.059 | 0.649 | 4.744 | 0.759 | 8.203 | 1.312 | 3.864 | 0.618 |
| IGHD5-5 | 20 | | 0.000 | 0.000 | 0.000 | 0.000 | 0.000 | 0.000 | 0.000 | 0.000 |
| IGHD5-12 | 23 | 20.75 | 3.624 | 0.834 | 2.003 | 0.461 | 1.767 | 0.406 | 3.278 | 0.754 |
| IGHD5-24 | 20 | | 4.414 | 0.883 | 0.964 | 0.193 | 0.304 | 0.061 | 0.276 | 0.055 |
| IGHD5-18 | 20 | | 2.781 | 0.556 | 1.709 | 0.342 | 1.528 | 0.306 | 1.588 | 0.318 |
| IGHD6-6 | 18 | | 2.883 | 0.519 | 0.457 | 0.082 | 0.264 | 0.048 | 0.117 | 0.021 |
| IGHD6-13 | 21 | 19.5 | 9.361 | 1.966 | 12.517 | 2.629 | 10.537 | 2.213 | 9.371 | 1.968 |
| IGHD6-19 | 21 | | 5.992 | 1.258 | 23.635 | 4.963 | 22.164 | 4.654 | 24.048 | 5.050 |
| IGHD6-25 | 18 | | 0.190 | 0.034 | 0.357 | 0.064 | 0.177 | 0.032 | 0.339 | 0.061 |
| IGHD7-27 | 11 | 11 | 0.002 | 0.025 | 6.677 | 0.734 | 9.982 | 1.098 | 6.664 | 0.733 |
| Average | | | | 21.741 | | 18.596 | | 18.108 | | 17.320 |

**Supplementary Table 10| Data collection of frequency use of P-N addition**

| | hPBMCs | Unimmunized | Immunized (individual A) | Immunized (Individual B) |
|---|---|---|---|---|
| V-D | 7.68 | 3.30 | 3.53 | 3.22 |
| D-J | 6.96 | 2.48 | 2.59 | 2.80 |
| Total average of the P-N addition | 14.64 | 5.78 | 6.12 | 6.02 |

### Kappa chain

**Supplementary Table 11| Frequency usage of V-gene segments.**

| | | hPBMC | Unimmunized | Immunized Individual A | Immunized Individual B |
|---|---|---|---|---|---|
| Proximal cluster | IGKV4-1 | 12.68 | 14.34 | 18.85 | 17.19 |
| | IGKV5-2 | 0.60 | 1.09 | 0.90 | 2.29 |
| | IGKV1-5 | 11.92 | 5.04 | 4.10 | 3.99 |
| | IGKV1-6 | 1.12 | 1.15 | 1.78 | 0.88 |
| | IGKV1-9 | 2.93 | 2.48 | 1.89 | 2.88 |
| | IGKV3-11 | 6.54 | 2.82 | 1.80 | 1.83 |
| | IGKV1-12 | 3.59 | 3.19 | 3.61 | 2.92 |
| | IGKV3-15 | 8.71 | 6.25 | 5.87 | 5.70 |
| | IGKV1-16 | 3.34 | 4.88 | 4.00 | 3.19 |
| | IGKV1-17 | 1.59 | 3.41 | 2.27 | 2.80 |
| | IGKV3-20 | 18.47 | 15.52 | 14.54 | 17.88 |
| | IGKV2-24 | 1.53 | 6.65 | 5.47 | 3.52 |
| | IGKV1-27 | 2.37 | 4.76 | 5.13 | 4.99 |
| | IGKV2-28 | 1.73 | 1.74 | 4.19 | 2.10 |
| | IGKV2-30 | 3.10 | 4.70 | 3.55 | 3.99 |
| | IGKV1-33 | 4.06 | 8.55 | 6.89 | 5.28 |
| | IGKV1-39 | 13.05 | 7.89 | 9.21 | 14.38 |
| | IGKV2-40 | 0.06 | 0.27 | 0.56 | 0.30 |
| | 800 kbp | | | | |
| Distal cluster | IGKV2D-40 | 0.00 | 0.00 | 0.00 | 0.00 |
| | IGKV1D-39 | 0.00 | 0.00 | 0.00 | 0.00 |
| | IGKV1D-33 | 0.00 | 0.00 | 0.01 | 0.00 |
| | IGKV2D-30 | 0.00 | 0.49 | 0.82 | 0.32 |
| | IGKV2D-29 | 0.12 | 0.87 | 1.33 | 1.18 |
| | IGKV2D-28 | 0.00 | 0.00 | 0.00 | 0.00 |
| | IGKV2D-24 | 0.05 | 0.10 | 0.07 | 0.05 |
| | IGKV3D-20 | 0.19 | 0.78 | 0.49 | 0.48 |
| | IGKV1D-17 | 0.00 | 0.01 | 0.00 | 0.00 |
| | IGKV1D-16 | 0.36 | 0.74 | 0.58 | 0.54 |
| | IGKV3D-15 | 0.92 | 0.77 | 0.95 | 0.69 |
| | IGKV1D-12 | 0.67 | 0.67 | 0.39 | 0.39 |
| | IGKV3D-11 | 0.04 | 0.02 | 0.01 | 0.01 |
| | IGKV1D-43 | 0.14 | 0.01 | 0.00 | 0.00 |
| | IGKV1D-8 | 0.12 | 0.80 | 0.74 | 0.22 |
| | IGKV3D-7 | 0.00 | 0.00 | 0.00 | 0.00 |

**Supplementary Table 12| Frequency usage of J-gene segments.**

| | hPBMC | Unimmunized | Immunized Individual A | Immunized Individual B |
|---|---|---|---|---|
| IGKJ1 | 29.07 | 28.66 | 28.64 | 26.88 |
| IGKJ2 | 7.57 | 5.91 | 6.16 | 4.27 |
| IGKJ3 | 22.47 | 24.18 | 25.11 | 27.74 |
| IGKJ4 | 29.79 | 29.88 | 30.67 | 26.60 |
| IGKJ5 | 11.12 | 11.37 | 9.42 | 14.51 |

The relative usage of human variable regions in hPBMCs and TC-mAb mice was indicated. The genes segments annotated as an open reading frame or pseudogene[18,19] were excluded.

**Supplementary Table 13| The expected value of the 50 most frequently used clone lineages.**

| | IGHV | | IGKV | |
|---|---|---|---|---|
| | Total reads | Expected value | Total reads | Expected value |
| **hPBMCs** | 15,558 | 4.38 | 136,101 | 8.82 |
| **TC-mAb rats** | | | | |
| **Unimmunized pooled (n=5)** | 136,101 | 2.21 | 80,398 | 2.23 |
| **Immunized individual A** | 57,226 | 3.68 | 79,259 | 3.75 |
| **Immunized individual B** | 88,094 | 3.70 | 78,723 | 3.89 |
| **TC-mAb mice** | | | | |
| **Unimmunized pool (n=5)** | 14,313 | 0.78 | 2,160 | 1.15 |
| **Immunized (n=1)** | 82,763 | 0.90 | 16,330 | 1.47 |

**Supplementary Table 14| Monoclonal antibody production against OVA.**

| Steps | TC-mAb rats, Individual A |
|---|---|
| **Immunization** | Primary and 5 boosters |
| **Lymphocytes** | $3.3 \times 10^8$ cells/rats |
| **Lymphocytes/cell fusion** | $1.0 \times 10^8$ cells |
| **ELISA positive well after HAT** | 285 well |
| **ELISA positive well after 2nd screening** | 80 clones |
| **Determination of subclass** | 31 clones |
| **Cloned by limiting dilution** | 30/30 clones |
| **(Success rate of cloning)** | 100% |
| **Subclass of the obtained 30 mAbs** | |
| **IgMκ** | 2 (6.7%) |
| **IgG1κ** | 27 (90%) |
| **IgG2κ** | 1 (3.3%) |
| **Others (Signals are found in both IgG1κ and IgMκ)** | 1 (3.3%) |

**Supplementary Table 15| Data collection of Antigen-specific cell..**

| Rats | Gating conditions | Wistar rat | | | |
|---|---|---|---|---|---|
| | | Unimmunized (n=3) | | Immunized (n=3) | |
| | | Absolute number | %Subset | Absolute number | %Subset |
| **Spleen** | | **(×$10^6$)** | | **(×$10^6$)** | |
| **Total cells** | | 265.7 (±54.2) | | 393.7 (±43.7) | |
| **Lymphocytes** | FSC, SSC, Autofluorescence, DAPI | 117.8 (±35.1) | | 167.8 (±33.0) | |
| | | | **%Lymphocyte** | | **%Lymphocyte** |
| **$CD45R^+$ B** | $CD45R^+$ | 51.8 (±22.4) | 41.6 (±8.1) | 68.0 (±11.7) | 40.7 (±2.4) |
| | | | **% $CD45R^+$ B** | | **% $CD45R^+$ B** |
| **$IgM^+$ B** | $CD45R^+IgM^+$ | 45.0 (±20.2) | 85.9 (±2.4) | 60.1 (±10.1) | 88.5 (±0.6) |
| **$IgG^+$ B** | $CD45R^+IgG^+$ | 1.3 (±0.4) | 3.1 (±1.4) | 3.8 (±1.6) | 5.3 (±1.4) |
| **Antigen-specific $IgM^+$ B** | $CD45R^+IgM^+OVA^+$ | 2.2 (±1.2) | 4.7 (±0.6) | 7.2 (±3.6) | 11.5 (±4.0) |
| **Antigen-specific $IgG^+$ B** | $CD45R^+IgG^+OVA^+$ | 0.1 (±0.04) | 8.0 (±1.2) | 1.2 (±0.5) | 32.8 (±2.9) |

| Rats | Gating conditions | TC-mAb rat | | | |
|---|---|---|---|---|---|
| | | Unimmunized (n=5) | | Immunized (n=3) | |
| | | Absolute number | %Subset | Absolute number | %Subset |
| **Spleen** | | **(×$10^6$)** | | **(×$10^6$)** | |
| **Total cells** | | 257.0 (±32.9) | | 237.0 (±25.4) | |
| **Lymphocytes** | FSC, SSC, Autofluorescence, DAPI | 115.3 (±7.8) | | 108.1 (±11.2) | |
| | $CD45R^+$ | | **%Lymphocyte** | | **%Lymphocyte** |
| **$CD45R^+$ B** | $CD45R^+$ | 25.7 (±2.6) | 24.8 (±0.8) | 29.3 (±1.9) | 29.4 (±1.1) |
| | | | **% $CD45R^+$ B** | | **% $CD45R^+$ B** |
| **$IgM^+$ B** | $CD45R^+IgM^+$ | 22.4 (±1.9) | 87.2 (±2.5) | 25.3 (±1.1) | 86.4 (±2.1) |
| **$IgG^+$ B** | $CD45R^+IgG^+$ | 0.3 (±0.1) | 1.3 (±0.2) | 0.9 (±0.3) | 3.0 (±0.7) |
| **Antigen-specific $IgM^+$ B** | $CD45R^+IgM^+OVA^+$ | 0.8 (±0.2) | 3.7 (±0.5) | 5.2 (±0.9) | 20.4 (±2.9) |
| **Antigen-specific $IgG^+$ B** | $CD45R^+IgG^+OVA^+$ | 0.1 (±0.1) | 19.7 (±13.8) | 0.7 (±0.2) | 77.5 (±3.8) |

Data represent the average ± SD of three or five independent experiments from different animals.

**Supplementary Table 16| Data collection of B cell development (IgM and IgD).**

| Rats | | Gating conditions | Wistar rat Unimmunized (n=3) | | TC-mAb rat Unimmunized (n=3) | |
|---|---|---|---|---|---|---|
| | | | Absolute number | %Subset | Absolute number | %Subset |
| **Bone marrow** | | | **(×$10^6$)** | **%Lymphocyte** | **(×$10^6$)** | **%Lymphocyte** |
| **Total cells** | | | 50.5 (±29.6) | | 48.9 (±22.9) | |
| **Lymphocytes** | | FSC, SSC, Autofluorescence, DAPI | 14.5 (±7.0) | | 14.5 (±7.1) | |
| **NAC-positive cells** | $EGFP^+$ | | - | | 14.1 (±7.2) | 96.4 (±3.2) |
| **$CD45R^+$ B** | | $CD45R^+$ | 11.3 (±5.1) | 79.0 (±2.5) | 10.7 (±5.3) | 76.5 (±2.1) |
| | | | | **% $CD45R^+$ B** | | **% $CD45R^+$ B** |
| **$IgM^+IgD^+$** | | $IgM^+IgD^+$ | 2.6 (±1.1) | 23.4 (±2.0) | 0.11 (±0.04) | 1.2 (±0.5) |
| **$IgM^+IgD^-$** | | $IgM^+IgD^-$ | 1.2 (±0.4) | 10.7 (±1.2) | 0.7 (±0.3) | 7.2 (±0.8) |
| **$IgM^-IgD^+$** | | $IgM^-IgD^+$ | 0.02 (±0.01) | 0.2 (±0.1) | 0.04 (±0.01) | 0.5 (±0.2) |
| **$IgM^-IgD^-$** | | $IgM^-IgD^-$ | 7.5 (±3.7) | 65.7 (±2.5) | 9.8 (±5.0) | 91.2 (±1.1) |

Data represent the average ± SD of three independent experiments from different animals.

**Supplementary Table 17| Antibody reagents used in experiments.**

| Antibody | Target species | fluorochrome | Dilution | Supplier | Clone |
|---|---|---|---|---|---|
| **CD45R** | Rat | BV650 | 1:100 | BD Biosciences | HIS24 |
| | Isotype control | BV650 | 1:100 | BD Biosciences | 27-35 |
| **CD90** | Rat | PE/Cy7 | 1:100 | Biolegend | OX-7 |
| | Isotype control | PE/Cy7 | 1:100 | BD Biosciences | MOPC-21 |
| **IgM** | Human | PE | 1:20 | Biolegend | MHM-88 |
| | Rat | PE | 1:100 | BD Biosciences | G53-238 |
| **IgD** | Human | PE/Cy7 | 1:100 | Biolegend | IA6-2 |
| | Rat | Biotin | 1:100 | Bio-rad | MARD-3 |
| **IgG** | Human | BV421 | 1:50 | Biolegend | M1310G05 |
| | Rat | BV421 | 1:100 | Jackson ImmunoResearch | Goat polyclonal 112-675-071 |
| **CD32 (FcR blocker)** | Rat | - | 1:50 | BD Biosciences | D34-485 |

**Supplementary Table 18| Primer list.**

| | Gene or aim | Primer name (forward) | Forward primer (5′-3′) | Primer name (reverse) | Reverse primer (5′-3′) | Product size |
|---|---|---|---|---|---|---|
| **Genomic PCR** | | | | | | |
| | Rat Igh | IGHMKO Cel1 Fw | TGCTGTGGCTCTGTCCCATA | IGHMKO Cel1 Rv | CCTCGGGAAGGGTTGGTTCT | 505 bp |
| | Rat Igh | TTD1 wt Fw | AGCAGCAACATGGAGACAGCAG | TTD1 wt Rv | CAAATTCTCATCAGACAGGGGG | 175 bp |
| | Rat Igκ | IGKKO Cel1 Fw2 | TGGTCTGGTATCTCTGTCTGATGCATGG | IGKKO Cel1 Rv2 | GGGCAAGGGGGAGAGTTTTATTTGTTGT | 979 bp |
| | Rat Igκ | IGKKO Cel1 Fw1 | CCATCCTCTGTGCTTCCTTCC | IGKKO Cel1 Rv1 | GCATGATCAAAGCCAAGGAAA | 503 bp |
| | Rat Igλ | rIGLCBDJ Fw | ATTTGCAGACCAAAGGGAAGGAAAGAT | rIGLCBDJ Rv | TGGTGTGATCAGAGGTCCAGAAGAAAGT | 1180 bp |
| | Rat Igλ | IGL up Cel1 Fw | CAAAGGCAACTGAAATGATGTCTTG | IGL up Cel1 Rv | TGGCCAGCTGATTCCACTCTT | 516/516 bp |
| | Rat Igλ | IGL down Cel1 Fw | TCACCCTGTTCCCACCTTCC | IGL down Cel1 Rv | GGCACTCCCTGGGGTAATGA | 550/552 bp |
| | IGKC | IGKC-F | TGGAAGGTGGATAACGCCCT | IGKC-R | TCATTCTCCTCCAACATTAGCA | 377 bp |
| | IGKV | IGKV-F | AGTCAGGGCATTAGCAGTGC | IGKV-R | GCTGCTGATGGTGAGAGTGA | 156 bp |
| | IGHV3-74 | VH3-F | AGTGAGATAAGCAGTGGATG | VH3-R | CTTGTGCTACTCCCATCACT | 247 bp |
| | IGHM | CH3F3 | AGGCCAGCATCTGCGAGGAT | CH4R2 | GTGGCAGCAAGTAGACATCG | 326 bp |
